# Versioning Biological Cells for Trustworthy Cell Engineering

**DOI:** 10.1101/2021.04.23.441106

**Authors:** Jonathan Tellechea-Luzardo, Leanne Hobbs, Elena Velázquez, Lenka Pelechova, Simon Woods, Victor de Lorenzo, Natalio Krasnogor

## Abstract

*“Full-stack”* biotechnology platforms for cell line (re)programming are on the horizon, due mostly to (a) advances in gene synthesis and editing techniques as well as (b) the growing integration with informatics, the internet of things and automation. These emerging platforms will accelerate the production and consumption of biological products. Hence, transparency, traceability and -ultimately-trustworthiness is required -from cradle to grave- for engineered cell lines and their engineering processes. We report here the first version control system for cell engineering that integrates a new cloud-based version control software for cell lines’ digital footprint with molecular barcoding of living samples. We argue that version control for cell engineering marks a significant step towards more open, reproducible, easier to trace and share, and more trustworthy engineering biology.

**One Sentence Summary:** We demonstrate a transparent and open way of engineering and sharing cell lines.

## Main Text

Engineering biology is exploding with advances ranging from new genome editing tools (1), to genetically encodable materials for advanced sensing of cells physiological states, electrical fields and mechanical stresses (2, 3), programmable and functional microbial-based living materials (4), environmental remediation and pollution control (5) to advanced *in vivo* data storage (6). Moreover, these advances in fundamental science are rapidly translating into new companies (7) and consumer products (8), which within the first half of this century, are set to impact most areas of our lives.

Perhaps the most convincing example of the pace of progress is the global scientific response to the current SARS-CoV-2 pandemic. In a matter of weeks after detecting the outbreak, the virus was isolated, its genome sequenced and published (9) and made available for research. Slightly less than a year later multiple vaccines were already being deployed to combat the virus. This would have been unimaginable just 10 years ago. Although this is an extreme example arising out of an extreme situation, it is to be expected that with the commoditization of synthetic DNA and the wider availability of powerful gene editing tools (10), the number of engineered strains will rapidly increase. Indeed, cheaper DNA synthesis technology and the development of high throughput, automated, cloning processes allows the creation of large plasmid and combinatorial dna libraries (11, 12) in a matter of days, including the modification of recalcitrant species’ genomes (13), which previously were difficult to edit.

**Version Control:** the practice of tracking changes in the source code of computer programs, which is assisted by the use of special software that keeps track of the code, its changes over time, the changes’ authors and other metadata.

And yet, while engineering biology has changed profoundly in the last few years, there are still deep gaps in the way the *process of strain engineering* is done and disseminated. For example, engineering a “synthetic biology agent” (14) produces large quantities of information: published articles, protocols, notebooks, models, databases, sequencing and other types of data (e.g. metabolomics, proteomics, lipidomics, etc). Combined, all these information sources, may add-up to terabytes of data but only a relative small percentage of it is being made available when results are published in specialized outlets. This gap in scientific practice has led to an on-going crisis in cell line misidentification (15,16,17), a recognized lack of reproducibility (18), sometimes causing high profile retractions (19) and often resulting in weakening public attitudes to new and emerging technologies. Public attitudes towards science in general, and in particular newly emerging technologies such as engineering biology, have been the focus of study over the last few decades. Some of the earlier studies were based on the so called ‘deficit model’ that assumes that publics’ support of science and novel technologies is built on their knowledge of science. Although this model is still used by some scientists, surveys and polls have shown that more knowledge of science (or technology) does not necessarily lead to more public support (20, 21). It was also shown that deference to science (22) had a large impact on perceptions of, e.g., nanotechnology, and that public trust in the claims made by experts or by scientific institutions (23, 24) is an important factor in understanding publics’ attitudes to science.

Similarly, although general support was found for the then emerging field of synthetic biology (25), it was tempered by fears over misuse, health and environmental impacts, control and governance. Taken together these findings suggest that publics’ concerns in relation to engineering biology lies more with the *method and processes* underpinning the research rather than in one or another specific technical advance. That is, transparency and accountability of the research are the main concern in publics’ attitudes to engineering biology (25,26,27).

Moreover, the potential for public mistrust in innovative science is echoed in the science community’s own crisis of conscience about the integrity of science itself (28). Concerns about the public trust in science have been raised by scientists as one factor amongst others contributing to the global critical concern about science culture and integrity (29). As it was argued (30), transparency, reproducibility and openness are the norms of ‘good’ science but are often in conflict with the competing forces of competition and the ‘success’ culture promoted within Universities. Similar concerns have been raised by other studies, UK Research and Innovation/ Research England’s survey on research integrity (31) confirmed that open and transparent research are regarded as central to rigour, reproducibility, and public trust.

Commentators on the ‘crisis’ broadly agree that the research community is overwhelmingly motivated by these values but that other factors within the research culture can make it difficult to uphold these principles. The Wellcome Trust reveals (32) that the quality and reproducibility of science is under threat as a consequence of the pressure to produce outputs and impact. In response there have been moves to promote measures that enable transparency, reproducibility and openness. The ‘FAIR Principles’ (33), namely findable, accessible, interoperable and re-usable, have been widely adopted. The UKRI Research Concordat on Open Research Data (34) has been adopted by multiple-stakeholders within the UK research community.

It is thus clear that the crisis within science culture requires a response on multiple fronts, to which this paper contributes in a number of practical ways with the introduction of CellRepo.

CellRepo is a version control system for cell engineering that integrates a new cloud-based version control software for cell lines’ digital footprint tracking and molecular barcoding of living samples. CellRepo, which substantially improves our earlier work (35), relies on two fundamental pillars. Firstly, during the process of engineering a new strain, changes introduced to the cell line are recorded in “commits” that include information such as the genotype, phenotype, author of the modifications, laboratory protocols, characterization profiles, etc. The history of commits tracks the digital footprint produced during the process of cell engineering. The second pillar is the physical linking of a living sample to a commit via the chromosomal introduction of a unique barcode related to the commit (Fig. 1). Unique barcodes have been recently recommended as the way forward for standardizing (14) synthetic biology agents and for aiding in microbial characterization and environmental risk assessment (36). To demonstrate the wide applicability of CellRepo, we show how the new platform -in conjunction with well-established peer-reviewed protocols to genetically engineer various organisms- can be applied to six of the most important and diverse microbial species used in both academia and industry (*Escherichia coli*, *Bacillus subtilis*, *Streptomyces albidoflavus*, *Pseudomonas putida*, *Saccharomyces cerevisiae* and *Komagataella phaffii* – previously known as *Pichia pastoris*). It is expected that the same principles can be applied to more, if not all, species which have been already domesticated and engineered.

**Fig 1.**
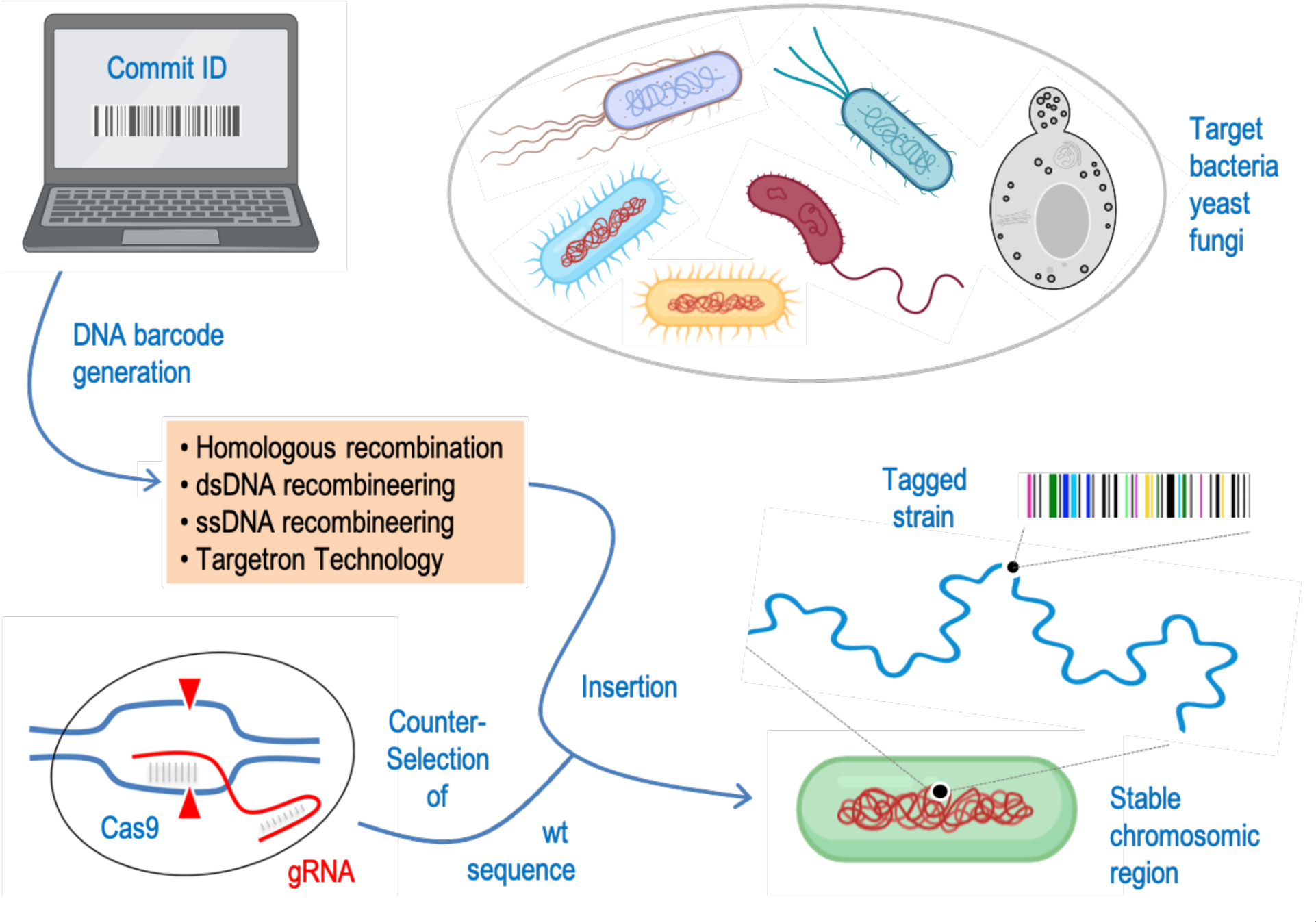
Workflow for barcoding any microorganism or strain of interest. The roadmap can be applied to basically any biological system amenable to genomic insertions of short DNA sequences. Once a specific barcode is committed, it is synthesized and delivered to a stable region of the genome of the target organism. The different approaches followed in this study to select appropriate barcoding locations are described in the Supplementary Material. This can be made with a whole collection of genetic tools available to this end in a fashion dependent or independent on homologous recombination. As the resulting insertion is expected not to generate a conspicuous phenotype, proper delivery of the barcode to the expected genomic site is secured and selected with CRISPR/Cas9 technology or via the more traditional antibiotic resistance or auxotrophic marker approach depending on the laboratory carrying out the barcoding.

## CellRepo: a cloud version control for engineering biology

As in any cloud-based application, the user needs to register, providing a name that will be public, e-mail address and password. An avatar picture can be uploaded to personalize the experience and to be more recognizable by other users. After registering, the user will receive a confirmation by email. Finally, the user can sign-in by typing the registration e-mail and password. Once a user signs in, they land in the homepage (Fig. 2a) which contains everything needed to build repositories of engineered strains, manage their accounts and the teams they work with. From the initial page it is also possible to access the system documentation (“Knowledge Base”). The blue upper horizontal quick menu links to all the aforementioned features and is present in every page in the website. This menu also contains a search-bar. This allows users to look for repositories and commits accessible to them (i.e. owned by them, in one of their team or repositories that are public) and look for identifiers to find the documentation on specific strains/plasmids.

**Fig. 2.**
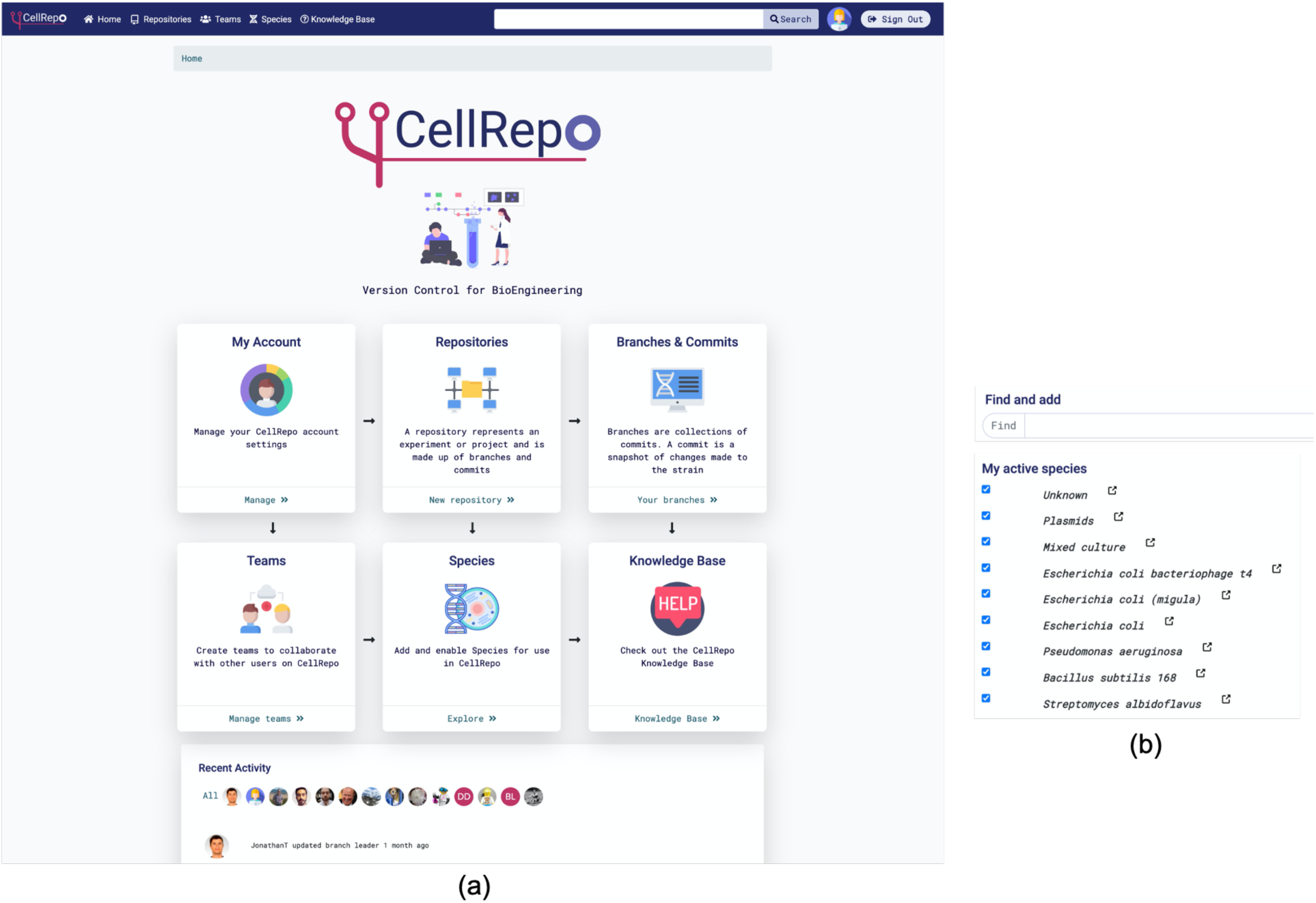
**(a)** Home page after a user signs in. From there they can search and browse their own strain repositories or those they participate as a team member. Also, they have access to any strain repository that has been made public. Users can also create new version control repositories, make commits to them, add new species, etc. The landing page also shows a recent activity registry of the users and the repositories they have access to. **(b)** Species search functionality: users can look up a species database and select the ones they want to use as base for a cell engineering project. If a species is not available in the database users can make a request to add it

The first step to start a repository is to select a species. The server is linked to up-to-date databases of organisms (Fig. 2b). This ensures that the users are always able to use the species they need, and that these are well-documented. To ease the finding of new species to work with, the users need to pre-select them from the database and add the species to their unique list of in-use organisms.

Repositories are projects or experiments (e.g. compound production, protein expression, etc.) and are usually linked to a specific species. Metadata information like the name and description of the project, as well as information about the purpose or how to use a specific strain repository can be added. Repositories may have different “visibilities,”: public (anyone can see the content of the repository), team (visible just for members of the same team or laboratory) and private (just the user can see and add changes). A user may change its repository visibility at any point in time. A repository can have many branches and in turn, branches are made of commits. The name of the initial branch can be set during the creation of the repository.

A repository (Fig. 3a) has a main or leader branch (named during the repository creation) and many other branches. Each branch represents a new direction or idea the users want to try in their projects (e.g. new protocols, different gene edition order, etc.)

**Fig. 3.**
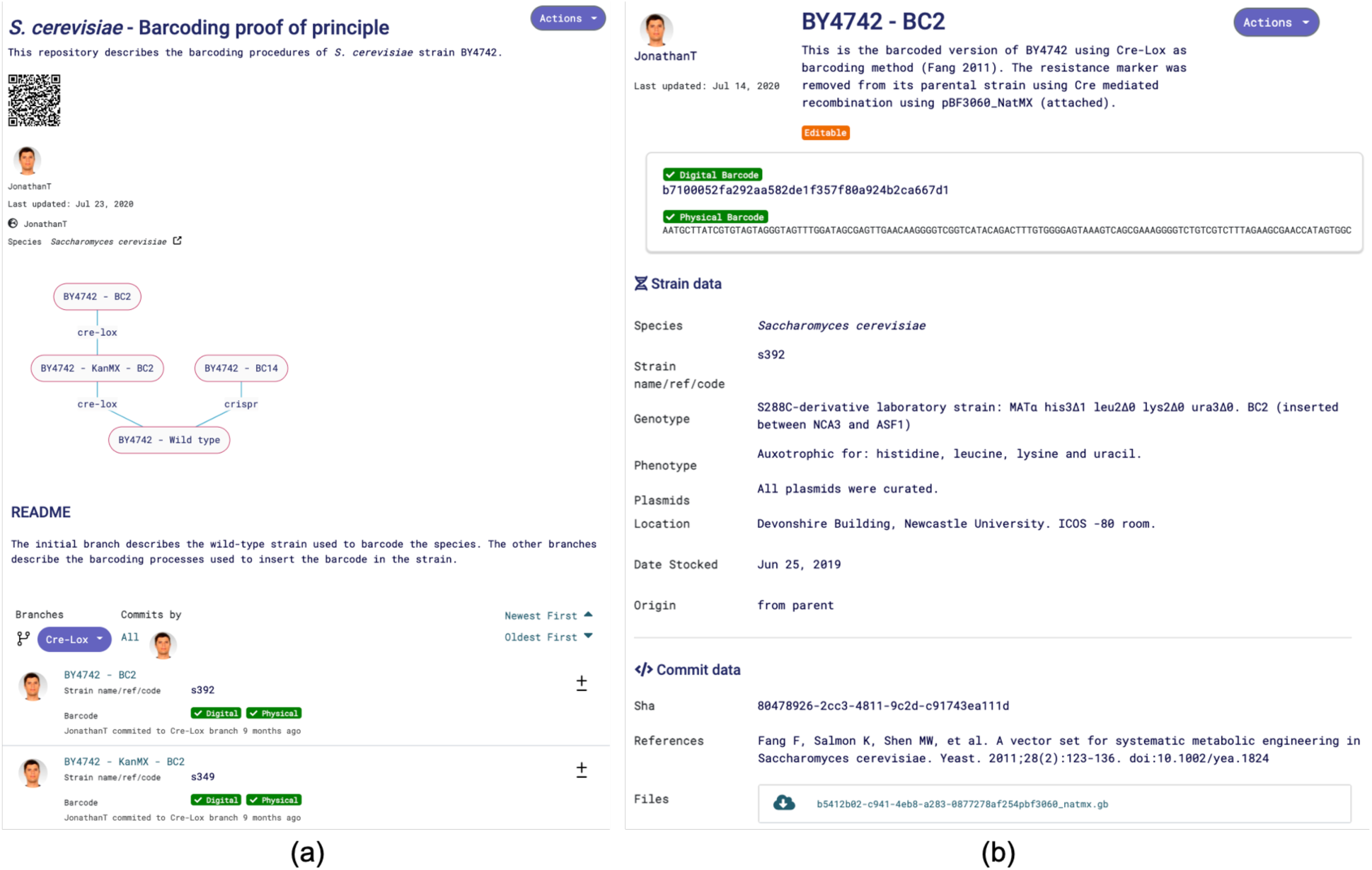
**(a)** Strain Engineering Repositories contain all the digital footprint produced during the engineering of a cell line. The repository provides general information about the cell line project (in this example, the barcoding proof of principle of S. cerevisiae). It also contains all the different “commits” that were made during the engineering process. **(b)** A commit represents a related set of changes introduced into a cell line. All commits have a unique digital identifier and some commits (decided by the user) may also have a physical identifier barcode that is physically inserted into the cell chromosome. Recovering the barcode by sequencing allows a cell engineer to recover the id of the commit containing all the digital footprint of the cell line.

A commit (Fig. 3b) captures the status of the engineered strain at a specific point in time. The amount of information contained in a commit is up to the user (it can be as simple as a new strain name or as complex as a brand-new strain creation by modifying the genome and adding several documents). Additionally, the commits are the containers of the uploaded documentation (which can be in the form of documents, models, sequences, etc.).

Once in a repository, the user can choose a branch to commit. The “new commit” button opens a new form in which various types of information can be inserted. For example, the user can name and describe the commit (what is being done? why? what for?). Importantly, all types of documentation (supporting the commit) can be uploaded in this page such as: construct sequence, electrophoresis gel pictures, SBOL (37) files, growths and fluorescence curves, sequencing results, automation worklist instructions, computer models, etc. The user can also provide genotype and phenotype information, storage location of the strain, safety information, acceptable material transfer agreements for the strain, etc.)

When creating a new commit, the user can decide whether the change is important enough to be physically barcoded into the cell. If that is the case, the system allows the generation of a new unique barcode sequence. The barcode can then be synthesized and inserted in the strain. Once created, the commit will be linked unequivocally to strain carrying the barcode sequence.

CellRepo allows users to be part of collaboration “teams” for cell engineering. Team members of a strain repository can make commits and created new branches to the cell line history.

Furthermore, teams of researchers can share repositories, track strains and be up to date on the experiments being carried out in their projects. Creating a new team is as easy as providing a name to the team and adding CellRepo users to it. Once established, it is possible to see all the members and the shared repositories and keep track of activity taking place on the repository.

## In vivo barcoding experiments

In this work we show that common genomic manipulation techniques could be used to insert the DNA barcodes sequences generated by CellRepo. Different barcoding protocols (detailed step-by-step protocols can be found in Supplementary material) were assessed for *Escherichia coli*, *Bacillus subtilis*, *Streptomyces albidoflavus*, *Pseudomonas putida*, *Saccharomyces cerevisiae* and *Komagataella phaffii* – previously known as *Pichia pastoris*. All the tested barcoding procedures successfully barcoded the target species (Table S5) and links and QRcodes for all CellRepo repositories for these experiments can be found in Table S6. We also carried out stability evaluation of the barcodes. We studied the effects on the strains of both barcode presence and insertion procedures. To do so growth profiles comparisons (fig. S5) and WGS experiments (Tables S7-S12) were carried out. Both experiments suggest that barcoding has no effect on the signed strains. Additionally, the stability of the barcode sequences was assessed under five different growth conditions. The barcodes were stable both in terms of presence (Table S13) and sequence integrity after the long-term experiments (fig. S6-S35). Finally, the usage of barcoded strains as a way to track the dissemination of GMOs was described (fig. S36). In the particular case of a gene drive (which has been proposed as a solution to some infectious diseases transmitted to humans from animal and insect vectors), barcode sequences could pinpoint the source of the released modified organisms (29) (intentionally or accidentally) in the environment.

## Conclusions

Engineering biology, in particular, strain engineering is hard, but the community working on cell engineering makes it harder by not adequately documenting, tracking and sharing the process of genetic engineering. Baker describes (38) the important contribution quality assurance processes can make to research integrity by enabling reproducibility and avoiding the opportunity for cherry-picking of results and data massaging. With this spirit in mind, in this paper we introduce the first version control system for cell engineering. Version control has been a pillar of software engineering and we believe it could substantially improve the quality of the engineering biology too. Widespread adoption of CellRepo (or similar competing resource in the future) will -we postulate-improve:

- **Reproducibility:** It will be easier to reproduce experiments and avoid false leads because one will have a complete long-term change history of every modification to cell lines of interest. This change history includes the author of the change, laboratory of origin, the date of the change and written notes on the purpose and intention of each modification. Having complete history of cell line modification also provides the ability to ‘revert’ back to previous versions of a cell line, which is great for bug fixing in software engineering, and we believe will be useful in cell engineering too. As biologists, this would mean knowing exactly what or someone else did at each commit-able stage in a project. Furthermore, this enables to base, with ease and confidence, a new project on a trustworthy pre-existing repositories and thus absorbing the history that came before it.
- **Traceability:** By physically linking a cell line chromosome to a commit id in the cloud, one is able to know the exact documentation for a strain. Besides technical information about the cell line, stored information also includes the intention behind genetic changes and allows proper allocation of credit (who created a particular commit in a cell line) for work done in the laboratory.
- **Collaboration:** CellRepo improves collaboration. Version control systems allows complex software to be written by single individuals as well as by remote teams. Similarly. CellRepo accommodates both single or multi scientist projects, maintaining a clear record of contributions. Furthermore, CellRepo does not force new workflows into laboratory users. Rather it is agnostic to the specific tools they already use and can accommodate uploads from any laboratory tools they may already be using. CellRepo also allow fine tuning of a cell engineering project visibility by allowing repositories to be entirely private, shareable or public.
- **Transparency:** Our proposed version control system for cell engineering calls for more transparency into the *process of making science*. As we argued, research ought to be transparent and transparency benefits internal team work and enhances public trust in science. CellRepo empowers the sharing of cell line repositories in the same way that version control systems such as GitHub, Bitbucket or Gitlab host and promote open source projects. Like with open source projects, each snapshot of a cell repository shows the “good, the bad, and the ugly” of each stage in the development of a cell line. Through transparency, “bugs in the bugs” would be more readily discovered and corrected. Furthermore, as recently argued (39), there are two growing trends in science. One seeks to make science more open and the other more reproducible, but the adherents of these two trends do not always works *concurrently* towards openness and reproducibility. We believe our paper is a step towards bridging these two camps.
- **Responsibility:** Because key cell lines can be tracked, branched, audited and ownership assigned both digitally and molecularly provenance, quality assurance and trustworthiness are enhanced.
- **Economics:** In software engineering, is possible to develop software without using any version control. However, doing so subjects project owners to data and source code loss risk, loss of project history and the inability to collaborate in real-time. No professional software engineering team should or would accept those risks. Thus, we expect that as CellRepo (or similar competing platform) become the norm in life sciences, important economic benefits will become more tangible.

We have adapted recombineering protocols to barcoding for version control in five bacterial species and one fungal species thus, in principle, one can create a truly universal tracking system for all lab-made cell lines. Altogether, our work demonstrates that barcoding technology can be applied to many industrially and academically relevant microbial species; the barcode sequences are stable under laboratory conditions, they do not affect the growth of the barcoded strains and they can be used as a backtrack signal during GMO dispersion experiments. In the future, other kingdoms of life will also be added to CellRepo and tested in similar ways.

Versioning biological cells will lead to more trustworthy cell engineering. But, perhaps more importantly and as we argued previously, rather than *‘more knowledge of a field leads to more public acceptability’*, what is needed for enhancing public trust of science is ‘*more knowledge about how science is conducted, what is being done and why’.* We believe that CellRepo represents a positive step in that direction.

## Acknowledgments

We acknowledge the Engineering and Physical Sciences Research Council (EPSRC) grant *“Synthetic Portabolomics: Leading the way at the crossroads of the Digital and the Bio Economies (EP/N031962/1)”*, a Royal Academy of Engineering *“Chair in Emerging Technologies”* award to N.K. and the *BioRoboost* (H2020-NMBP-BIO-CSA-2018-820699) Contract of the European Commission.

## List of Supplementary Materials

### Supplementary file.pdf

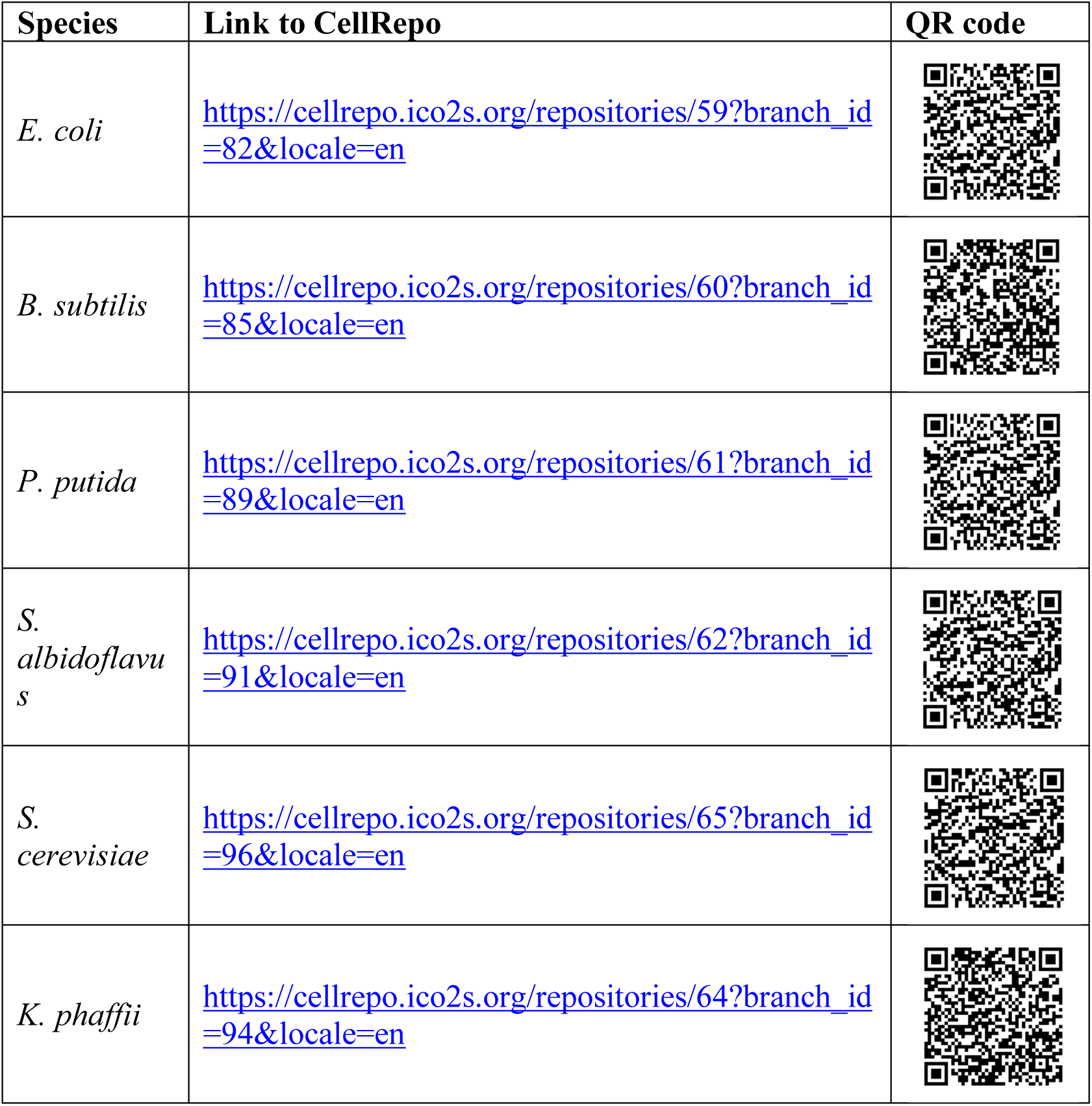

## Supplementary Materials for

### Materials and Methods

#### Barcoding site selection

*E. coli* and *B. subtilis*, have known lists of essential genes (*40, 41*). Using these lists, it was possible to create a simple python script to get possible candidates of essential gene pairs. The script used as input the GFF3 annotation file of the strain and the essential gene list. Both files had to be curated to obtain the same gene nomenclature in both files. As output, the algorithm gives back a pair of essential genes next to each other, the orientation of both genes and the DNA sequence that separates them. Then, databases (*42*) and prediction tools (*43, 44*) were used to check for the presence of regulatory elements in the intergenic sequence not shown in the annotation files. Once a good candidate pair was obtained, the target region was aligned against the most common laboratory strains of the specific species to check for the presence of the possible barcoding region in them.

No list of core essential genes for *P. putida* strains were found in the literature except for conditional essential genes in some conditions (*45*) and essential genes in the related specie *Pseudomonas aeruginosa* (*46, 47, 48*). For this reason, well-known generally-conserved essential genes were taken into consideration to choose one possible insertion locus for the barcode. *GlmS* gene was chosen as a good candidate as it is a broad-host conserved gene in many species and it has a long enough intergenic region for the insertion of Ll.LtrB intron carrying barcodes. The procedure to select the exact insertion site inside the intergenic region downstream of *glmS* was adapted from (*49*). In general, the *PP5408-glmS* intergenic region was surveyed for good Ll.LtrB intron insertion sites in the Clostron.com website. Obviously, the sequences needed for tn7 insertion were avoided as we did not want to hinder the possibility of using this insertion method before or after the barcoding procedure. From the retrieved list of insertion loci, one was chosen from previous data verifying the correct insertion of Ll.LtrB in this site (*50*).

Similarly, to *P. putida*, no essential gene information was found for *S. albidoflavus* (previously known as *Streptomyces albus*). However, in this case we wanted to follow a different approach to showcase the flexibility of the proposed system for new candidate species to be barcoded. A close relative, *Streptomyces coelicolor*, is the model species for the study of the *Streptomyces* genus. For this species, there is a genome-scale metabolic model with gene essentiality data available that can be applied to other *Streptomycetes* (*51*). Using this model, just reactions catalysed by a known *S. coelicolor* gene, essential for all conditions tested and with no isoenzymes were considered. Using this data, it was possible to create a list of putative essential genes (Table S1) for *S. albidoflavus* by aligning each of the *S. coelicolor* A3(2) essential genes to *S. albidoflavus* J1074 database.

Feeding this list to the script described for *E. coli* and *B. subtilis*, however, found no good candidate pair. The number of essential genes next to each other was too low and the few candidate pairs found had problematic intergenic regions. A simpler approach was followed. Using the list of putative essential genes for J1074, the whole genome was analysed looking for clusters of nearby essential genes. In this case, the resulting candidate pair genes were not next to each other but separated by one or more non-essential genes.

To check if the putative essential gene pair was conserved among different *Streptomyces* species, the protein sequences of the chosen candidate genes were aligned against the *Streptomyces* protein database using BLAST to check if the genes were conserved among different species (fig. S1). The results suggest that the two selected genes are conserved and are good candidates for essentiality.

*S. cerevisia*e is well known among the synthetic biology community and there is plenty of well-curated information about it. The user of CellRepo could choose to insert the barcode sequence in already known and curated insertion sites. These sites are used for example in microbial cell factories experiments to insert heterologous genes. By using this type of site, the possibility to barcode a strain, while the user’s desired edition occurs is shown feasible. It was decided to go for an already known insertion site flanked by essential genes used in microbial cell factories experiments (*52*). Also, this site has previously been used to test CRISPR plasmids set in *S. cerevisiae* (*53*).

*K. phaffii* (previously known as *Pichia pastoris*) is known for its ability to produce high amounts of recombinant protein. The alcohol oxidase AOX1 promoter insertion site is commonly used because of its tight regulation and strength (*54*). For these reasons, the AOX1 promoter site was selected as the insertion site in *K. phaffii* without considering closeness to essential genes.

#### Barcoding protocols

Different barcoding methods were designed for each species. For the purpose of this study, all the selection markers were removed from the final strains, except for *K. phaffii*.

For *E. coli*, *B. subtilis*, *S. albidoflavus*, *S. cerevisiae* and *K. phaffii* vectors containing a restriction site to allow the easy cloning of the barcode sequence by Hi-Fi assembly (NEB) using restriction-linearized vectors were used. The barcode DNA sequences were synthesised as dsDNA fragments (Integrated DNA Technologies, IDT) and inserted into the plasmids. The barcoding vectors of *P. putida* were built using the procedures described in (*49*) adapted to this microorganism (50).

Please see Table S2 and Fig. S2 for a detailed description of the vectors used in this study.

To simplify experiments for this paper, just one DNA sequence was used to barcode each species except for *S. cerevisiae* which each of the two barcoding procedures used a different barcode (Table S3). In a real-life scenario, usage of CellRepo produces different DNA sequence barcodes for each commit (to avoid clashes and ambiguity).

The following sections contain a detailed description of the protocols used to barcode each species and the different growth media used in our studies.

*E. coli* and *B. subtilis* barcoding protocols follow (*35*).

##### E. coli barcoding protocols

Two different methods for *E. coli* BW25113: lambda-red recombineering (*55*) and CRISPR-Cas9 mediated edition (testing two different sgRNA sequences) were chosen.

Note: all the barcoding procedures assume that the barcode has been already cloned into the barcode delivering plasmid and/or that the barcoding cassette has been amplified by PCR.

###### *Lambda-red recombineering. Adapted from* (*55*)

Day 1:

1. Start an overnight culture (37 °C) by inoculating LB medium with a single colony.

Day 2:

1. Prepare competent cells following your favourite protocol.
2. Transform *E. coli* cells with plasmid pKD46 and plate the cells at 30 °C in LB supplemented with 100 µg/mL ampicillin (or carbenicillin).

Day 3:

1. Start an overnight culture at 30 °C in LB/Amp from a single colony transformant.

Day 4:

1. Next morning refresh the culture (1:100) with LB/Amp and grow the cells until OD_600_ reaches 0.1.
2. Add arabinose to a final concentration of 30 mM and grow the cells to an OD_600_ =0.5 (recombination proteins are being expressed at this point).
3. Freeze cells on ice for 20 minutes and prepare electrocompetent cells by washing bacteria with ice-cold milli-Q water after spinning aliquots 10 minutes at 5000 rpm in a 4 °C centrifuge.
4. After two washes, resuspend cells in the residual water and electroporate with 500 ng of the barcode DNA cassette (coming from the amplification of pEC-Red2-BC) with a Gene Pulser (25 µF, 200 Ω at 1.8 kV).
5. After electroporating the cells, add 950 µl of fresh LB without antibiotics to samples and resuspended cultures are grown for 2 hours at 37 °C.
6. Plate cells in LB supplemented with chloramphenicol 25 µg/mL.

Day 5:

1. Restreak colonies on LB/CM and grow overnight at 37°C.

Day 6:

1. Perform PCR and sequencing experiments to confirm the insertion of the barcode.
2. To remove the antibiotic cassette, the pCP20 plasmid is transformed. Prepare liquid culture of cells containing the barcoding cassette in LB/Cam.

Day 7:

1. Prepare competent cells and transform pCP20 at 30 °C.
2. Plate in LB/Cam/Amp

Day 8:

1. After pCP20 transformation, inoculate single colonies in LB/Amp/Cam and grow overnight at 30 °C.

Day 9:

1. Next morning, dilute cells in LB and grow at 30 °C until OD_600_ reaches 0.1
2. Swap cells to 42 °C incubator and grow until OD_600_ reaches 0.9.
3. Spot 30 µL in LB plate, streak over the plate and incubated at 37 °C.

Day 10:

1. Barcode presence is checked again by PCR and sequencing.
2. Restreak single colonies in three different plates (LB, Cam and Amp) to check that the resistance is loss.

###### *CRISPR. Adapted from* (*56*)

Note: Different sgRNA sequences were used in different experiments (obtained using (*57*)):

- Final sgRNA1: 5’ - promoter – ATTCCGCGTAAGTATCGCGG– scaffold – terminator 3’
- Final sgRNA2: 5’ - promoter – CGTACAAAAGTACGTGAGGA– scaffold – terminator 3’

Day 1:

1. Start an overnight culture (37 °C) by inoculating LB medium from a single colony.

Day 2:

1. Prepare competent cells following your favorite protocol.
2. Transform *E. coli* cells with plasmid pREDCas9 and plate the cells at 30 °C in LB supplemented with 50 µg/mL spectinomycin.

Day 3:

1. Start an overnight culture at 30 °C in LB/Spec from a single colony.

Day 4:

1. Next morning refresh the culture with LB/Spec and grow the cells until OD_600_ reaches 0.1.
2. Add IPTG to a final concentration of 2 mM and grow the cells to an OD_600_ =0.6 (recombination proteins are being expressed at this point).
3. Frost cells on ice for 20 minutes and electrocompetent cells are then prepared by washing bacteria with ice-cold milliQ water after spinning aliquots 10 minutes at 5000 rpm in a 4 °C centrifuge. (Heat shock transformation also could be performed)
4. After two washes, resuspend cells in the residual milliQ water and electroporate with pEC-CRISPR2-BC.
5. Add 950 µl of fresh LB without antibiotics and resuspend cells. Incubate cells 1 hour at 30 °C.
6. Then, spread 100 µL of bacterial culture on LB plates supplemented with Spec (50 µg/mL) /Amp (100 µg/mL).

Day 5:

1. Check colonies for barcode presence by colony-PCR.
2. Inoculate a positive clone in 2 mL of LB/Spec/Ara (30 mM) (sgRNA targeting pUC origin in pEC-CRISPR2-BC is expressed).
3. Grow for 4-6 hours.
4. Plate on LB/Spec.

Day 6:

1. Check some colonies for Ampicillin resistance by restreaking them on LB/Amp.

Day 7:

1. Take a sensitive clone and restreak it on LB plates at 37°C.

Day 8:

1. Check again by colony PCR the presence of the barcode and restreak on LB/Spec and LB/Amp plates to double check plasmid curing.

##### B. subtilis barcoding protocols

For *B. subtilis* 168, a mazF toxin-based (*58*), a Cre-Lox based (*59*) and a CRISPR based (*60, 61*) engineering methods were selected. The toxin-antibiotic cassette assembly was achieved by HiFi assembly (NEB) of all the parts. The final product was PCR amplified and transformed into 168 cells. pJOE8999.1 plasmid was modified to change the sgRNA promoter to pVeg as in reference (*61*).

###### *mazF toxin-mediated barcoding. Adapted from* (*58*)

Day 1:

1. Inoculate *B. subtilis* cells from a single colony overnight at 37 °C in minimal medium (MM: 10 ml SMM basic salts, 125 µl 40 % (w/v) glucose, 100 µl 2 % (w/v) tryptophan, 60 µl 1 M Mg_2_SO_4_*7H_2_O, 10 µl 20 % (w/v) casamino acids, 5 µl 2.2 mg/ml ferric ammonium citrate)

Day 2:

1. In the morning, dilute cells 1:100 in MM and incubate for 3 h at 37 °C.
2. Dilute 1:2 in SM (SM: 10 ml SMM basic salts, 125 µl 40 % (w/v) glucose, 60 µl 1 M Mg_2_SO_4_*7H_2_O). Incubate 2 h at 37 °C.
3. Mix 400 µL of cells with 1µg of barcode DNA cassette coming from the PCR amplification of the HiFi assembly of all the parts. Incubate 1 h at 37 °C.
4. Plate cells on NA supplemented Zeocin (20 µg/mL) plates. Incubate overnight at 37 °C.

Day 3:

1. Check integration by colony PCR.
2. Incubate overnight one positive clone at 37 °C in LB/0.4% (w/v) glucose/zeocin.

Day 4:

1. Dilute the culture to OD_600_=0.1 in fresh LB/O.4% (w/v) glucose without antibiotics and grow to OD_600_=0.4.
2. Add 1 % (w/v) xylose. Incubate 8 h at 37 °C.

Day 5:

1. Plate cells on NA supplemented with 1 % (w/v) xylose.

Day 6:

1. Restreak individual colonies on NA and NA/Zeocin and colony-PCR to test for cassette removal.

###### *Cre-Lox. Adapted from* (*59*)

Day 1:

1. Start an overnight culture 37 °C in minimal medium.

Day 2:

1. Dilute cells 1:10 in MM and grown for 3 h at 37 °C. Meanwhile, SM is prepared and prewarmed at 37 °C.
2. Dilute in 1:2 SM and make competent cells with a further 2 h incubation period at 37 °C.
3. 400 µl cell aliquots are mixed with 1 µg recombinant DNA PCR (coming from the amplification of pBS-CreLox-BC). Incubate 1 h at 37 °C.
4. Cells are spun down, concentrated and plated on NA/Zeocin (20 µg/mL) and incubated at 37°C.

Day 3:

1. Colonies are tested for the integration of the recombinant DNA by PCR and sequencing.
2. Grow a positive clone overnight at 37 °C in MM.

Day 5:

1. Dilute cells 1:10 in MM and grown for 3h at 37°C. Meanwhile, SM is prepared and prewarmed at 37 °C.
2. The culture is diluted 1:2 in SM and cells are made competent with a further 3 h incubation period at 37 °C.
3. 400 µl cell aliquots are mixed with 100 ng pDR244.
4. Cells are spun down, concentrated and plated on LB supplemented with spectinomycin (100 µg/mL) and incubated at 30 °C.

Day 6:

1. Check selection cassette removal by colony PCR.
2. Positive clones are plated on NA/Zeocin (to check ABR loss) and on LB at 37 °C to cure pDR244.

Day 7:

1. Check pDR244 curation by plating cells on LB/Spec.

###### *CRISPR. Adapted from* (*60*)

Prototype sgRNA: 5’ - promoter – ATTTAGAGCCCTGCCGTGCA – scaffold – terminator 3’ Final sgRNA: 5’ - promoter – GGAAAAGAGTATATTAGATA – scaffold – terminator 3’. sgRNA sequences obtained using (*57*).

Day 1:

1. Start an overnight culture 37 °C in minimal medium.,

Day 2:

1. Dilute cells 1:10 in MM and grown for 3 h at 37 °C. Meanwhile, starvation medium is prepared and prewarmed at 37 °C.
2. Dilute in 1:2 SM and make competent cells with a further 2 h incubation period at 37 °C.
3. Mix 400 µl cell with 400 ng pBS-CRISPR-BC. Spin down cells and plate on LB supplemented with 5 µg/mL kanamycin and 0.2% mannose and incubate at 30 °C for 1 h.

Day 3:

1. Check barcode presence by colony-PCR.
2. Cure positive clones from pJOE-BC by restreaking them at 37 °C in LB plates

Day 4:

1. Store kanamycin sensitive clones as barcoded.

##### P. putida barcoding protocol

The barcoding protocol using TargeTron was adapted from *E. coli* (*10*) to *P. putida* (*50*). The barcode sequence was synthesized by PCR with two overlapping oligonucleotides. Afterwards, this PCR fragment was directly cloned into MluI-digested pSEVA6511-GIIi by Gibson Assembly to generate pSEVA6511-GIIi-BC. The generation of pSEVA231-C-94a is explained elsewhere (*62*). Details on how the TargeTron system works as a barcode deliverer in *P. putida* can be found in Fig. S3. After generating these two plasmids, the barcoding protocol was carried out as follows:

<u>Day 1:</u>

1. Start an overnight culture (30° C) by inoculating LB medium from a single colony.

<u>Day 2:</u>

1. Transform plasmid pSEVA421-Cas9tr into *P. putida* strain either by electroporation (*63*) or conjugation (*64*):

a. <u>Electroporation:</u> Briefly, wash an overnight culture with 300mM sucrose at room temperature four times. At the end, resuspend pellet in 400 µL of sucrose and separate in 100 µL aliquots. After adding 100 ng of pSEVA421-Cas9tr to one aliquot, carry out the electrical shock (program in Gene Pulser from Bio-Rad: 2.5 kV, 25 µF, 200 Ω.) and plate on LB supplemented with streptomycin (100 µg/mL).
b. <u>Conjugation:</u> Tri-parental mating can be performed for the mobilization of pSEVA plasmids from *E. coli* to *P. putida* strains. Briefly, take 1 mL from overnight cultures of donor (*E. coli* strain bearing pSEVA421-Cas9tr), helper (in our case, *E. coli* HB101 with plasmid pRK600) and recipient *P. putida* strain. Spin down cells and wash once with 10mM MgSO4. Mix 100 µL from each washed culture and spin down cells to eliminate the supernatant. Finally, resuspend the final pellet in 20 µL of MgSO_4_ and spot on a LB agar plate. When dry, incubate for at least 4h at 30°C and then, recuperate cells, resuspend them on MgSO_4_ and plate serial dilution on M9 minimal media supplemented with sodium citrate at 0.2% (w/v) and streptomycin.

<u>Day 3:</u>

1. Select single colonies resistant to streptomycin and start a new culture.

<u>Day 4:</u>

1. Transform plasmid pSEVA6511-GIIi-BC by following one of the same two procedures described above.

<u>Day 5:</u>

1. Select single colonies resistant to streptomycin and gentamycin (15 µg/mL)
2. Start a new culture (20 mL of LB supplemented with streptomycin and gentamycin is recommended)

<u>Day 6:</u>

1. Induce the overnight culture with 1mM cyclohexanone for 4h at 30° C
2. Make competent cells by following the procedure explained before Electroporate 100 ng of pSEVA231-C-94a into one aliquot of competent cells and recover the culture for 2 h in LB supplemented with streptomycin.
3. Plate the cells on LB plates supplemented with streptomycin and kanamycin (50 µg/mL).

<u>Day 7:</u>

1. Check by PCR the presence of the barcode at the correct locus and start giving passages with no antibiotic selection to cure plasmids.

##### S. albidoflavus barcoding protocol

pCRISPomyces-2 vector was used to engineer *S. albidoflavus* J1074. Using the vector construction protocol detailed in (*65*), pSA-CRISPR vector was built (testing two different sgRNA sequences). By Hi-Fi assembly, the barcode sequence was cloned. To transform this vector into *S. albidoflavus* J1074, the protocol detailed in (*66*) was followed. pCRISPomyces-2 was a gift from Huimin Zhao (Addgene plasmid # 61737).

Note: Two sgRNA sequences used (obtained using (*57*)):

- sgRNA1: 5’ - promoter – TCATCGTTCTCAATACACCG– scaffold – terminator 3’
- sgRNA2: 5’ - promoter – TGCAACCTCCGTGATCATTC– scaffold – terminator 3’

Day 1:

1. Start an overnight culture of *E. coli* ET12567 (conjugative strain used to transform by the plasmid into *Streptomyces* species) at 37°C in LB supplemented with Chloramphenicol and Kanamycin.

Day 2:

1. Prepare competent cells of *E. coli* ET12567.
2. Transform using your favourite protocol with pSA-CRISPR-BC.
3. Plate in LB supplemented with chloramphenicol (25 µg/mL), kanamycin (50 µg/mL) and apramycin (50 µg/mL).

Day 3:

1. Pick one colony and inoculate 5 mL of LB supplemented with Chloramphenicol, Kanamycin and Apramycin.

Day 4:

1. Pellet down cells.
2. Wash cells 3 times with 1 mL 2xYT media.
3. Heat shock ∼10^8^ *S. albidoflavus* spores in 100 uL of 2xYT at 50°C for 10 min.
4. Use the 100uL spores to resuspend the *E. coli* pellet.
5. Plate on MS-agar supplemented with 20mM MgCl_2_.
6. Incubate overnight at 30°C.

Day 5:

1. Dissolve Apramycin and Nalidixic Acid (which does not act against *S. albidoflavus)* in 1mL H_2_O.
2. Overlay the antibiotic mixture on the plates. Let them dry.
3. Incubate at 30°C for 1 week.

Day 6:

1. Restreak 10 *S. albidoflavus* colonies on MS/Nal at 37°C to cure the plasmids.
2. Grow until colonies appear.
3. Repeat three times.

Day 7:

1. Restreak on MS-agar/Apra to check plasmid curing.

Day 8:

1. Extract genomic DNA of cells using Sigma’s Genomic extraction Kit.
2. Check barcode presence by PCR.

##### S. cerevisiae barcoding protocols

To barcode *S. cerevisiae* BY4742 cells two methods were chosen: Cre-Lox and CRISPR. pSC was built from pCfB2899.

###### *Cre-Lox. Adapted from* (*67*)

pBF3060 was a gift from Nancy DaSilva & Suzanne Sandmeyer (Addgene plasmid # 26850).

Day 1:

1. Start an overnight culture of *S. cerevisiae* cells in YPD.

Day 2:

1. Inoculate 5mL of YPD with 500uL of culture at 30°C.
2. Incubate until OD=0.2-0.3.
3. Prepare competent cells by the LiAc method.
4. Transform with the PCR product coming from the cassette amplification of pSC-CreLox-BC ((both KanMX and URA3 markers were tested)
5. Incubate in YPD for 2 hours (if the selected marker is URA3, this step can be skipped).
6. Plate on YPD supplemented with 200 µg/mL of G418 (or SC -uracil) at 30°C.

Day 3:

1. Wait until colonies appear.
2. Check by PCR barcode presence.
3. Inoculate one positive clone in YPD.

Day 4:

1. Repeat protocol of Day 1 to transform pBF3060_NatMX (cre recombinase expression).
2. Plate on YPD supplemented with 100 µg/mL Neurothreocin.
3. Plate at 30°C.

Day 5:

1. Inoculate 2 mL of appropriate media to maintain selection but with 2% Raffinose /0.1% Glucose as the carbon source.
2. Grow overnight at 30°C.

Day 6:

1. Make a 1/10 dilution using the appropriate selection media supplemented with 2% Galactose and 0.1% Glucose (this induces the expression of CreA).
2. Plate cells in YPD/Neurothreocin.

Day 7:

1. selection cassette removal by colony PCR
2. Inoculate 2 mL of YPD without antibiotic with a positive clone (this step may need to be repeated 2 or 3 days to allow the plasmid curation).
3. Grow overnight at 30°C.

Day 8:

1. Plate on YPD after dilution looking for single colonies.
2. Check the single colonies for Neurothreocin sensitivity (pSC-Crelox-BC curation).

###### *CRISPR. Adapted from* (*53*)

In short, the procedure is based on two plasmids and a PCR repair template.

pCfBf2312, pCfBf2899 and pCfBf3020 were a gift from Irina Borodina (Addgene plasmid # 78231, 73271 and 73282 respectively).

sgRNA: 5’ - promoter – CTCTCGAAGTGGTCACGTGC– scaffold – terminator 3’.

Day 1:

1. Start an overnight culture of *S. cerevisiae* cells in YPD.

Day 2:

1. Inoculate 5mL of YPD with 500uL of culture at 30°C.
2. Incubate until OD=0.2-0.3
3. Prepare competent cells by the LiAc method.
4. Transform with pCfBf2312 (Cas9).
5. Incubate in YPD for 2 hours.
6. Plate on YPD supplemented with 200 µg/mL G418 at 30°C.

Day 3:

1. Wait until colonies appear.
2. Inoculate a positive clone into YPD/G418 at 30°C.

Day 4:

1. Repeat protocol of Day 1 to co-transform pCfBf3020 (gRNA) and the PCR product coming from pSC-CRISPR-BC.
2. Incubate in YPD for 2 hours.
3. Plate on YPD supplemented with Neurothreocin (100 µg/mL) and G418 (200 µg/mL) at 30°C

Day 5:

1. Wait until colonies appear.
2. Check by PCR barcode presence
3. Inoculate a positive clone in YPD without antibiotic (this step may need to be repeated 2 or 3 days to allow the plasmids curation).

Day 6:

1. Plate on YPD after dilution looking for single colonies.
2. Test Neurothreocin and G418 sensitive colonies.

##### K. phaffi barcoding protocol

Adapted from “Thermo Fisher Scientific’s pPICZ A, B, and C transformation protocol (Manual part no. 25-0148)”. A version of pPICZ vector, pICXNH3 (*68*) was used to create our vector. The barcode sequence was inserted after the AOX1 terminator sequence. The vector allows the integration of any desired protein after the AOX1 promoter. SacI was used to linearize the plasmid. pICXNH3 was a gift from Raimund Dutzler & Eric Geertsma (Addgene plasmid # 49020).

Day 1:

1. Digest ∼5–10 µg of barcoded plasmid DNA with SacI
2. Check linearization in agarose gel
3. Column purify the digestion reaction
4. Grow 5 mL of *Pichia pastoris* strain in YPD at 30°C overnight.

Day 2:

1. Inoculate 50 mL of fresh medium with the overnight culture.
2. Grow overnight again to an OD_600_ = 1.3–1.5.

Day 3:

1. Centrifuge the cells at 1,500 × g for 5 minutes at 4°C. Resuspend the pellet with 500 ml of ice-cold, sterile water.
2. Centrifuge again, then resuspend the pellet with 250 ml of ice cold, sterile water.
3. Centrifuge again, then resuspend the pellet in 20 ml of ice-cold 1 M sorbitol.
4. Centrifuge again, then resuspend the pellet in 1 ml of ice-cold 1 M sorbitol for a final volume of approximately 1.5 ml. Keep the cells on ice and use that day. Do not store cells.
5. Mix 80 µL of the cells from Step 6 (previous page) with 5–10 µg of linearized DNA (in 5–10 µL sterile water) and transfer them to an ice-cold 0.2 cm electroporation cuvette.
6. Incubate the cuvette with the cells on ice for 5 minutes.
7. Pulse the cells using the manufacturer’s instructions for *Saccharomyces cerevisiae*.
8. Immediately add 1 ml of ice-cold 1 M sorbitol to the cuvette. Transfer the cuvette contents to a sterile 15-ml tube and incubate at 30°C without shaking for 1 to 2 hours.
9. Spread 10, 25, 50, 100, and 200 µl each on separate, labelled YPDS plates containing 100 µg/ml Zeocin. Plating at low cell densities favours efficient Zeocin™ selection.
10. Incubate plates from 3–10 days at 30°C until colonies form.

Day 4:

1. Pick 10–20 colonies and purify (streak for single colonies) on fresh YPD or YPDS plates containing 100 µg/ml Zeocin.

Day 5:

1. Colony PCR to check barcode presence.

#### Growth curves

For these experiments, all selection markers inserted in the strains were removed (except for *K. phaffii*). All the growth curve experiments were carried out in a CLARIOstar® Plus (BMG Labtech) plate reader using a polystyrene sterile plate, at 300 rpm, using 3 biological replicates per strain. *E. coli* and *B. subtilis* were grown in LB medium at 37°C measuring the absorbance at 600 nm. *P. putida* was grown in LB, at 30°C*. S. cerevisiae* and *K. phaffii* cells were grown at 30° C in YPD medium in 24-well plates. *S. albidoflavus* plate reader experiment was carried out in TS-agar as previously described in (*69*) for *S. coelicolor*.

#### Whole genome sequencing

The genomic DNA was extracted using: GenElute™ Bacterial Genomic DNA Kit Protocol (Sigma) (*E. coli*, *B. subtilis, P. putida* and *S. albidoflavus*) and YeaStar Genomic DNA Kit (Zymo Research) (*S. cerevisiae* and *K. phaffii*).

NGS library was prepared using NEB Next® Ultra™ DNA Library Prep Kit (Cat No. E7370L). Whole genome sequencing was performed on an Illumina NovaSeq 6000 platform at Novogene (Beijing, China).

The reads were aligned against the reference genome of each species (Table S3) using Geneious Prime 2019.2.3 (https://www.geneious.com). First, reads were trimmed using BBDuk (Adapter/Quality Trimming Version 38.37 by Brian Bushnell) and then duplicates were removed using Dedupe (Duplicate Read Remover 38.37 by Brian Bushnell) with default settings in Geneious. The reads were then mapped against the reference genomes using the following settings: Mapper “Geneious”, Sensitivity “Medium/Low/Fast” and selecting “Find structural variants of any size” option. To annotate SNPs, Geneious integrated algorithm with default settings and a “Minimum variant frequency” value of 0.5 was used.

All the mutations also found in the wild type strain genome were discarded from the analysis following the next steps:

(2) Use the comparison tool in Geneious to remove all the SNPs present in the wild type strains.
(3) Manually curate the rest of SNPs, focusing especially on low coverage and repetitive regions (frequent in both S. cerevisiae and K. phaffii). SNPs that flagged in barcoded strains are not in wild type strains and vice-versa. This is because some detected variants qualify as such in one strain but not in the other due to coverage, quality, etc.

#### Long-term barcode survivability test

The stability of the barcode sequence was tested by growing each species during 10 days under five different growth conditions.

**Condition 1.** 10 mL of the growth media (LB for *E. coli*, *B. subtilis* and *P. putida*; TSB for *S. albidoflavus*; YPD for *S. cerevisiae* and *K. phaffii*) were inoculated and grown overnight at 200 rpm. Each morning, during the following 10 days, 100 µL of the culture were re-inoculated in 10 mL of fresh media.

**Condition 2.** 50 mL of the growth media were inoculated and grown at 200 rpm for 10 days.

**Condition 3.** Using a BioXplorer 400 (HEL, London) bioreactor, the cells were grown in 75 mL of growth media with impeller agitation (400 rpm) and filtered air supply (100 mL/min). Cells were grown overnight. After the first night, cells were grown continuously for 10 days at a dilution rate (D) of 0.024 h^−1^ (minimum possible setting of the system). Antifoam 204 (Sigma A6426) was added to the liquid media before autoclaving at 0.01%.

**Condition 4.** Using the same bioreactor system described in the previous condition cells were grown overnight. After the first night, cells were grown continuously for 10 days at a dilution rate (D) of: 0.3 h^−1^ for *E. coli*, *B. subtilis* and *P. putida*; 0.2 h^−1^ for *S. abidoflavus*, *S. cerevisiae* and *K. phaffii*. The dilution rates were inferred from commonly used values for continuous culture (*70*) and previously described growth curves.

**Condition 5.** Three colony replicas of the barcoded strains were restreaked on solid media for 10 passes.

For conditions 1-4, samples were taken periodically and plated on solid media to ensure no contamination had occurred. The last day of the experiments, a sample was taken and spread on agar plates of the same growth media. Single colonies were restreaked. For all conditions, the barcode region was amplified by PCR after genomic DNA extraction and sent to Sanger sequencing (Eurofins Genomics). The sequencing results were aligned against the reference barcode sequence for each species.

The genomic DNA of the microbial population of the conditions 1-4 was isolated as previously described. Also, as a control experiment that did not go through the 10-day culture period, the gDNA of the glycerol stock used to start the long-term experiments was extracted as well. The barcoded region was amplified by PCR and the amplicon was used for NGS analysis. DNA library preparations, sequencing reactions, and initial bioinformatics analysis were conducted at GENEWIZ, Inc. (South Plainfield, NJ, USA). DNA amplicons with partial adapters were indexed and enriched by limited cycle PCR. The DNA library was validated using TapeStation (Agilent Technologies, Palo Alto, CA, USA), and was quantified using Qubit 2.0 Fluorometer and real-time PCR (Applied Biosystems, Carlsbad, CA, USA). The pooled DNA libraries were loaded on the Illumina instrument according to manufacturer’s instructions. The samples were sequenced using a 2×250 paired-end (PE) configuration. Image analysis and base calling were conducted by the Illumina Control Software (HCS) on the Illumina instrument.

Raw Fastq data was first trimmed to remove low quality data using sickle (https://github.com/najoshi/sickle). PANDAseq (https://github.com/neufeld/pandaseq)(71) was then used to merge read1 and read2 of each sample. The merged reads of each sample were mapped to the target reference sequence using BWA (http://bio-bwa.sourceforge.net/) (*72*). Then variants were detected using GENEWIZ’s inhouse script. The primers used for NGS amplicon analysis can be found in Table S4.

#### Yeast gene drive experiments

Special contingency and sterility measures were taken to perform the gene drive experiments. All the experiments were carried out in a Class 2 safety cabinet. All the surfaces were UV-light and chemically sterilized. All the agar plates were sealed using Parafilm.

pGD-ADE2 was assembled containing homologous regions to ADE2 gene, URA3 marker (from BS-Ura3Kl (*73*)), a sgRNA targeting ADE2 and a barcode sequence (both chemically synthesised as gBlocks). A modified version of the protocol detailed in (*74*) was followed. Briefly, the PCR amplification product of the previous cassette was transformed into BY4741 cells. pCfBf2312 (Cas9) plasmid was transformed afterwards. BY4741-GeneDrive-pCfBf2312 cells were mated with BY4742-pXP622 (used just to select diploids) and plated in SC -Leucine containing G418 (200 µg/mL). Diploid cells were then grown in GNA solid media overnight. Cells were transferred to SPOR plates and sporulated following the protocol described in (*75*). Briefly, SPOR plates were incubated at room temperature overnight and at 30°C for 5 days. When tetrads were observed, some cells were scraped from the plate and cell wall digested using Zymolyase solution after incubation at 37°C for 20 min. Tetrads were dissected using SporePlay+ (Sanger Instruments). Single spores were grown on YPD plates until colony formation. Cells were resuspended in water and 5 µL were transferred to GNA and SC -Uracil plates.

pXP622 was a gift from Nancy DaSilva & Suzanne Sandmeyer (Addgene plasmid # 26849). BS-Ura3Kl was a gift from Zhiping Xie (Addgene plasmid # 69195).

##### Recipes of media used in this study

LB (1 L):

- 10 g tryptone
- 10 g NaCl
- 5 g yeast extract
- Up to 1 L distilled water

SMM (1 L):

- 2 g Ammonium sulphate
- 14 g Dipotassium hydrogen phosphate
- 6 g potassium dihydrogen phosphate
- 1 g trisodioum citrate dihydrate
- 0.2 g magnesium sulphate heptahydrate
- Up to 1 L distilled water

*B. subtilis* MM (10 mL):

- 10 mL SMM basic salts
- 125 µL 40% (w/v) glucose
- 100 µL 2% (w/v) tryptophan
- 60 µL 1M Mg2SO_4_*7H_2_O
- 10 µL 20% (w/v) casamino acids
- 5 µL 2.2mg/ml ferric ammonium citrate.

*B. subtilis* starvation media (10 mL):

- 10 mL SMM basic salts
- 125 µL 40% (w/v) glucose
- 60 µL 1M Mg2SO_4_*7H_2_O

M9 minimal medium recipe (1L):

- 6 g Na_2_HPO_4_
- 3 g KH_2_PO_4_
- 0.5 g NH_4_Cl
- 0.5 g NaCl
- 0.2 g MgSO_4_·7H_2_O

NA plates (100 mL):

- 2.8 g oxoid nutrient agar
- Up to 100 mL distilled water

TSB (1 L):

- 30 g Tryptic Soy Broth powder
- Up to 1 L distilled water

2xYT (1 L)

- 16 g tryptone
- 10 g yeast extract
- 5 g NaCl
- Up to 1 L distilled water.

MS plates (1 L):

- 20 g mannitol
- 20 g soy flour
- 20 g agar
- Up to 1 L distilled water

YPD (1 L):

- 20 g peptone
- 10 g yeast extract
- Up to 1 L distilled water
- Add glucose to 2% final concentration after autoclaving

SC dropout plates (100 mL) (compounds from Formedium):

- 690 mg Nitrogen base without amino acids
- 162.2 mg of Leucine dropout or 192.6 mg of Uracil dropout mixture
- 2.4 g agar
- Up to 100 mL

SC dropout plates without ammonium sulphate (100 mL) (compounds from Formedium):

- 190 mg Nitrogen base without amino acids and without ammonium sulphate
- 162.2 mg of Leucine dropout or 192.6 mg of Uracil dropout mixture
- 2.4 g agar
- Up to 100 mL

GNA plates (100 mL):

- 5 g D-glucose
- 3 g Nutrient broth
- 1 g yeast extract
- 2.4 g agar
- Up to 100 mL distilled water

SPOR plates (100 mL):

- 1 g potassium acetate
- 100 mg yeast extract
- 50 mg glucose
- 2.4 g agar
- Up to 100 mL distilled water

### Supplementary Text

#### Barcoding site selection and insertion

In previous works (*11*) we investigated a barcode location for both *E. coli* and *B. subtilis* strains based basically on the non-proximity to crucial operons for the cell proper functioning. We wanted, in the current work, to try a different approach. For the barcode to be stable two factors come to play: possible mutations in the barcode DNA sequence and total or partial loss of the barcode sequence.

To tackle the later we decided to investigate in this case if it was possible to barcode strains using insertion sites close to or in between essential genes. In theory, the proximity of essential genes would “protect” the barcode of any recombination event that may occur (*7*). Table S3 and Fig. S4 describe the insertion sites used for all the species in this study.

Combined with the already described (*11*) alternative, different solutions to the insertion site question have been explored: in conserved silent regions, between or close to essential genes and using previously known insertion sites useful for different purposes. The decision of where to insert the identifier sequence would, in the end, be made by the users of the system depending on what is the final goal of the strain, on their experience manipulating its genome and on their time availability.

The different methods used for each species were proved useful to insert the barcode sequence. Both the insertion efficiency and the PCR and sequencing tests showed that these small DNA identifiers can be inserted using any of the described methods with high efficiency. Table S5 shows the efficiency of each barcoding method used in each species.

Six different repositories were created to document the different barcoding procedures. They act also as proof-of-principle to demonstrate their usability for biotechnology related purposes. Links to the repositories can be found in Table S6.

##### The effect on barcoded strains

The system proposed here should minimise the effect on the modified strain, its performance and suitability for a specific experiment or industrial process. In our previous work, initial experiments on the stability of the barcode were described. In this article, this crucial identifier feature was assessed in two ways: growth curves and WGS.

First, the growth profile of barcoded cells was compared to the one of the wild type strains. Barcoding a strain should have little to no effect on its growth profile (Fig. S5).

The six growth profiles show no big differences between barcoded strains and the corresponding non-barcoded parental strains. This confirms that the barcode insertion has little effect on the growth of the different species.

In addition to the physiological test, a general idea of the genomic mutations one should expect after the barcoding process had ended was needed (either naturally occurring mutations caused during the process or mutations that are specifically caused by the barcoding methodology). For instance, this can be helpful in choosing a specific barcoding method over another.

To do this, the whole genome of the barcoded strains and the wild type strains used was sequenced. The results can be found in Tables S7-S12.

*E. coli* results show that the three clones barcoded using Lambda-Red method had the same point mutation in an intergenic region (Table S7). This may be explained by the fact that the initial colony chosen to start the insertion process already contained the mutation or it was acquired during the process. In any case the mutation is intergenic and does not seem to affect the cells. In one of the clones barcoded using gRNA1, two-point mutation appear in different CDS. Strains barcoded using gRNA2 do not show any mutation.

*B. subtilis* was barcoded using three different methods. Both CRISPR (only one gRNA tested) and Toxin-mediated barcoded cells show no mutations in all the different clones. In one of the barcoded strains using Cre-Lox, a mutation appears. In a different clone, two different SNPs could be detected. All the mutations are in different CDS (Table S8).

For *P. putida* two clones were barcoded using CRISPR/targetron system and both showed one mutation in different CDS (Table S9).

*S. cervisiae* was barcoded using two methods. In both of them, most of the mutations that appear are tandem repeat related. These could be acquired during the insertion process or could be sequencing artefacts related to this type of repetitive sequences. Two strains barcoded using Cre-Lox show single point mutations. Four mutations (tandem repeats) are observed in Strain 2 (CRISPR). A single point SNP can be observed in Strain 3 (Table S10).

One of the two strains of *K. phaffii* show two-point mutations in intergenic regions (Table S11).

Finally, the NGS analysis of *S. albidoflavus*, shows the larger number of variants. Strain 1 barcoded with gRNA1, shows two different base pair changes in different CDS. However, no mutations were found for Strain 2 (gRNA1) (Table S12). The three strains produced using gRNA2 count six, three and two SNPs respectively.

Next, we describe a plausible explanation for the higher variation observed in *S. albidoflavus* NGS analysis.

*S. albidoflavus* was barcoded using CRISPR. In eukaryotes, CRISPR-caused double strand breaks (DSB) can be repaired by Non-Homologous End Joining (NHEJ) or Homologous Recombination (HR) in the presence of a repair template. NHEJ repair is usually imprecise and indels occur. In the case of *S. cerevisiae* however, it has been observed that NHEJ hugely decreases the cell survival and, when a repair template is provided, HR is the prevalent repair mechanism in this species (*76*). Prokaryotes usually lack NHEJ repair mechanisms. Nevertheless, it has been described in *Streptomyces coelicolor* among other bacterial species. In this *Actinobacteria*, closely related to *S. albidoflavus*, researches knocked-out genes using CRISPR without providing a repair template and allowing the native NHEJ system to – wrongly – repair the DSB (*77*). It may be possible that the gRNAs caused CRISPR off-target activity that was repaired by NHEJ causing mutations to appear. Together with the fact that the genome of *S. albidoflavus* has a high GC content and produced the worst sequencing quality of the analysed species, this may explain the higher mutation count. Even though other mechanisms may explain the observed variation (see next), users wanting to use pSA-CRISPR-gRNA2 should bear in mind these extra mutations.

The NGS analysis shows that for all the sequenced strains the number of total mutations found in each method is low. The mutations are not constant in the different replicas sent to sequencing. We hypothesise that the mutations (if not sequencing artefacts) are caused by the natural mutation rate in each species during several cycles of growth (both in liquid and solid media). This is supported by (*77*). In this *S. coelicolor* CRISPR edition experiment, the control strain, in which an empty (no target) gRNA was provided to the cells caused a total of seven mutations, three of which were in coding regions. Similar results were found in a CRISPR experiment in *S. cerevisiae* where they detected 10 SNPs that were probably caused by the successive transformation rounds required for the experiment (*78*).

No structural variants were found for any of all the sequenced strains.

The NGS analysis suggests that the barcoding procedures do not change the genome of the strains more than what would be expected while carrying out conventional genetic engineering protocols. CellRepo users can use this information to choose the barcoding method specific for each species that best fits them.

##### Barcode survival after long-term growth

The barcode sequence needs to be stable over time. In our previous work (*11*) it was confirmed that the barcode in *E. coli* and *B. subtilis* is stable during ∼200 generations in exponential phase and after 10 subcultures. In this work, it was tested if the DNA sequence is also stable during continuous stationary phase growth and other non-exponential growth profiles, common in laboratory and industrial processes like batch fermentation growth and restreaks on solid media. Stationary phase mutagenesis occurs to microorganisms when they are deprived from nutrients. Mutations may arise without active cell division or global DNA replication (*79, 80*). This phenomenon has been demonstrated in *E. coli* (*81, 82*), *B. subtilis* (*83*), *P. putida* (*84*) and *S. cerevisiae* (*85, 86*).

To assess that the barcode stays in the insertion site and that its sequence is still retrievable even after long periods of growth, we ran on all six species five different experiments for ten days.

As a preliminary experiment, for each condition, the final day single colonies were re-streaked and the barcode region was PCR amplified and sent to Sanger sequencing (Eurofins Genomics). For all the colonies tested, we were able to confirm the barcode presence by PCR in all the cases. Table S13 describes in detail the sequencing results of this experiment.

To have a more thorough view of what happened to the barcode during the long-term experiments, an NGS analysis of the PCR purified product of the barcoded region of the cell population in the conditions 1-4 was performed.

Mutation analysis of the barcode sequences for all species can be found in Fig. S6-S35.

*E. coli* control (Fig. S6) showed a base pair change in 50% of the reads. To check if the glycerol stock had any mutation 10 colonies were isolated and the PCR product of the barcode region sent to Sanger sequencing. No mutations were found. A point mutation in the initial PCR cycles of the reaction sent to NGS may explain this result.

The percentage of reads showing either INDELs or base-pair changes stayed at the same value as the one observed in the control experiment (lower than 0.05%).

Both the single colonies and the population level experiments show that the barcode was still present after the long-term incubation period and that the sequence was stable on all five experimental conditions.

##### Barcodes provide a backtrack signal for GMO dispersion experiments

Gene drive technology allows the researchers to propagate a specific genetic modification through a population (*87, 88*). The scientific community needs to assess the risk of this kind of research. Barcode sequences can be helpful in this matter and uniquely identify the laboratories where a gene drive experiment was carried out, the purpose of the modification and any other relevant data (e.g. safety measures implemented).

Fig S36a graphically describes the gene drive molecular mechanism. The barcode identifier was coupled with the intended modification (ADE2 deletion cassette). Fig. S36b shows that the barcoded cells (red pigment) can survive in SC -Uracil media. Haploid cells coming from the unmodified parent cell show red pigment only when pCas9 plasmid was also present. In all cases, it was possible to PCR and sequence the barcode sequence from each haploid individual.

---

**Fig. S1.**
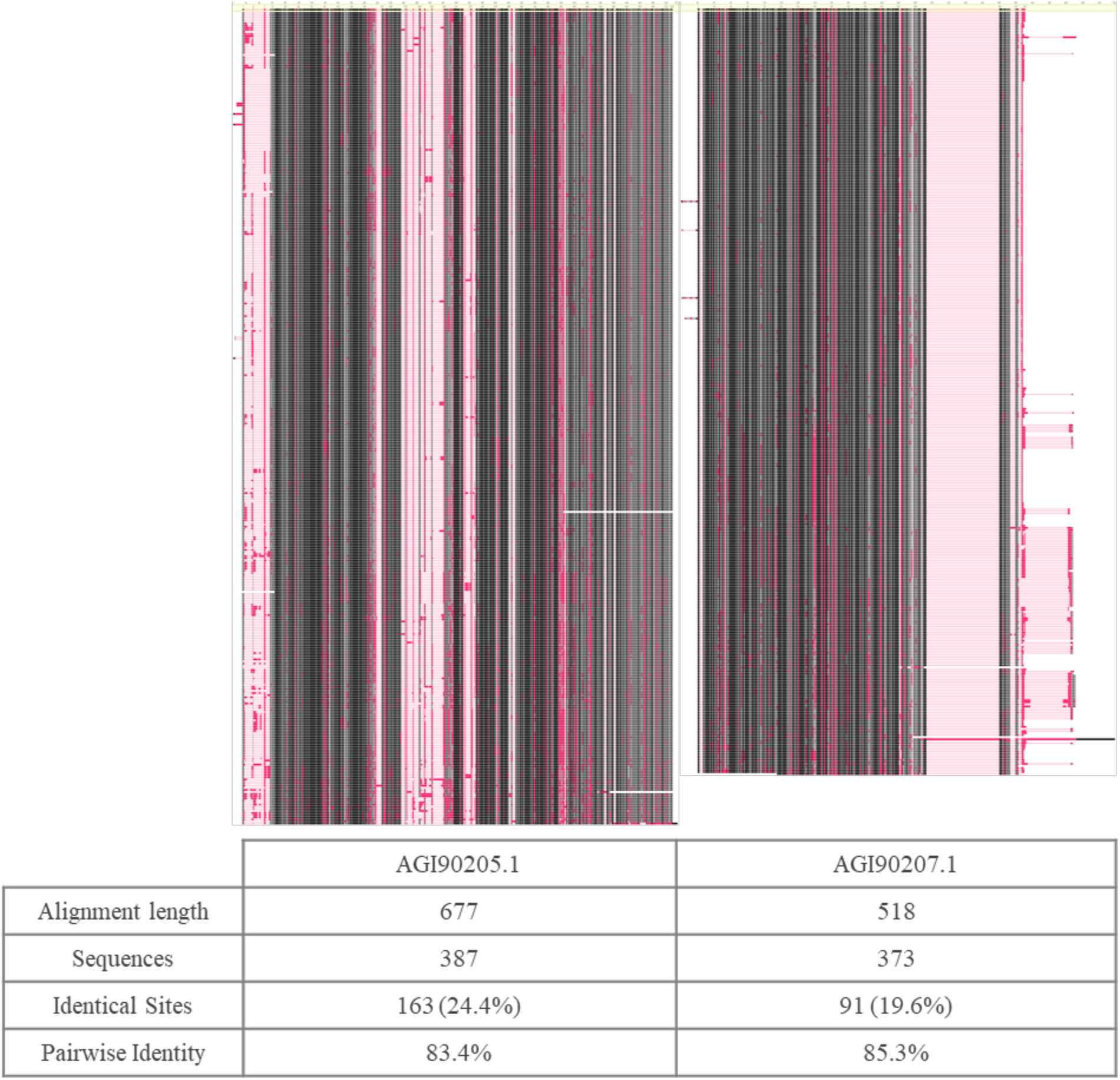
Conservation pattern of the putative essential gene pair AGI90205.1 (IMP cyclohydrolase / Phosphoribosylaminoimidazolecarboxamide formyltransferase) and AGI90207.1 (Bifunctional protein FolD) (encoded by loci XNR_3870 and XNR_3872 respectively). Both protein sequences were aligned against the classified Streptomyces database using BLAST. The hits were aligned using MAFFT included in Geneious Software package. Black colour indicates a higher similarity. Pink colour indicates a lower similarity. Spaces indicate alignment gaps. The alignment statistics were calculated by Geneious.

**Fig. S2.**
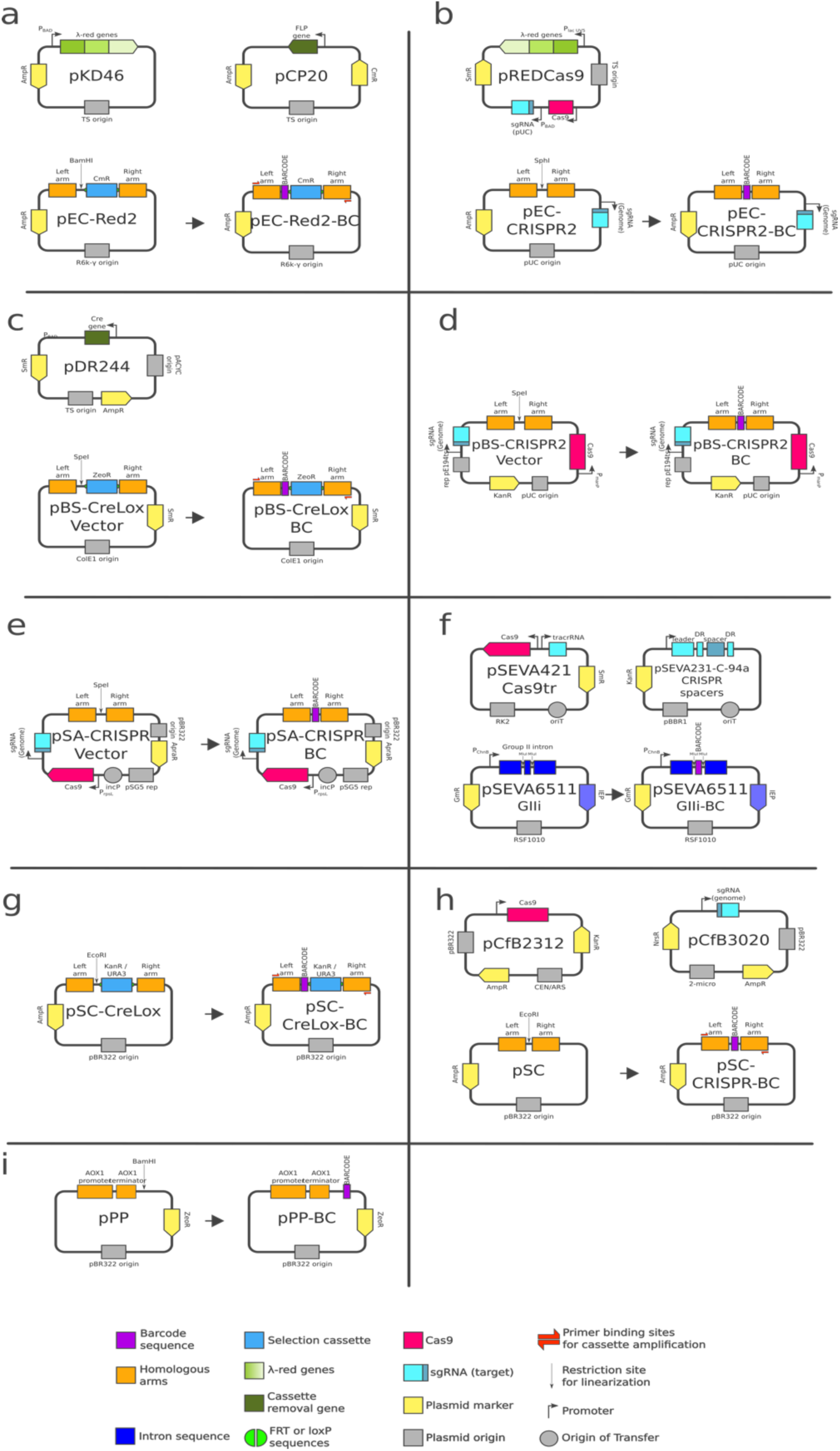
Maps of the barcoding plasmids used in this study. a) *E. coli* lambda-red. b) *E. coli* CRISPR. c) *B. subtilis* Cre-Lox. d) *B. subtilis* CRISPR. e) *S. albidoflavus* CRISPR. f) *P. putida* CRISPR/targetron. g) *S. cerevisiae* Cre-Lox. h) *S. cerevisiae* CRISPR. i) *K. phaffii* AOX1 insertion.

**Fig. S3.**
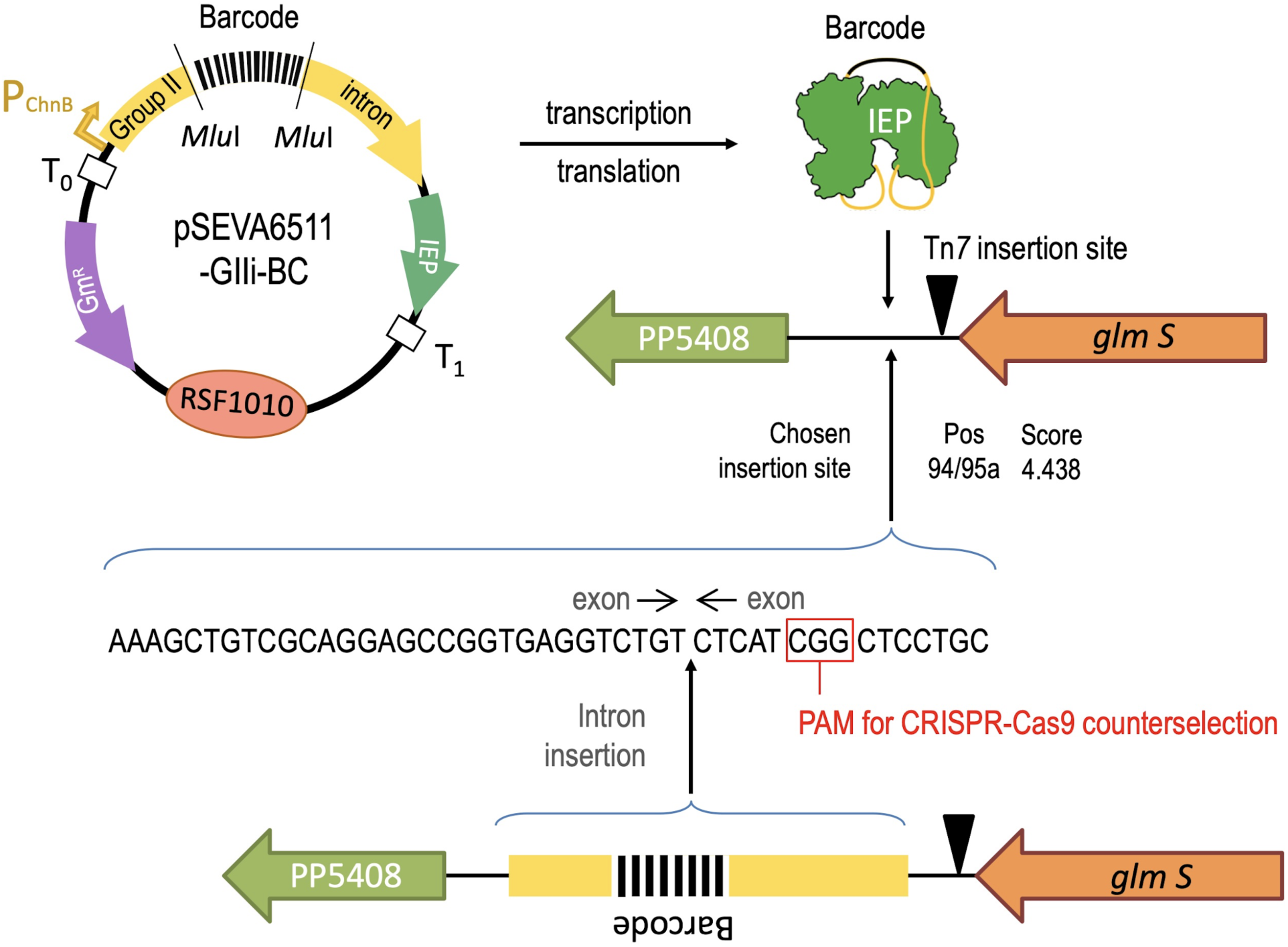
Insertion of a barcode in a permissive site of the genome of *Pseudomonas putida*. Once committed, the barcode was assembled as part of the Group II intron sequence of plasmid pSEVA6511-GIIi-BC, which co-transcribes the corresponding RNA along with an intron-encoded protein (IEP) with reverse transcriptase activity. Upon induction of the expression system with cyclohexanone, a ribonucleoprotein (RNP) complex is formed that contains the excised intron RNA and IEP. After RNA splicing, the group II intron RNP recognizes DNA target sequences for intron insertion by using both the IEP and base pairing of the intron RNA. In our case the site of insertion is chosen in the proximity of the Tn7 insertion site and is predicted to occur with the target identification algorithm in the site indicated. Note also the PAM site exploited for counterselection of the non-inserted, wild-type region with the CRISPR/Cas9 system explained in the text.

**Fig. S4.**
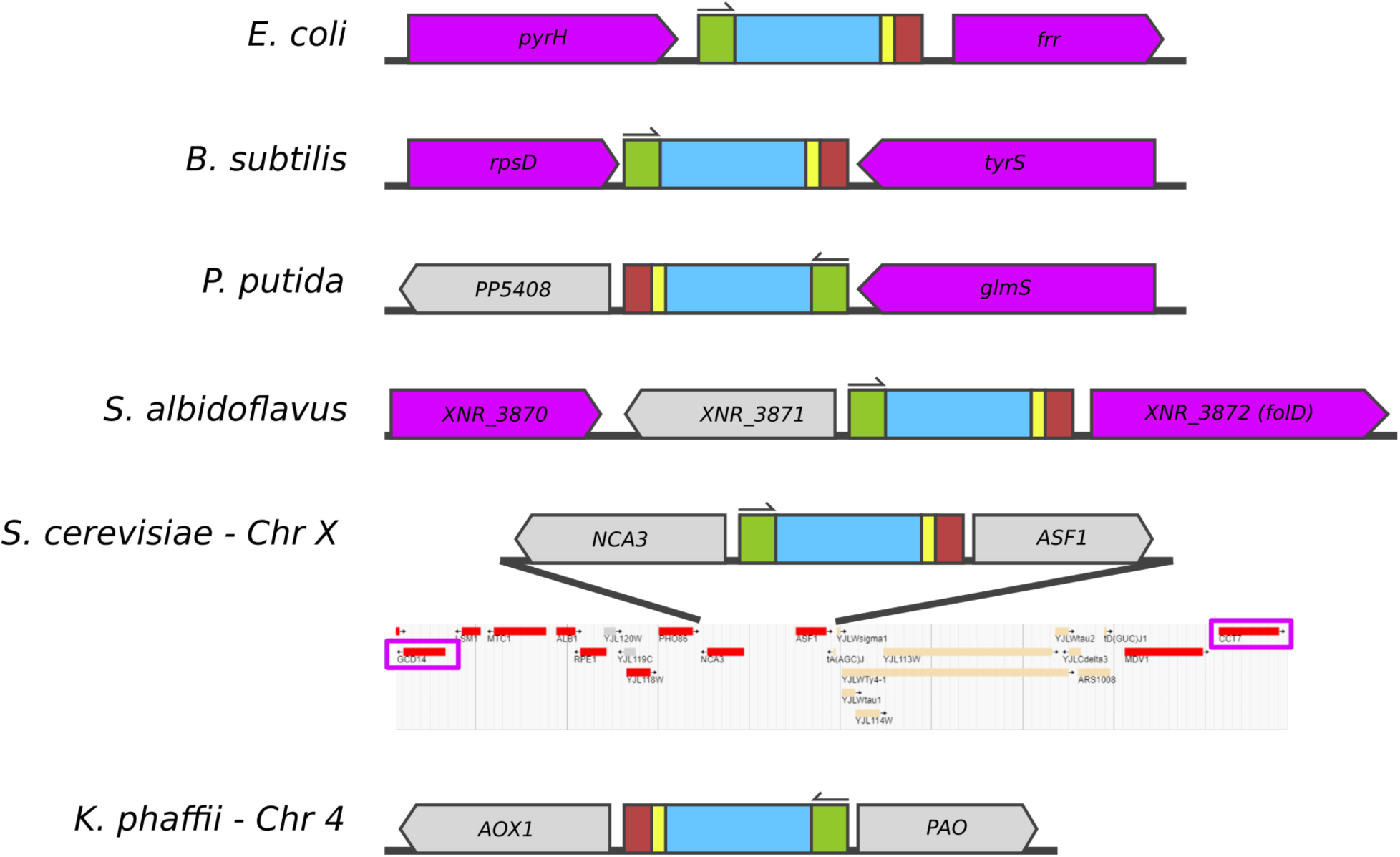
Barcoding locations used in this study. Purple colour indicates essential genes. Universal sequencing primer site (green), barcode sequence (blue), synchronisation sequence (yellow) and checksum (red). S. cerevisiae genomic map was obtained from https://www.yeastgenome.org/

**Fig. S5.**
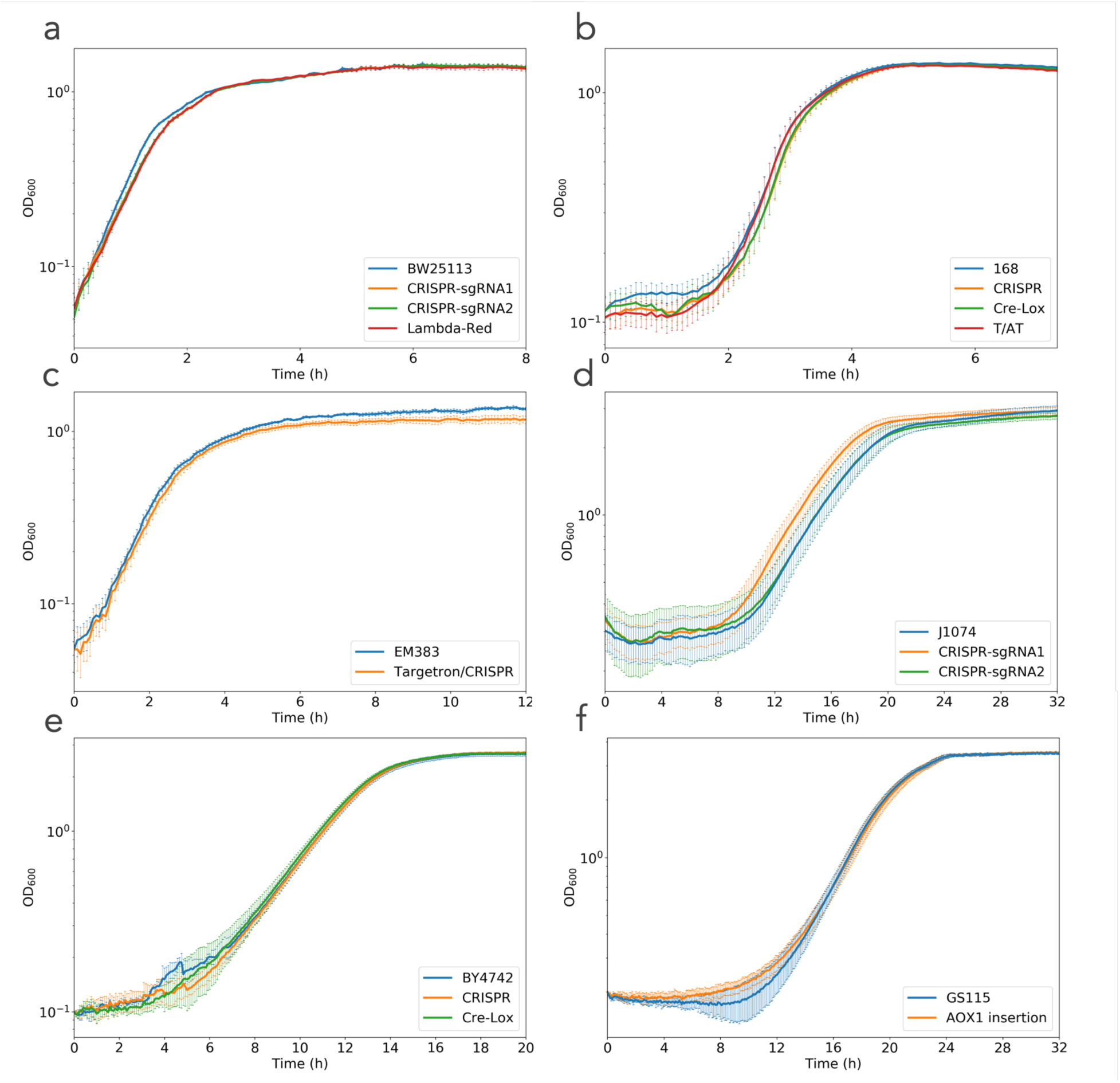
Growth profile comparison between wild-type strains and barcoded strains. Wild-type strains are shown in blue. a) *E. coli.* b) *B. subtilis*. c) *P. putida*. d) *S. albidoflavus*. e) *S. cerevisiae*. f) *K. phaffii.* Note that the variation in S. albidoflavus graph can be explained because it was grown on solid tryptic soy agar.

**Fig. S6.**
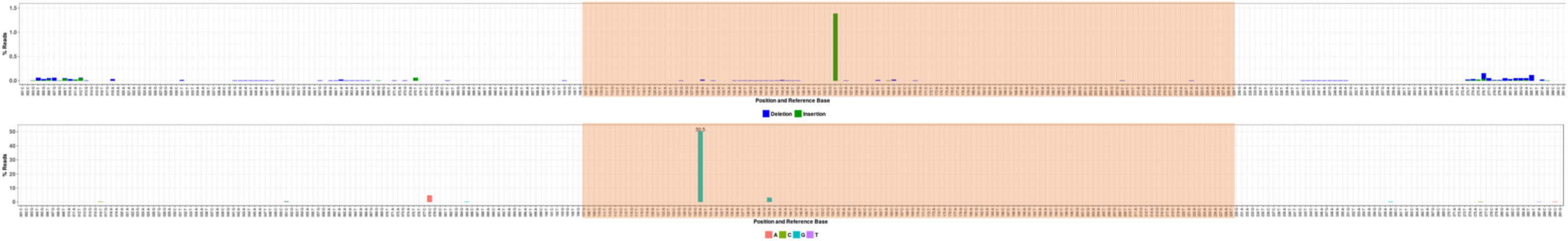
*E. coli* amplicon NGS. Glycerol control. Barcode sequence indicated by the orange region. Upper graph: percentage of reads showing a specific deletion/insertion. Lower graph: percentage of reads showing a specific base pair change. ∼17000 read depth.

**Fig. S7.**
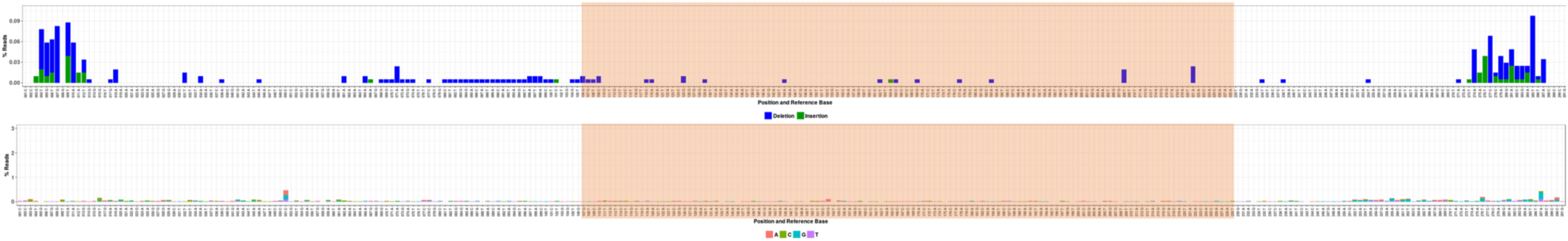
*E. coli* amplicon NGS. Condition 1. Barcode sequence indicated by the orange region. Upper graph: percentage of reads showing a specific deletion/insertion. Lower graph: percentage of reads showing a specific base pair change. ∼19500 read depth.

**Fig. S8.**
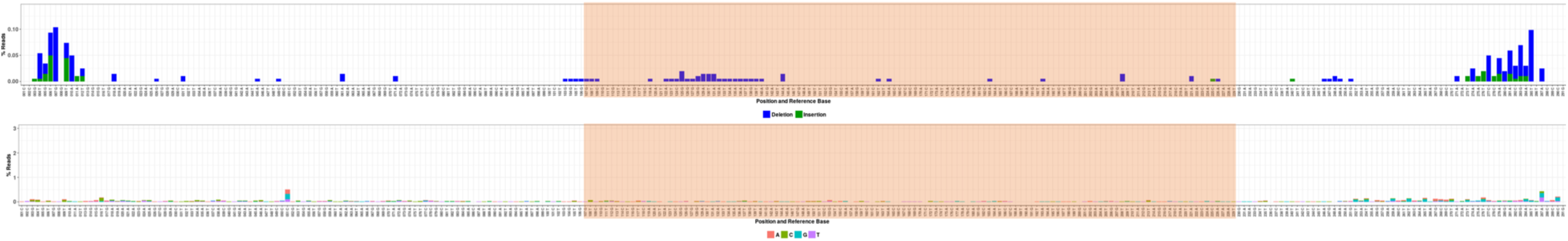
*E. coli* amplicon NGS. Condition 2. Barcode sequence indicated by the orange region. Upper graph: percentage of reads showing a specific deletion/insertion. Lower graph: percentage of reads showing a specific base pair change. ∼19200 read depth.

**Fig. S9.**
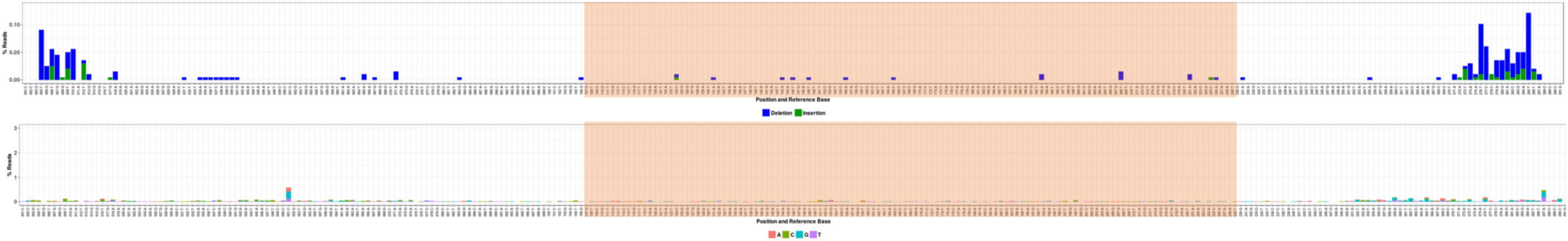
*E. coli* amplicon NGS. Condition 3. Barcode sequence indicated by the orange region. Upper graph: percentage of reads showing a specific deletion/insertion. Lower graph: percentage of reads showing a specific base pair change. ∼18700 read depth.

**Fig. S10.**
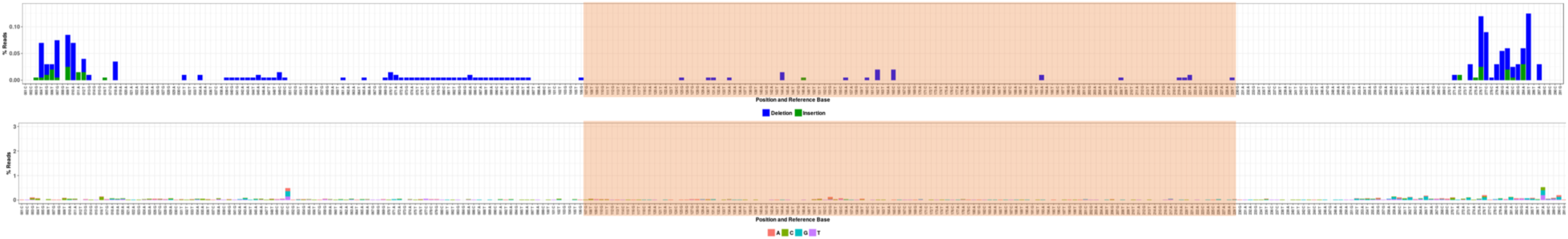
*E. coli* amplicon NGS. Condition 4. Barcode sequence indicated by the orange region. Upper graph: percentage of reads showing a specific deletion/insertion. Lower graph: percentage of reads showing a specific base pair change. ∼19200 read depth.

**Fig. S11.**
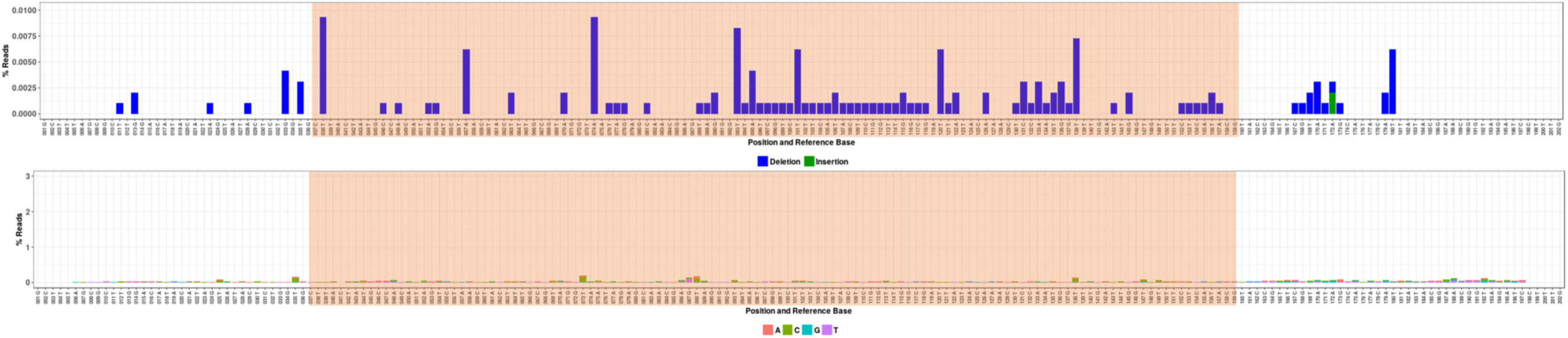
*B. subtilis* amplicon NGS. Glycerol control. Barcode sequence indicated by the orange region. Upper graph: percentage of reads showing a specific deletion/insertion. Lower graph: percentage of reads showing a specific base pair change. ∼95500 read depth.

**Fig. S12.**
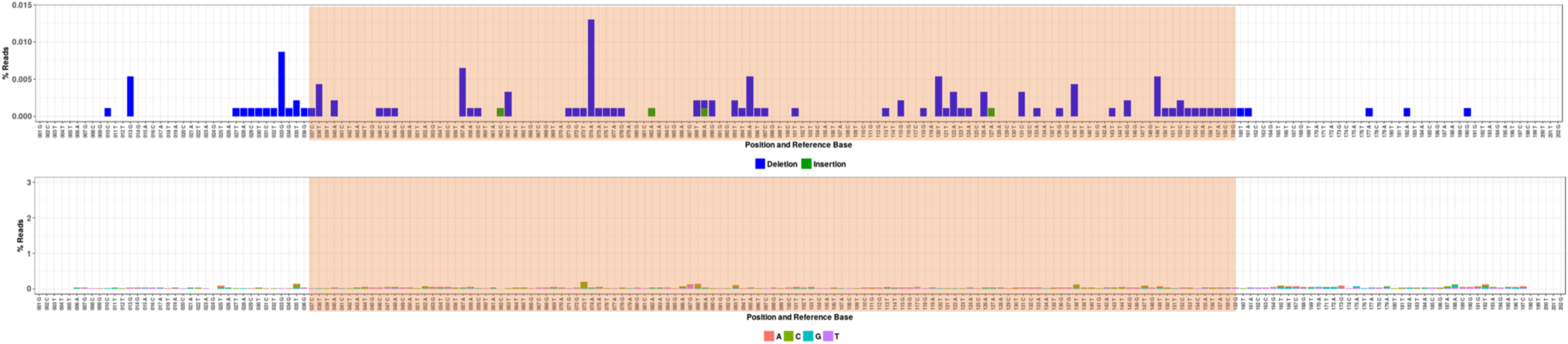
*B. subtilis* amplicon NGS. Condition 1. Barcode sequence indicated by the orange region. Upper graph: percentage of reads showing a specific deletion/insertion. Lower graph: percentage of reads showing a specific base pair change. ∼91400 read depth.

**Fig. S13.**
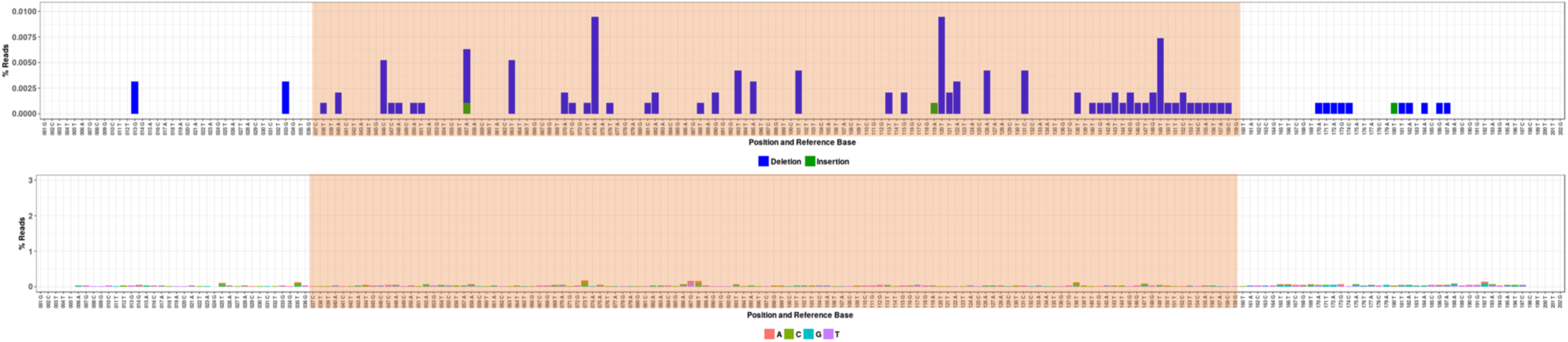
*B. subtilis* amplicon NGS. Condition 2. Barcode sequence indicated by the orange region. Upper graph: percentage of reads showing a specific deletion/insertion. Lower graph: percentage of reads showing a specific base pair change. ∼94300 read depth.

**Fig. S14.**
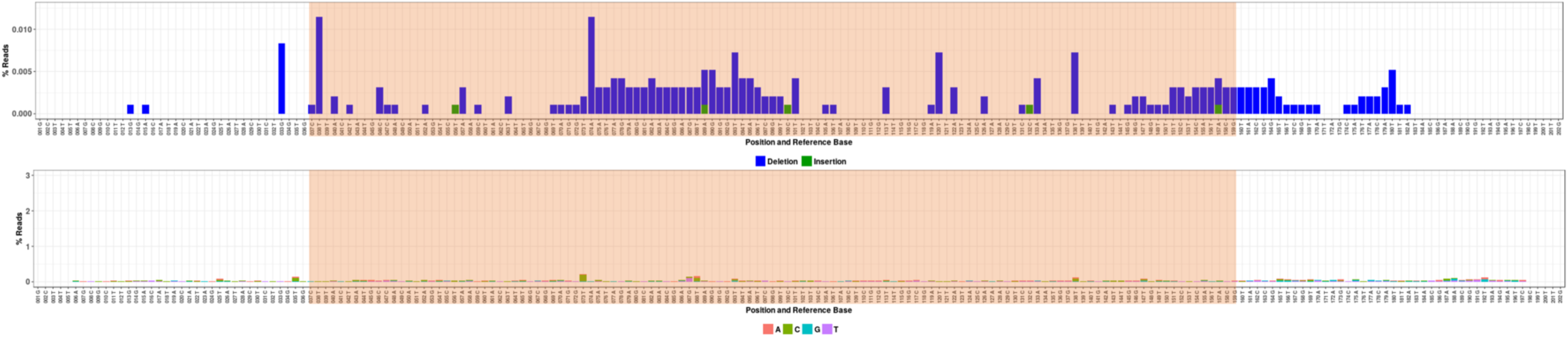
*B. subtilis* amplicon NGS. Condition 3. Barcode sequence indicated by the orange region. Upper graph: percentage of reads showing a specific deletion/insertion. Lower graph: percentage of reads showing a specific base pair change. ∼95000 read depth.

**Fig. S15.**
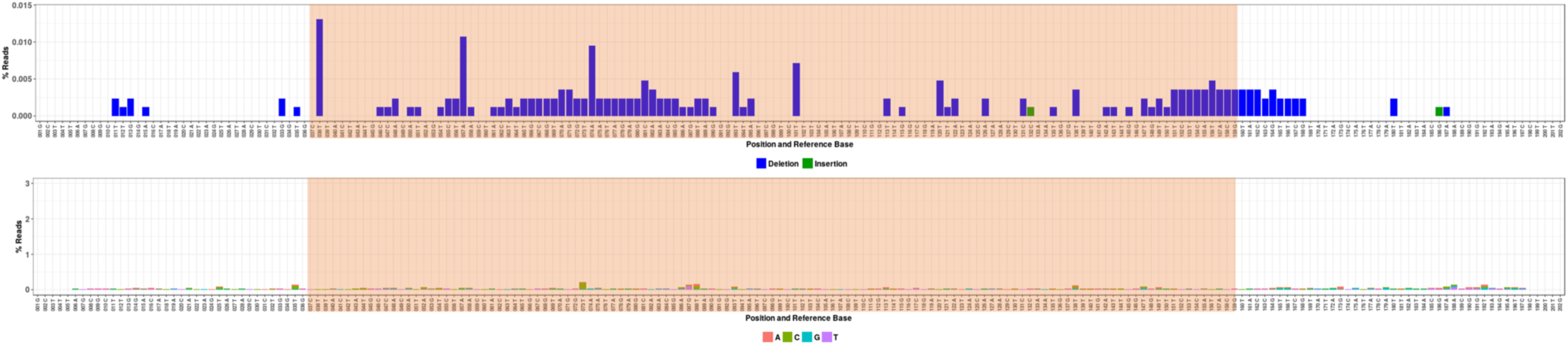
*B. subtilis* amplicon NGS. Condition 4. Barcode sequence indicated by the orange region. Upper graph: percentage of reads showing a specific deletion/insertion. Lower graph: percentage of reads showing a specific base pair change. ∼83200 read depth.

**Fig. S16.**
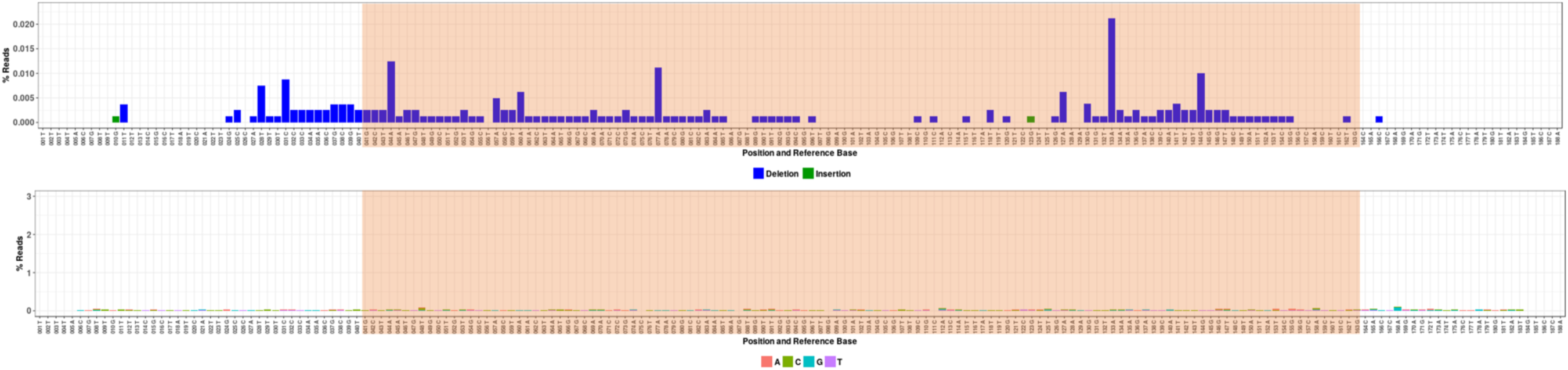
*P. putida* amplicon NGS. Glycerol control. Barcode sequence indicated by the orange region. Upper graph: percentage of reads showing a specific deletion/insertion. Lower graph: percentage of reads showing a specific base pair change. ∼79900 read depth.

**Fig. S17.**
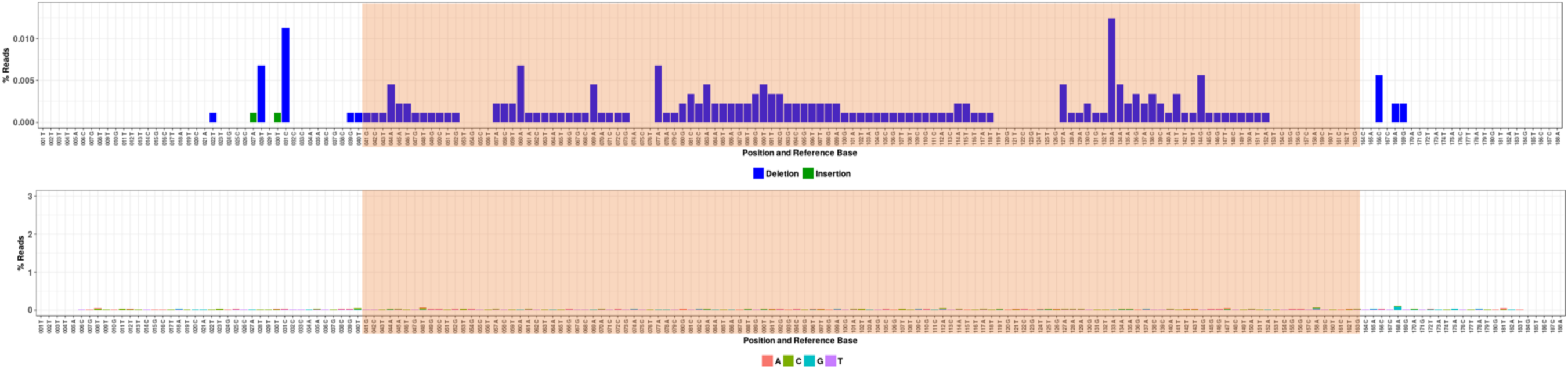
*P. putida* amplicon NGS. Condition 1. Barcode sequence indicated by the orange region. Upper graph: percentage of reads showing a specific deletion/insertion. Lower graph: percentage of reads showing a specific base pair change. ∼88300 read depth.

**Fig. S18.**
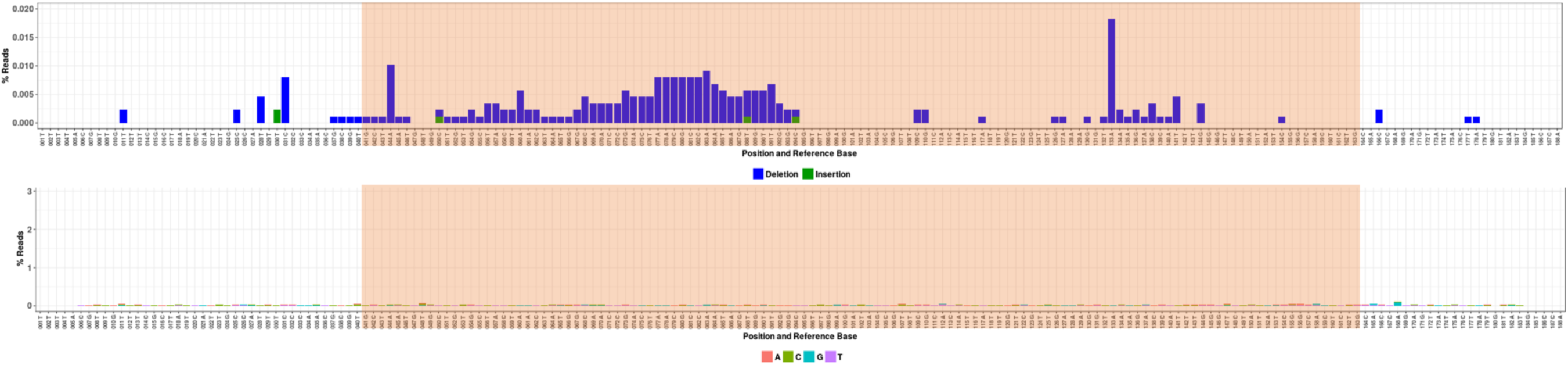
*P. putida* amplicon NGS. Condition 2. Barcode sequence indicated by the orange region. Upper graph: percentage of reads showing a specific deletion/insertion. Lower graph: percentage of reads showing a specific base pair change. ∼87400 read depth.

**Fig. S19.**
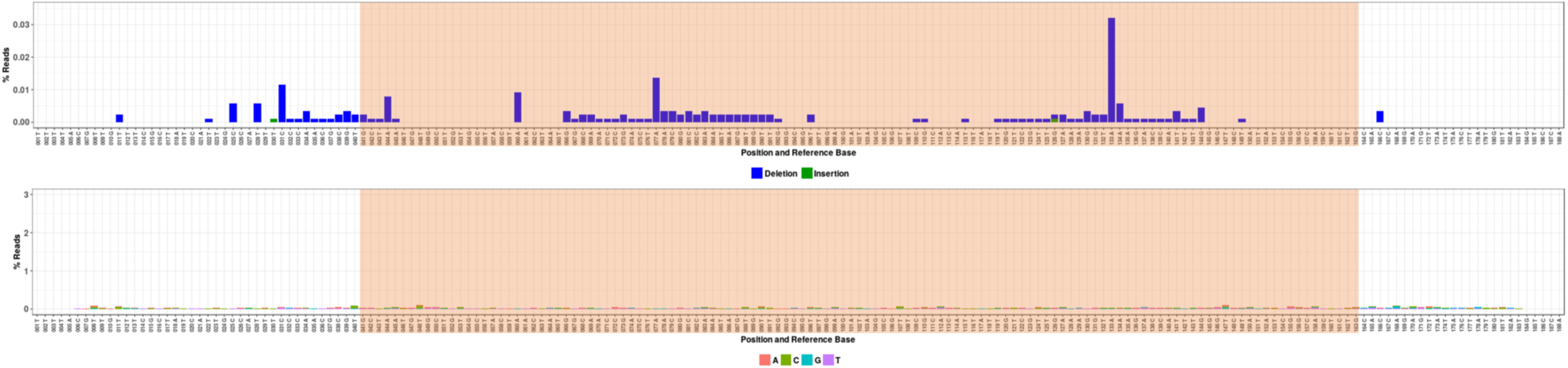
*P. putida* amplicon NGS. Condition 3. Barcode sequence indicated by the orange region. Upper graph: percentage of reads showing a specific deletion/insertion. Lower graph: percentage of reads showing a specific base pair change. ∼86800 read depth.

**Fig. S20.**
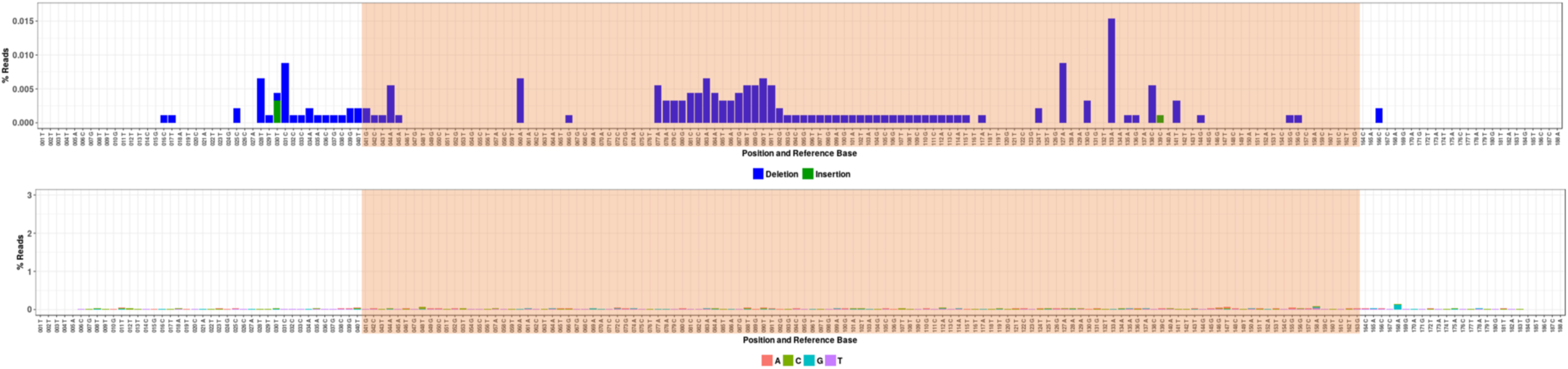
*P. putida* amplicon NGS. Condition 4. Barcode sequence indicated by the orange region. Upper graph: percentage of reads showing a specific deletion/insertion. Lower graph: percentage of reads showing a specific base pair change. ∼90800 read depth.

**Fig. S21.**
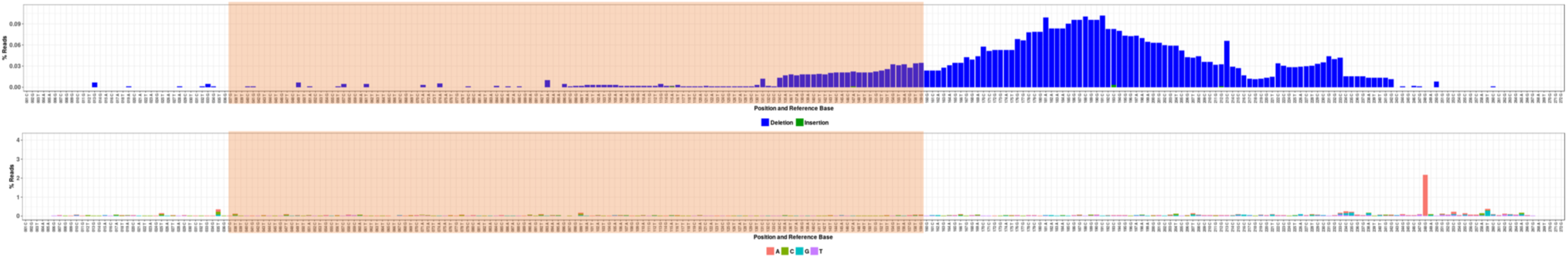
*S. albidoflavus* amplicon NGS. Glycerol control. Barcode sequence indicated by the orange region. Upper graph: percentage of reads showing a specific deletion/insertion. Lower graph: percentage of reads showing a specific base pair change. ∼87400 read depth.

**Fig. S22.**
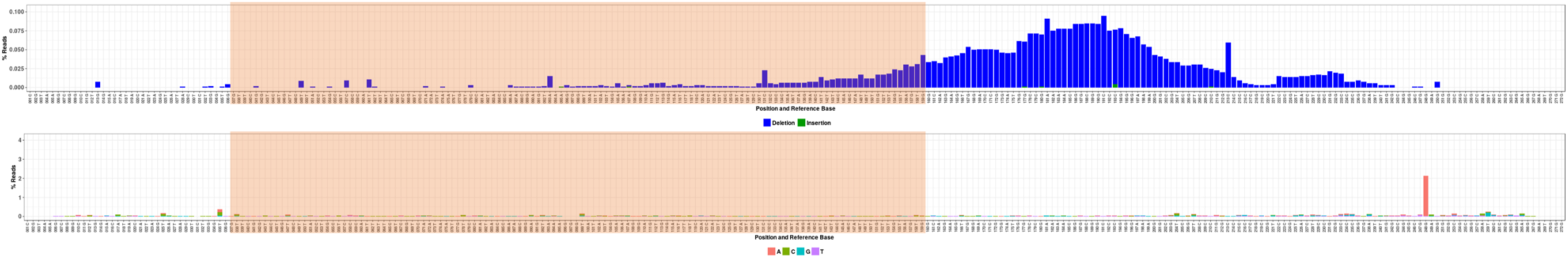
*S. albidoflavus* amplicon NGS. Condition 1. Barcode sequence indicated by the orange region. Upper graph: percentage of reads showing a specific deletion/insertion. Lower graph: percentage of reads showing a specific base pair change. ∼91700 read depth.

**Fig. S23.**
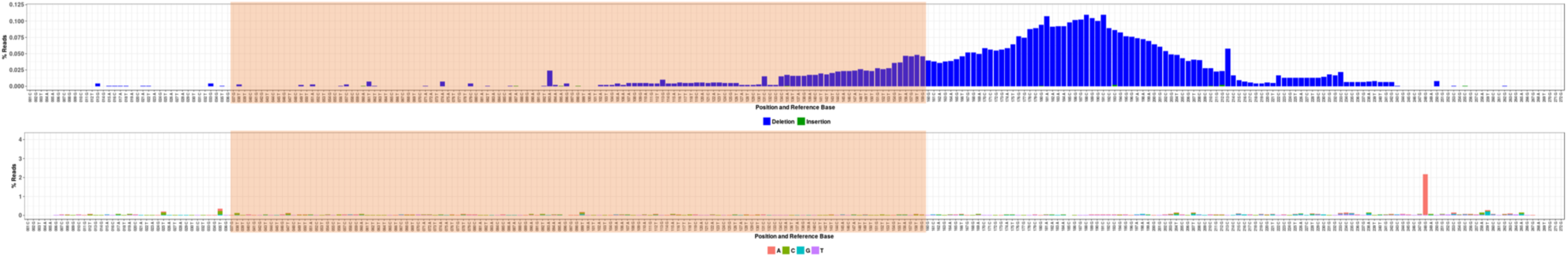
*S. albidoflavus* amplicon NGS. Condition 2. Barcode sequence indicated by the orange region. Upper graph: percentage of reads showing a specific deletion/insertion. Lower graph: percentage of reads showing a specific base pair change. ∼96800 read depth.

**Fig. S24.**
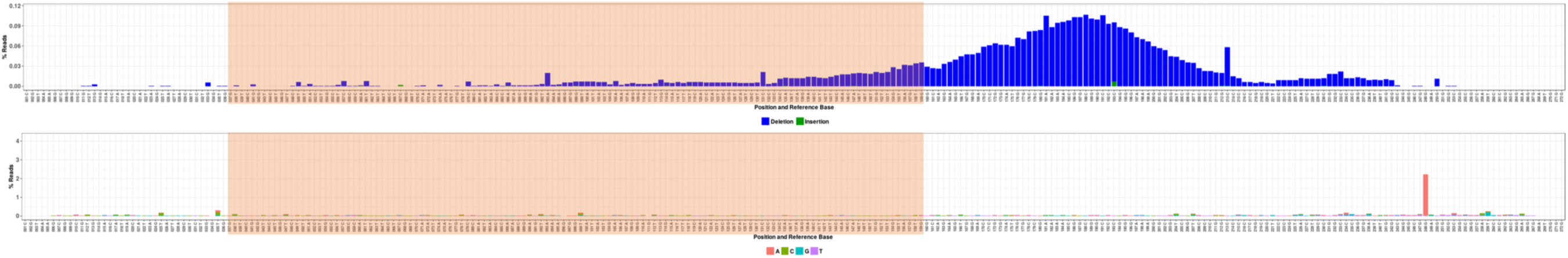
*S. albidoflavus* amplicon NGS. Condition 3. Barcode sequence indicated by the orange region. Upper graph: percentage of reads showing a specific deletion/insertion. Lower graph: percentage of reads showing a specific base pair change. ∼128500 read depth.

**Fig. S25.**
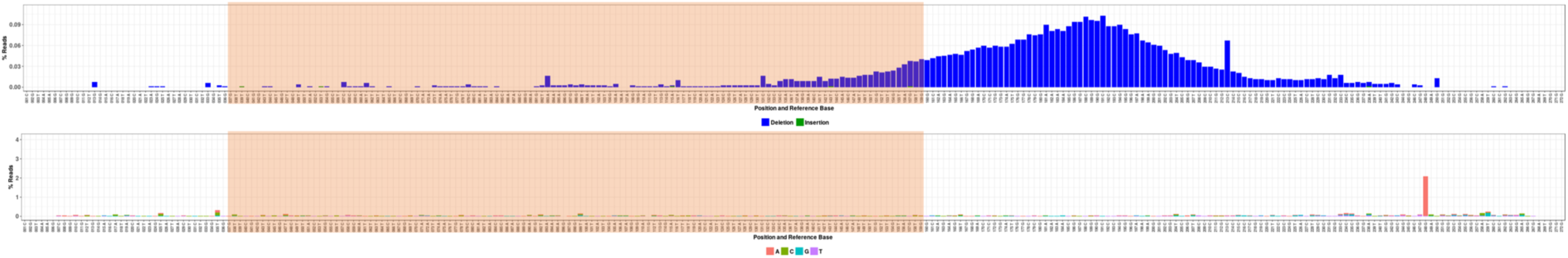
*S. albidoflavus* amplicon NGS. Condition 4. Barcode sequence indicated by the orange region. Upper graph: percentage of reads showing a specific deletion/insertion. Lower graph: percentage of reads showing a specific base pair change. ∼78100 read depth.

**Fig. S26.**
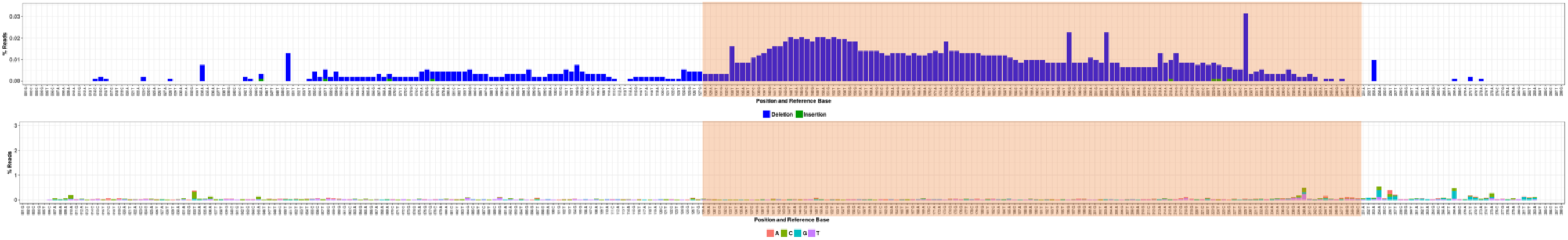
*S. cerevisiae* amplicon NGS. Glycerol control. Barcode sequence indicated by the orange region. Upper graph: percentage of reads showing a specific deletion/insertion. Lower graph: percentage of reads showing a specific base pair change. ∼91500 read depth.

**Fig. S27.**
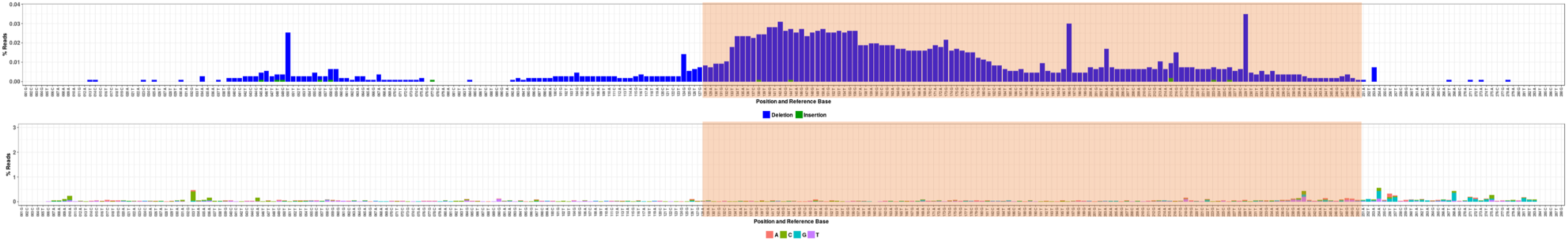
*S. cerevisiae* amplicon NGS. Condition 1. Barcode sequence indicated by the orange region. Upper graph: percentage of reads showing a specific deletion/insertion. Lower graph: percentage of reads showing a specific base pair change. ∼105000 read depth.

**Fig. S28.**
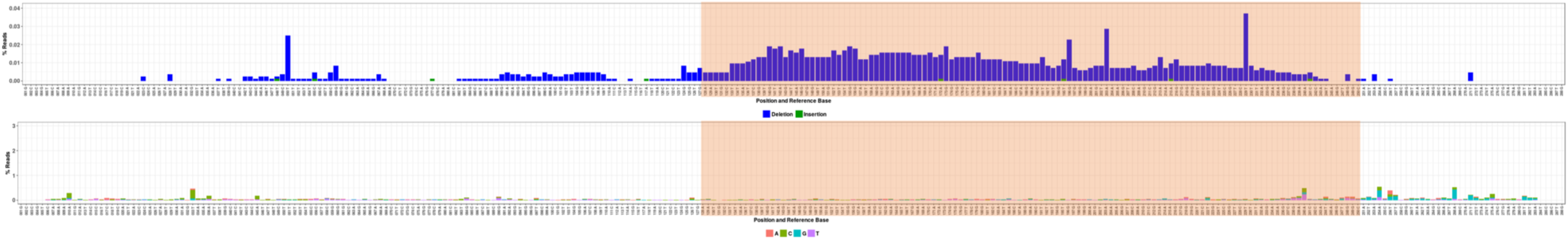
*S. cerevisiae* amplicon NGS. Condition 2. Barcode sequence indicated by the orange region. Upper graph: percentage of reads showing a specific deletion/insertion. Lower graph: percentage of reads showing a specific base pair change. ∼82600 read depth.

**Fig. S29.**
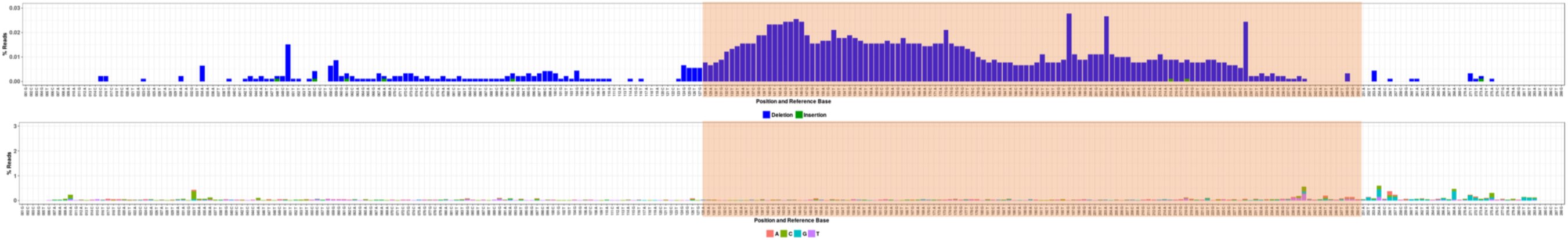
*S. cerevisiae* amplicon NGS. Condition 3. Barcode sequence indicated by the orange region. Upper graph: percentage of reads showing a specific deletion/insertion. Lower graph: percentage of reads showing a specific base pair change. ∼89000 read depth.

**Fig. S30.**
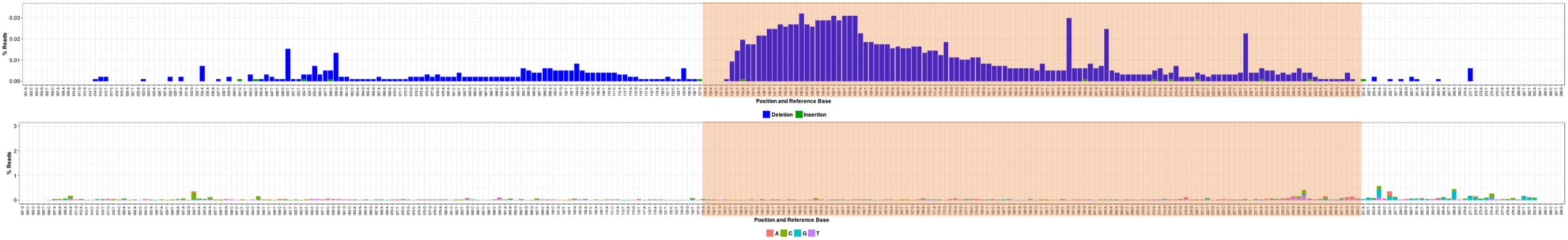
*S. cerevisiae* amplicon NGS. Condition 4. Barcode sequence indicated by the orange region. Upper graph: percentage of reads showing a specific deletion/insertion. Lower graph: percentage of reads showing a specific base pair change. ∼95500 read depth.

**Fig. S31.**
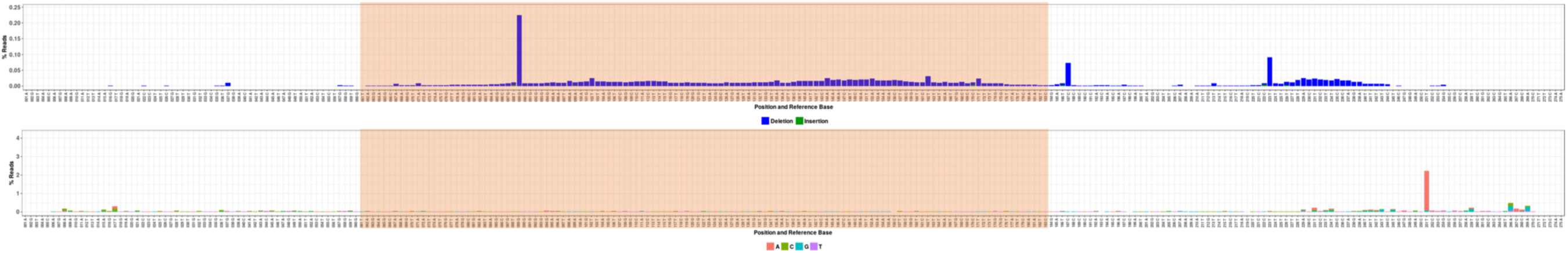
*K. phaffii* amplicon NGS. Glycerol control. Barcode sequence indicated by the orange region. Upper graph: percentage of reads showing a specific deletion/insertion. Lower graph: percentage of reads showing a specific base pair change. ∼73000 read depth.

**Fig. S32.**
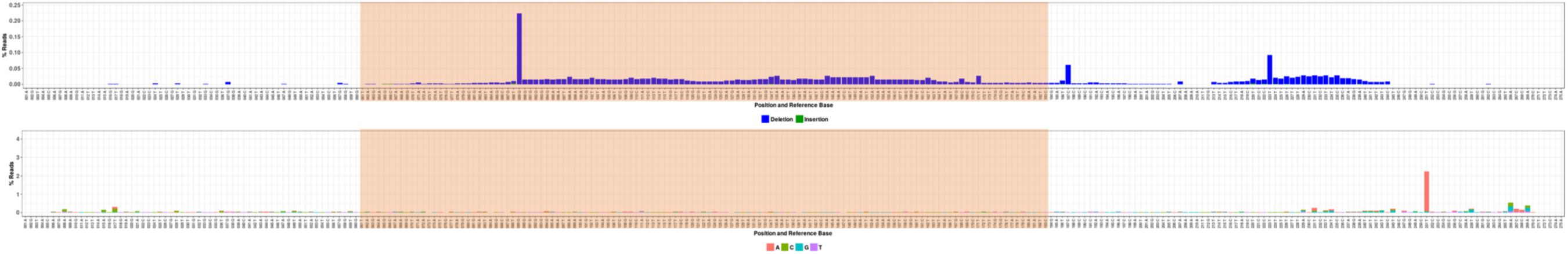
*K. phaffii* amplicon NGS. Condition 1. Barcode sequence indicated by the orange region. Upper graph: percentage of reads showing a specific deletion/insertion. Lower graph: percentage of reads showing a specific base pair change. ∼82900 read depth.

**Fig. S33.**
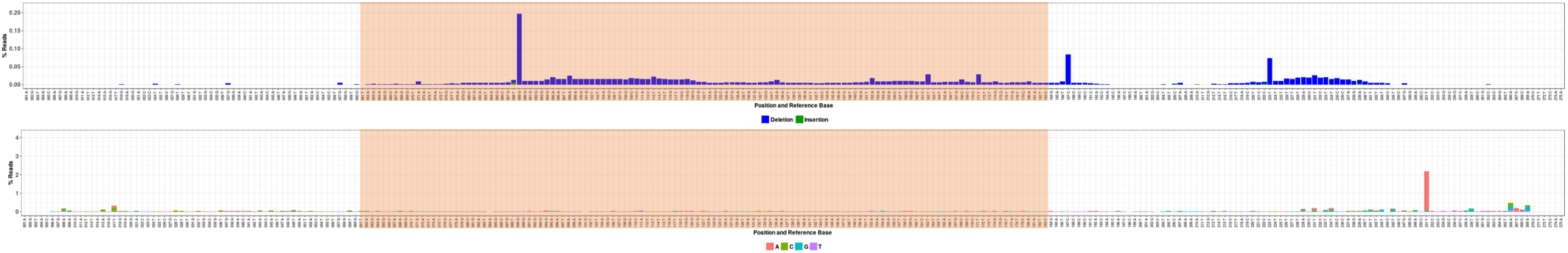
*K. phaffii* amplicon NGS. Condition 2. Barcode sequence indicated by the orange region. Upper graph: percentage of reads showing a specific deletion/insertion. Lower graph: percentage of reads showing a specific base pair change. ∼85400 read depth.

**Fig. S34.**
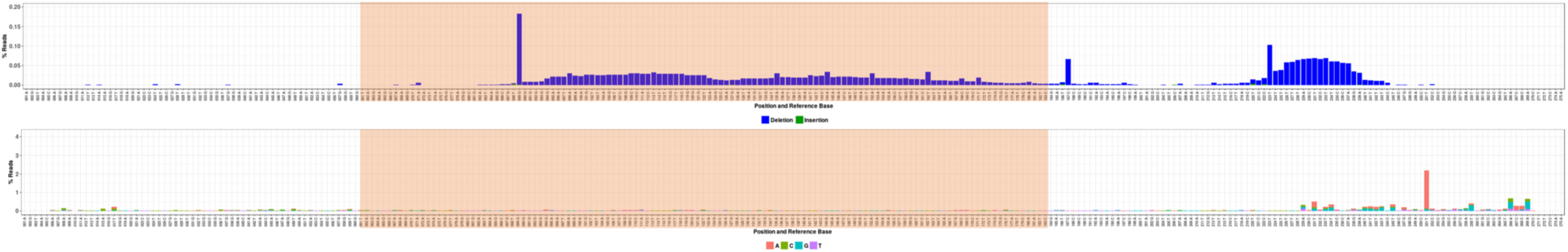
*K. phaffii* amplicon NGS. Condition 3. Barcode sequence indicated by the orange region. Upper graph: percentage of reads showing a specific deletion/insertion. Lower graph: percentage of reads showing a specific base pair change. ∼90600 read depth.

**Fig. S35.**
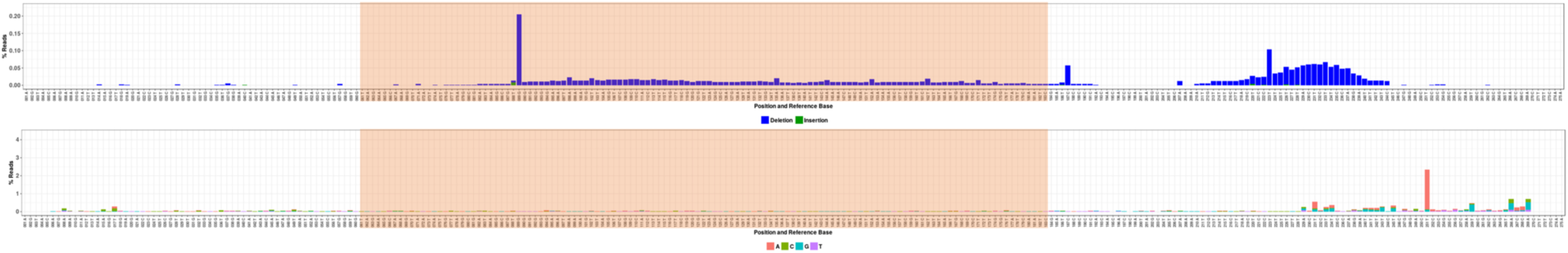
*K. phaffii* amplicon NGS. Condition 4. Barcode sequence indicated by the orange region. Upper graph: percentage of reads showing a specific deletion/insertion. Lower graph: percentage of reads showing a specific base pair change. ∼84600 read depth.

**Fig. S36.**
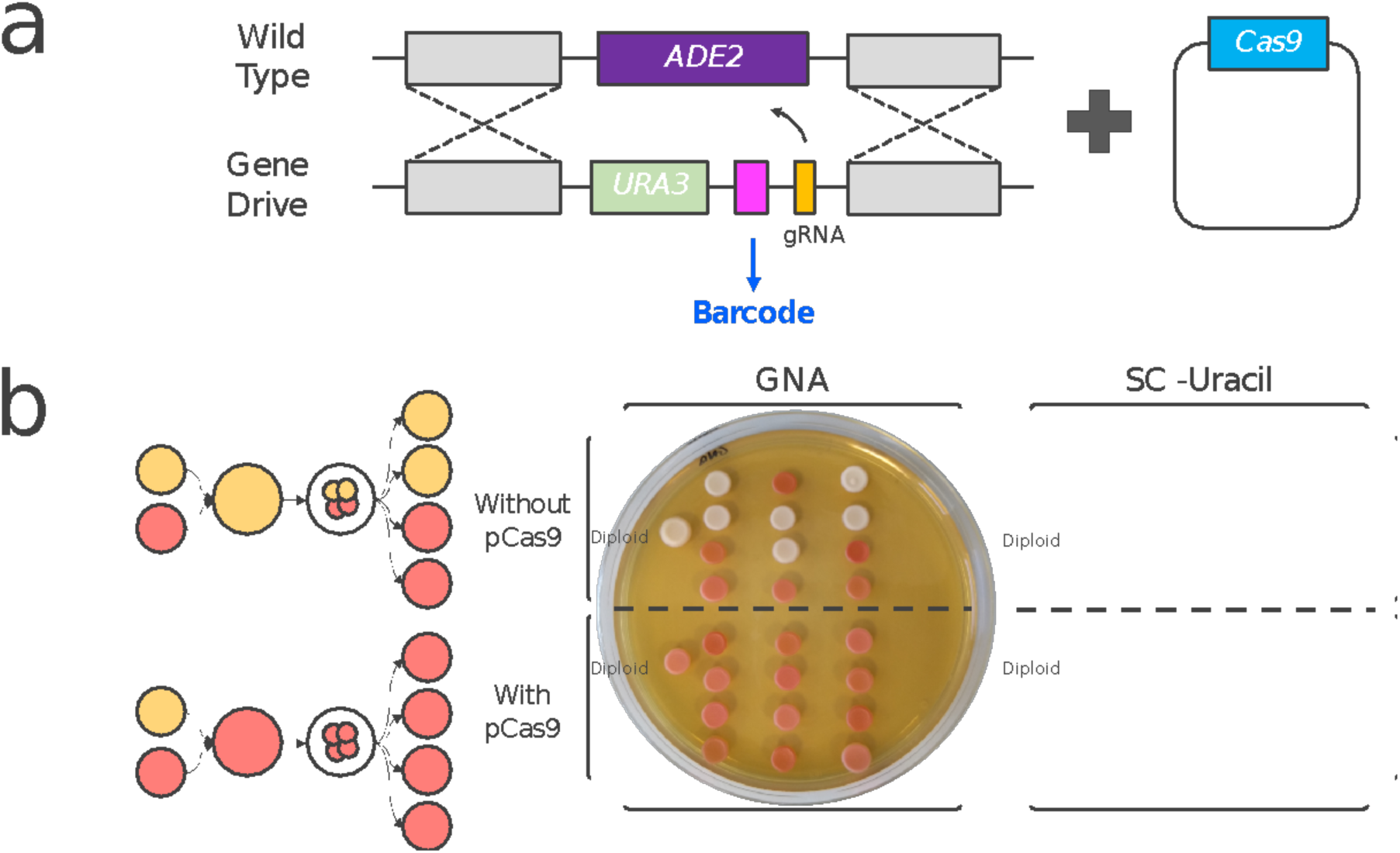
Gene drive experiment in *S. cerevisiae*. a) Graphic representation of the gene drive mechanism used to propagate a barcode. A haploid strain (gene drive) containing a deletion of ADE2 gene substituted by URA3 marker, the barcode sequence and a gRNA targeting ADE2 is mated with a wild type strain of the other sex. In the presence of a plasmid carrying the Cas9 gene, the CRISPR machinery edits the ADE2 gene propagating the desired barcoded phenotype (red colonies, uracil auxotrophic). b) Yeast gene drive experiment results. Haploid cells containing the gene drive cassette were mated with wild type cells in the presence of Cas9. The diploid cells resulting from this were allowed to sporulate and the tetrads were dissected. When Cas9 was not present, the barcoded cassette was inherited by 50% of the final haploid population. When Cas9 was present, the drive allowed the copy of URA3 gene and the barcode DNA in all the spores.

**Table S1.**
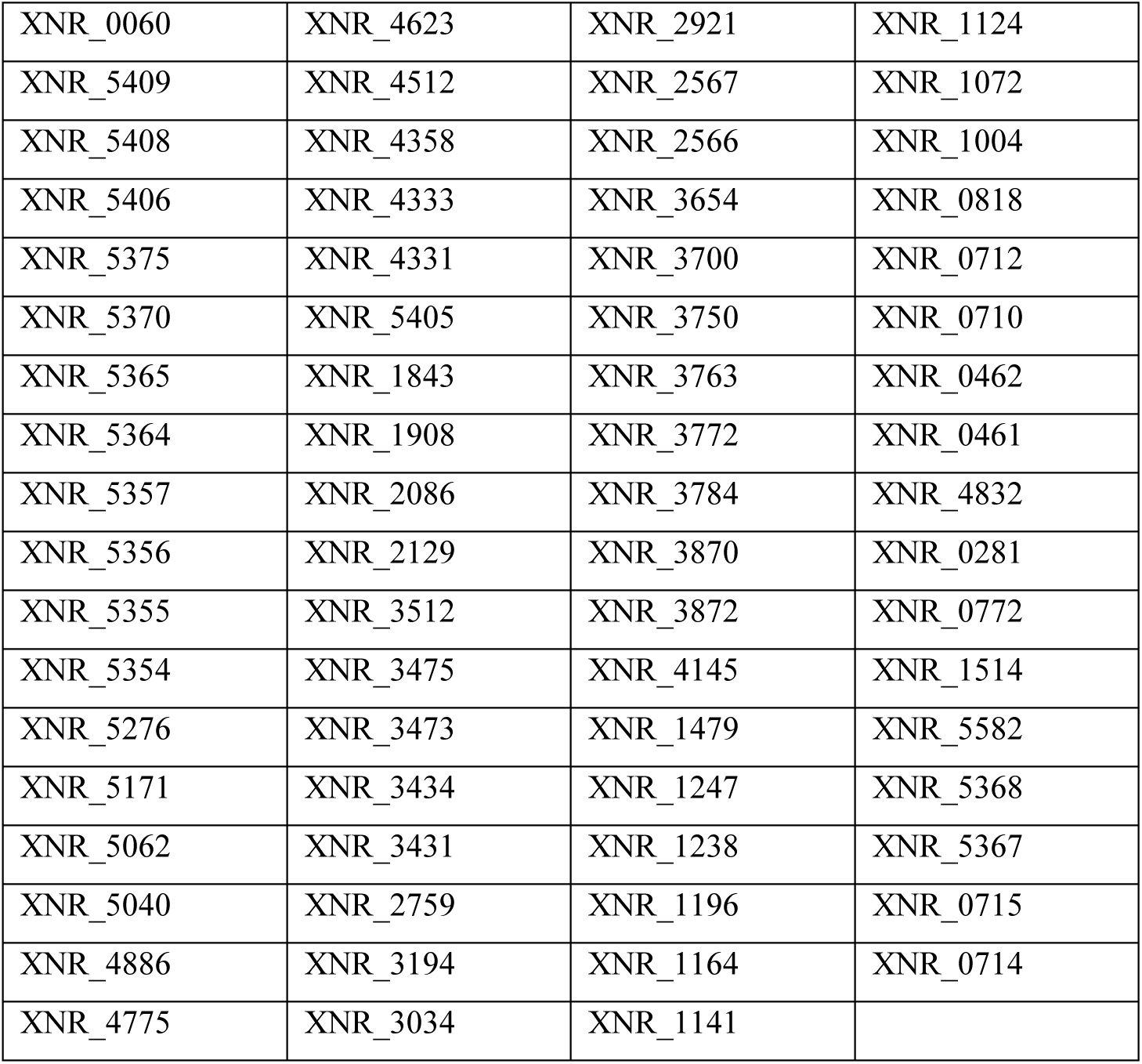
Putative essential gene list of *S. albidoflavus*. Locus tag names from NCBI accession number CP004370.1.

**Table S2.**
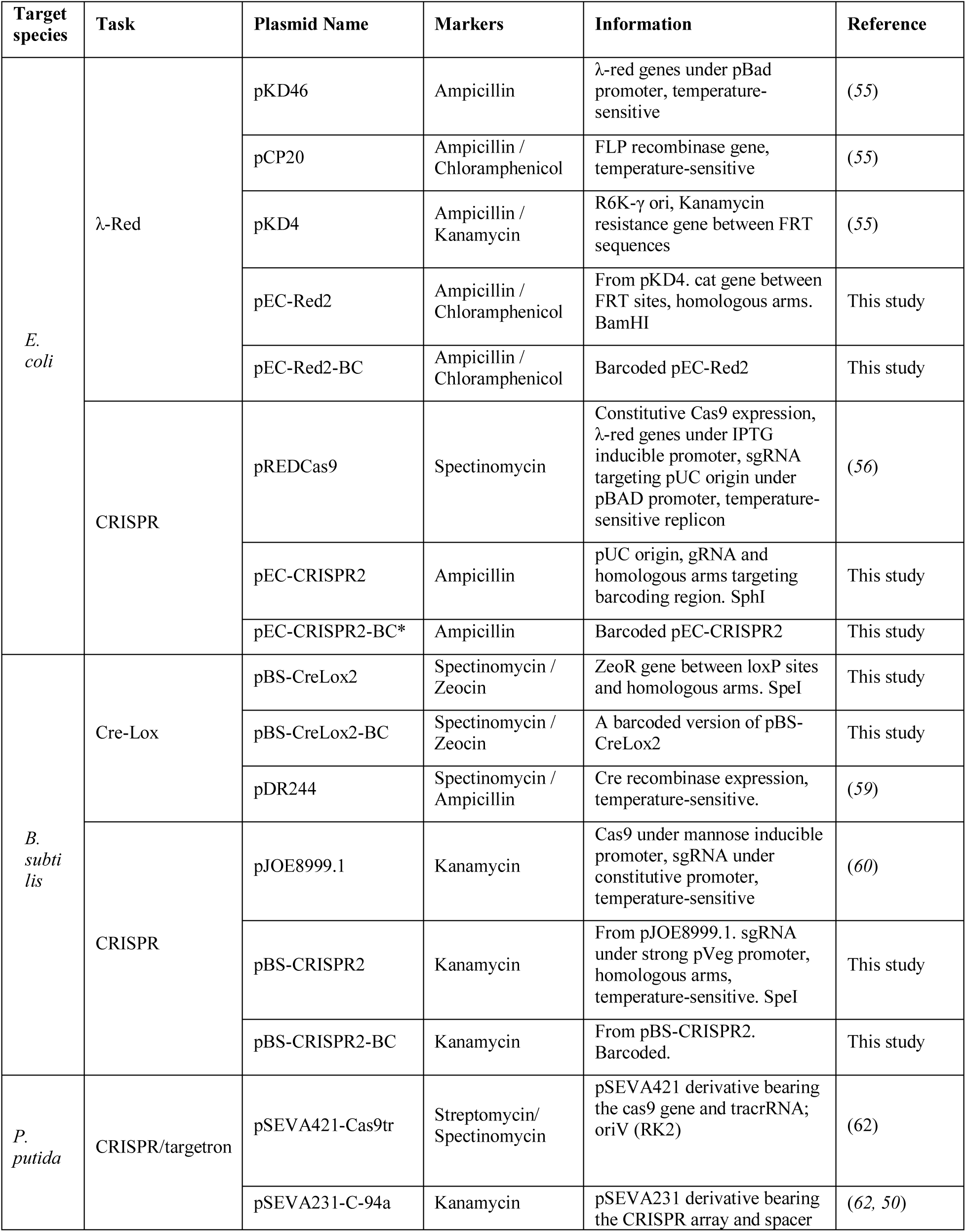
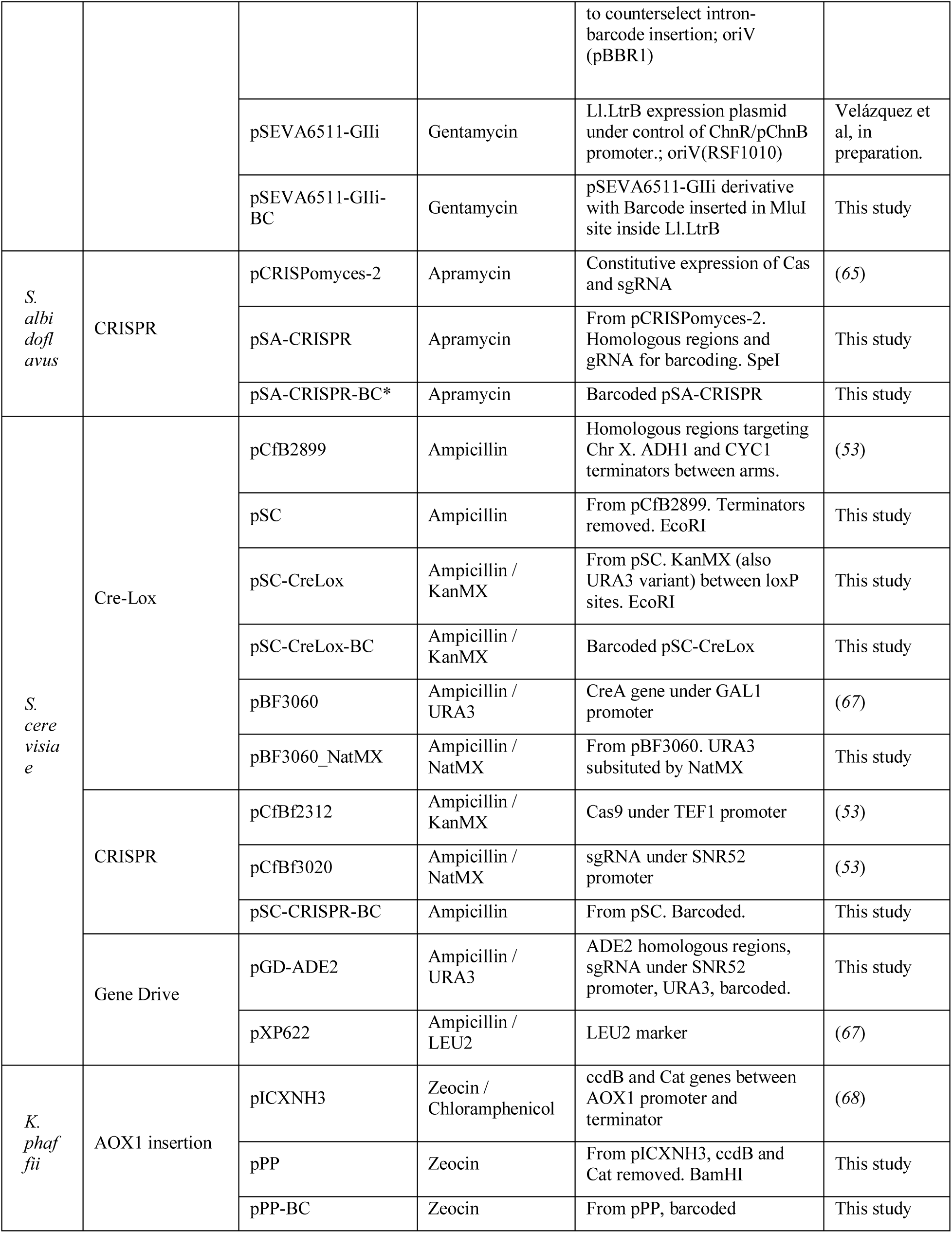
Plasmids used in this study. Restriction sites used to linearize the vectors are shown in italics. (*) Two different plasmids were built with two sgRNA sequences.

**Table S3.**
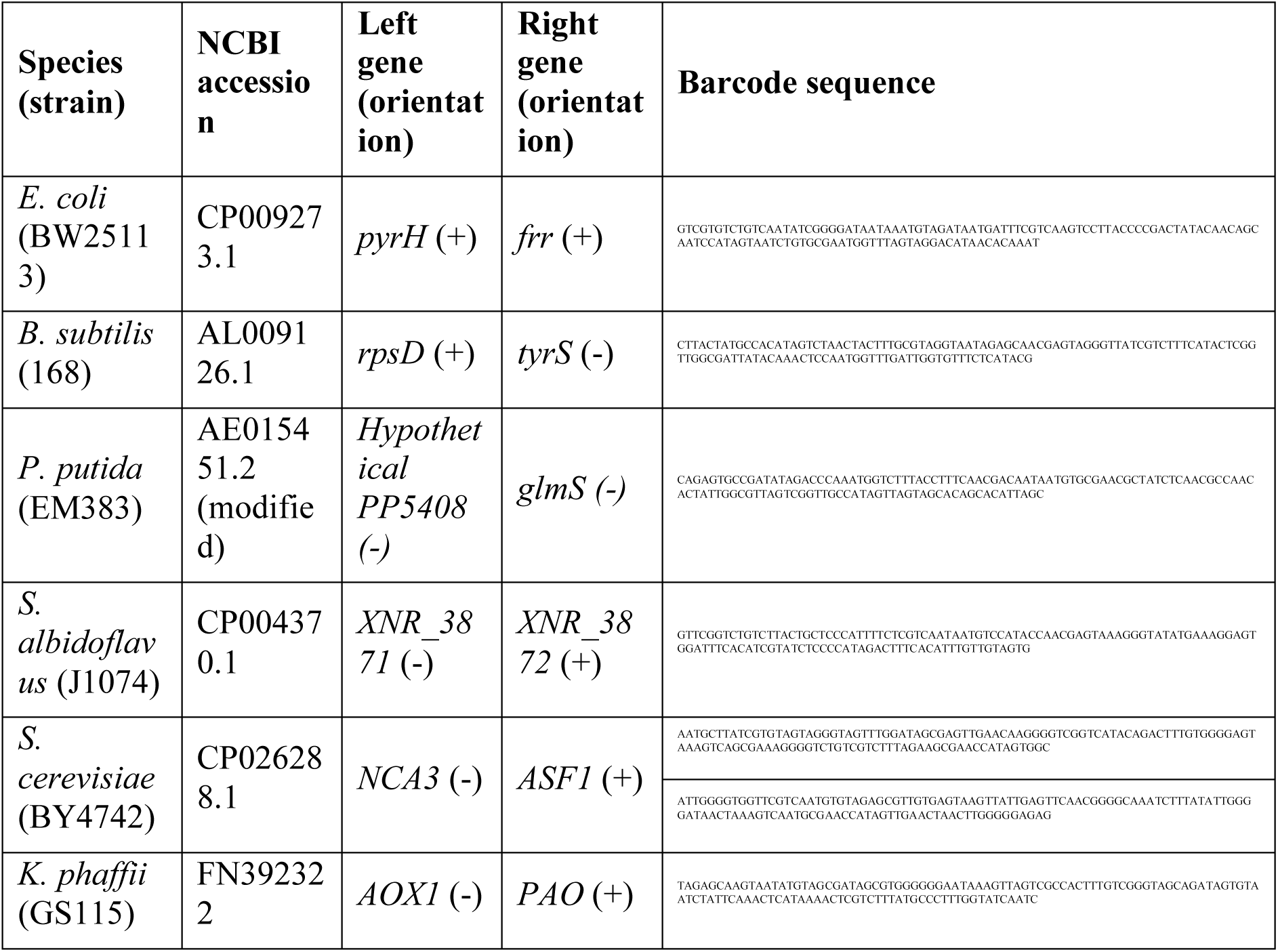
Barcoding site description and barcode sequence information.

**Table S4.**
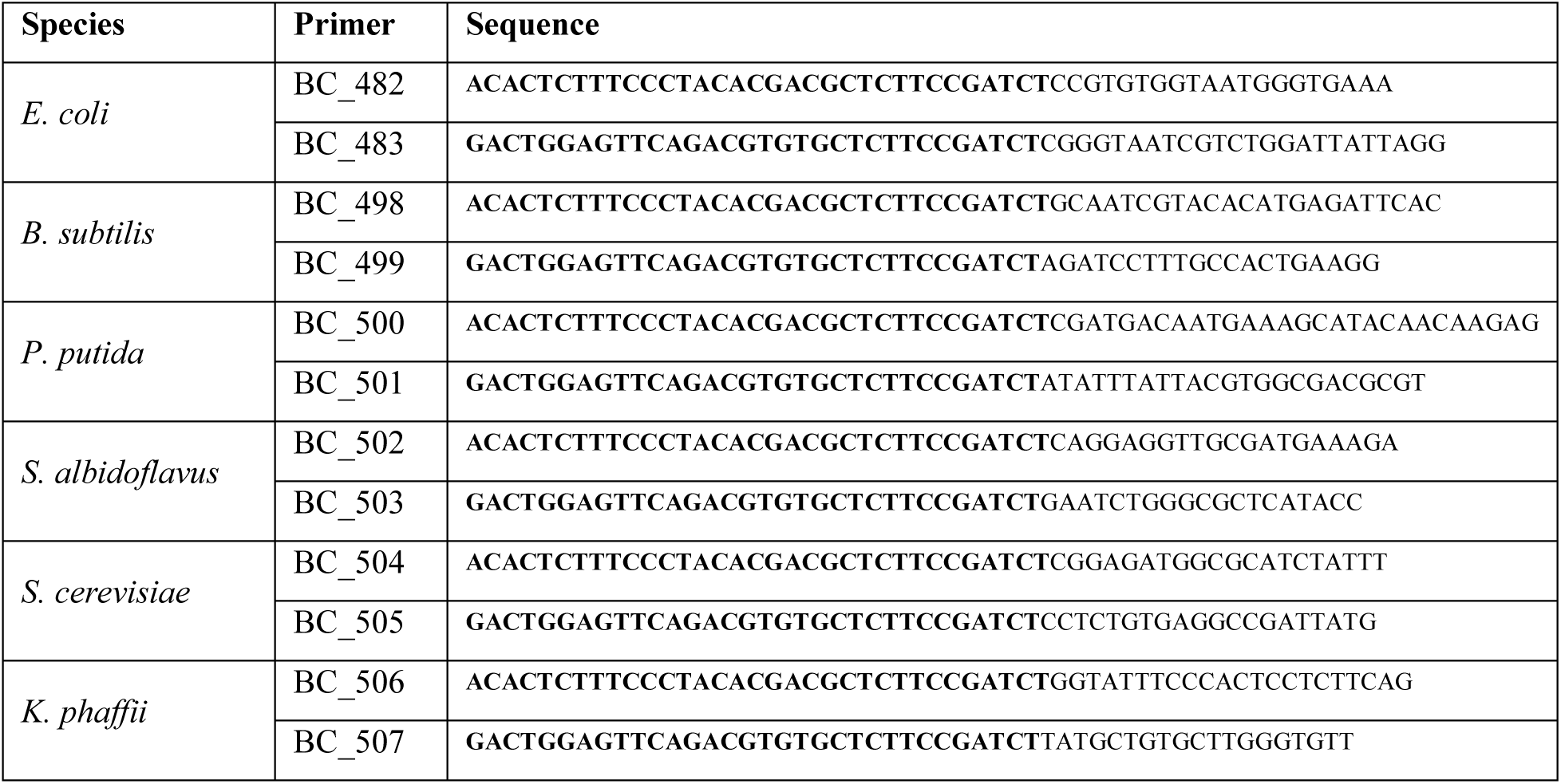
Primers used to sequence the barcoded PCR amplicon. The Illumina adapter sequences appear in bold.

**Table S5.**
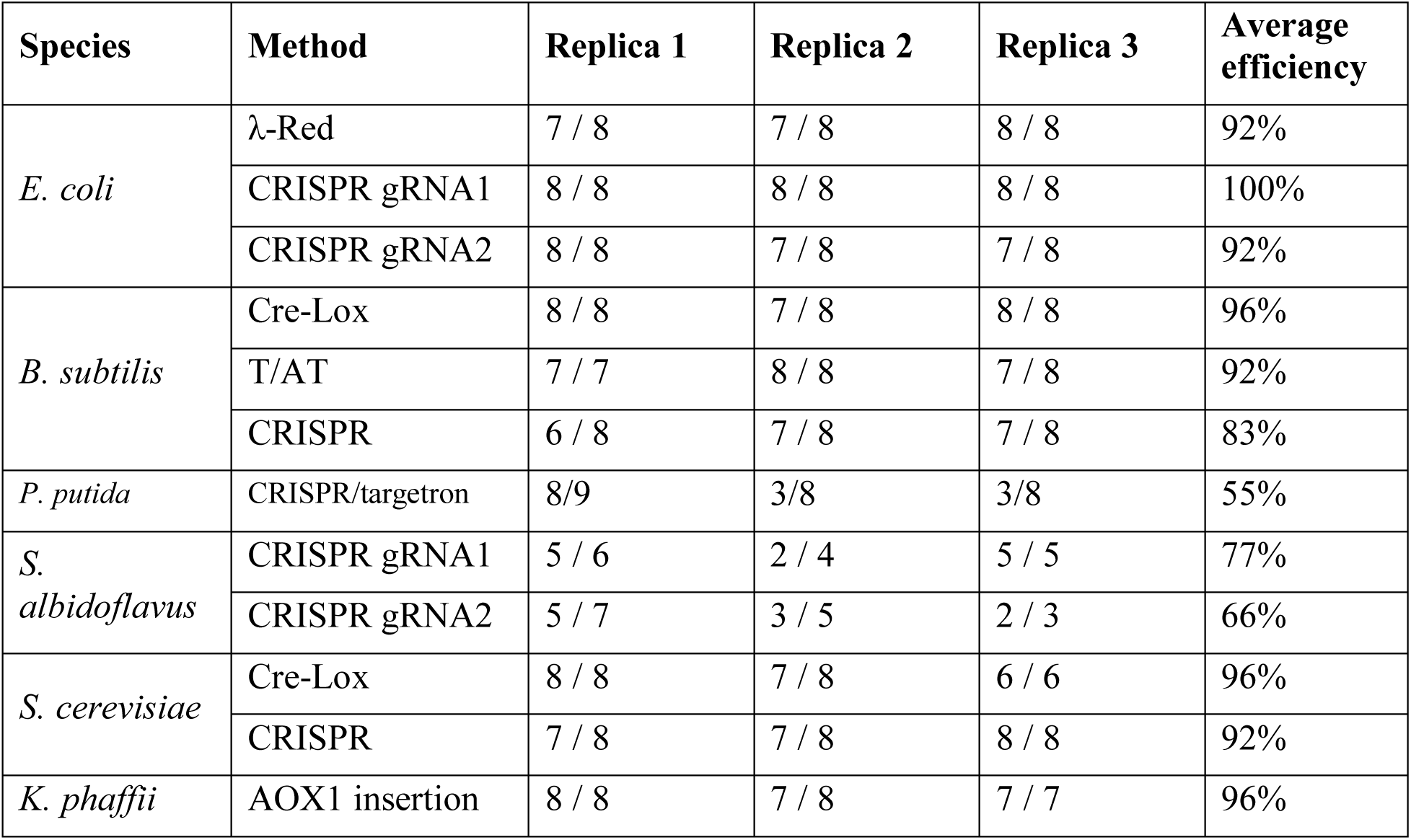
The efficiency of the described barcoding methods used in this article for each species. All the experiments were carried out in three different replicas. After transformation with the PCR product/plasmid colonies were boiled and used as PCR template. The table shows how many positive clones were obtained in all experiments out of the number of tested colonies.

**Table S6.**
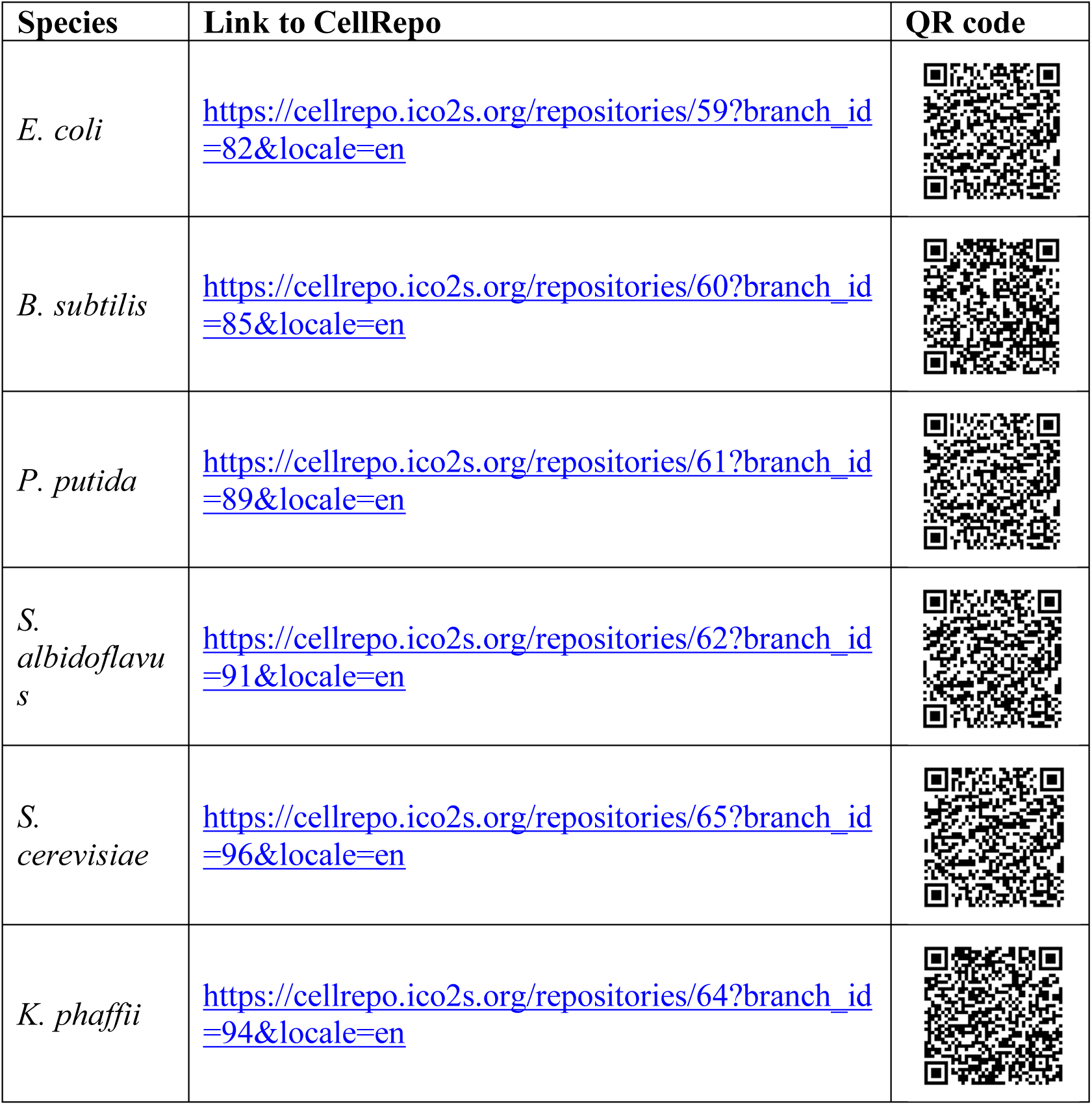
CellRepo links of the barcoded strains in this study.

**Table S7.**
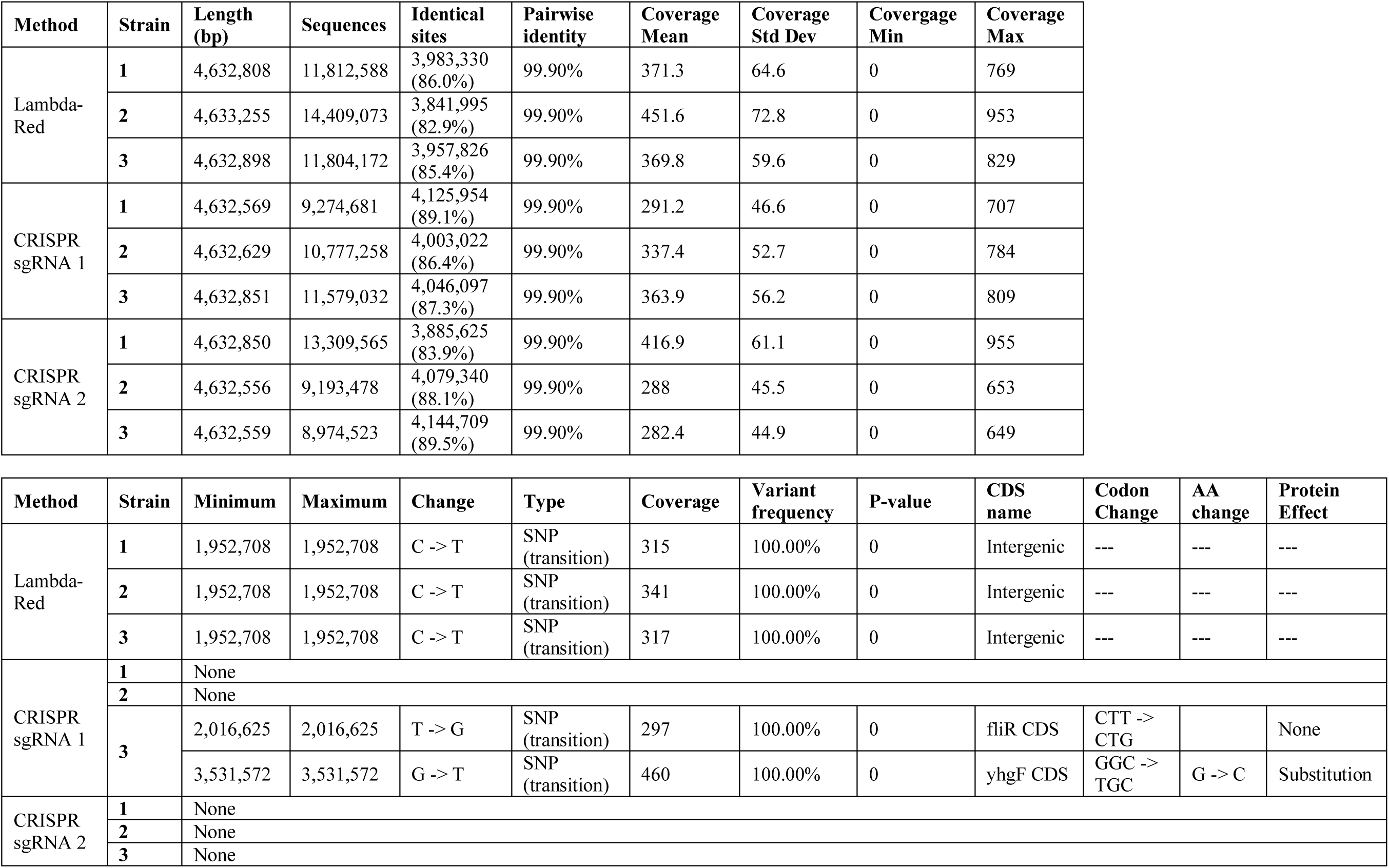
Sequencing statistics (upper table) and mutation analysis (lower table) of E. coli clones barcoded using Lambda-Red and CRISPR (using two different gRNAs) and compared against BW25113 reference genome (CP009273.1).

**Table S8.**
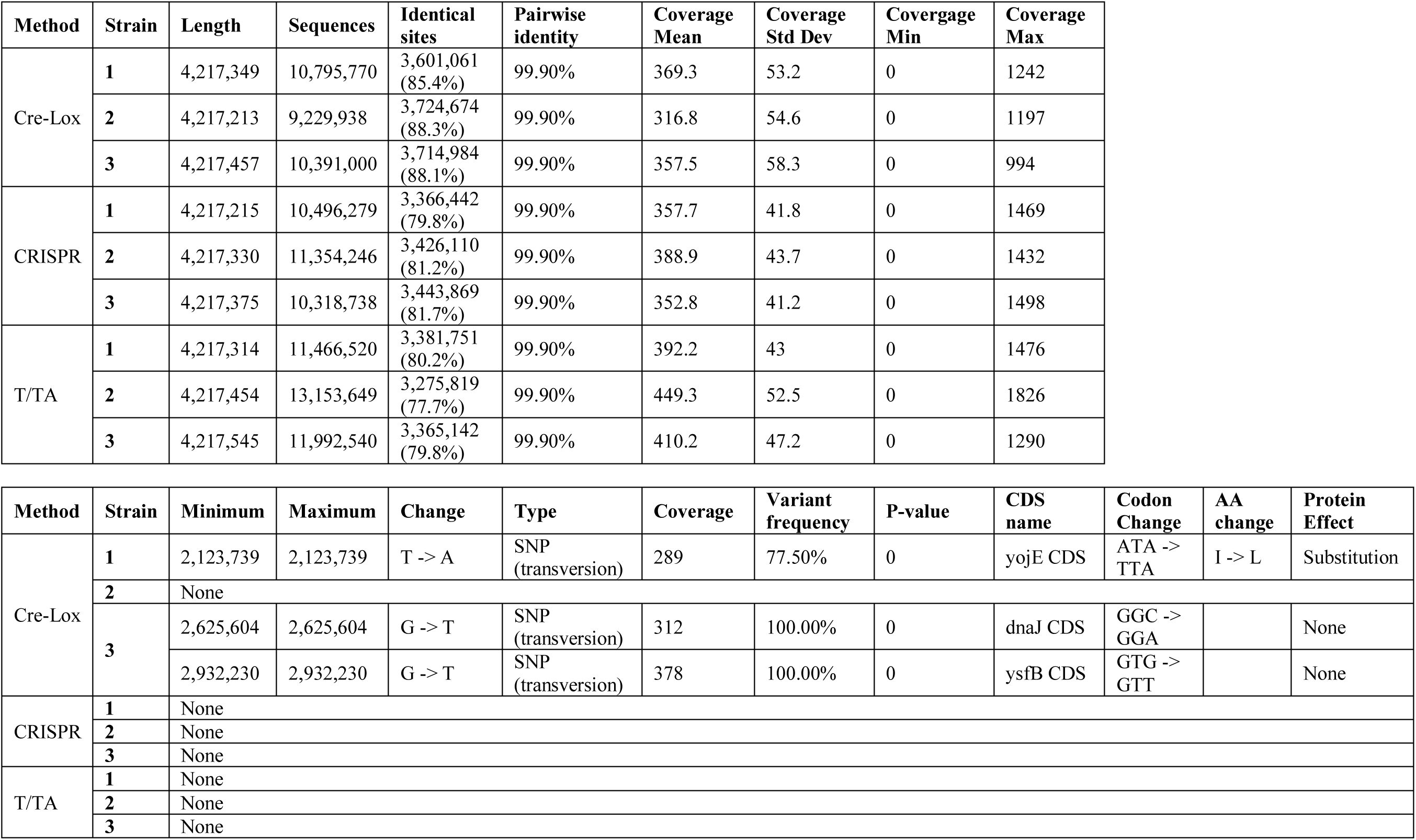
Sequencing statistics (upper table) and mutation analysis (lower table) of *B. subtilis* clones barcoded using three different methods and compared against 168 reference genome (AL009126.1).

**Table S9.**
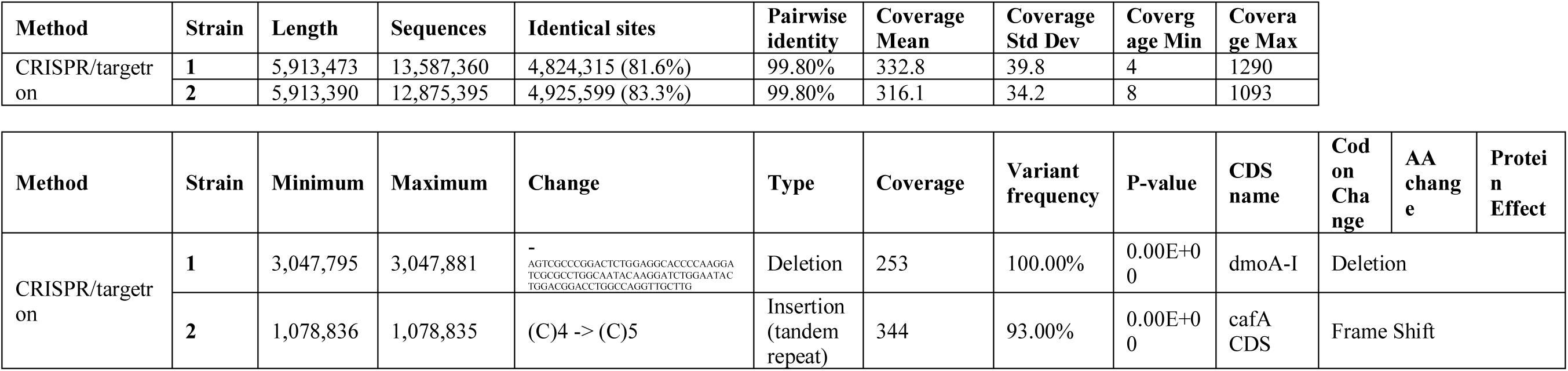
Sequencing statistics (upper table) and mutation analysis (lower table) of P. putida clones barcoded CRISPR/targetron method and compared against EM383 reference genome (a modified version of AE015451.2).

**Table S10.**
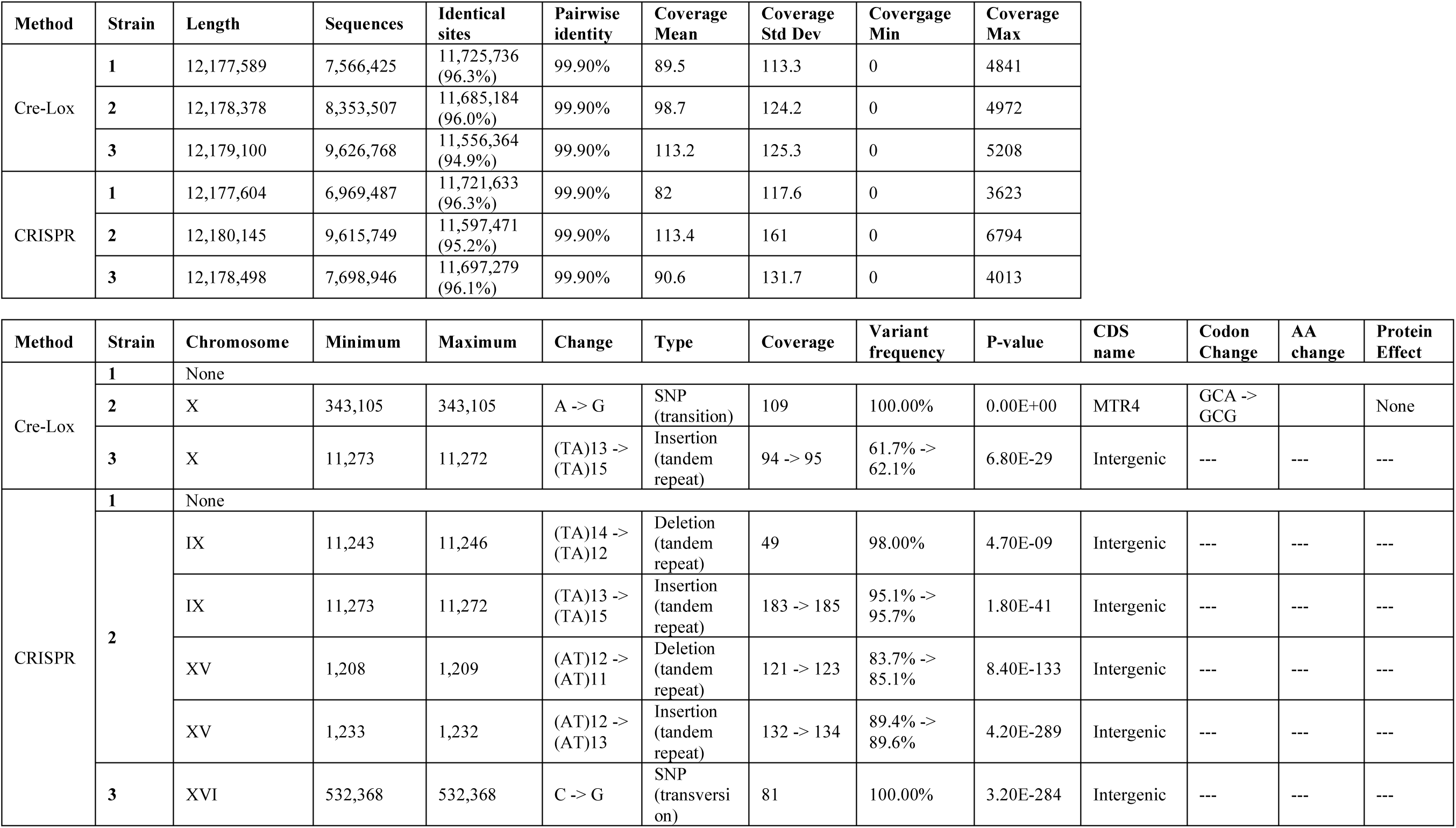
Sequencing statistics (upper table) and mutation analysis (lower table) of S. cerevisiae clones barcoded using two methods and compared against BY4742 reference genome (GenBank assembly accession: GCA_003086655.1).

**Table S11.**
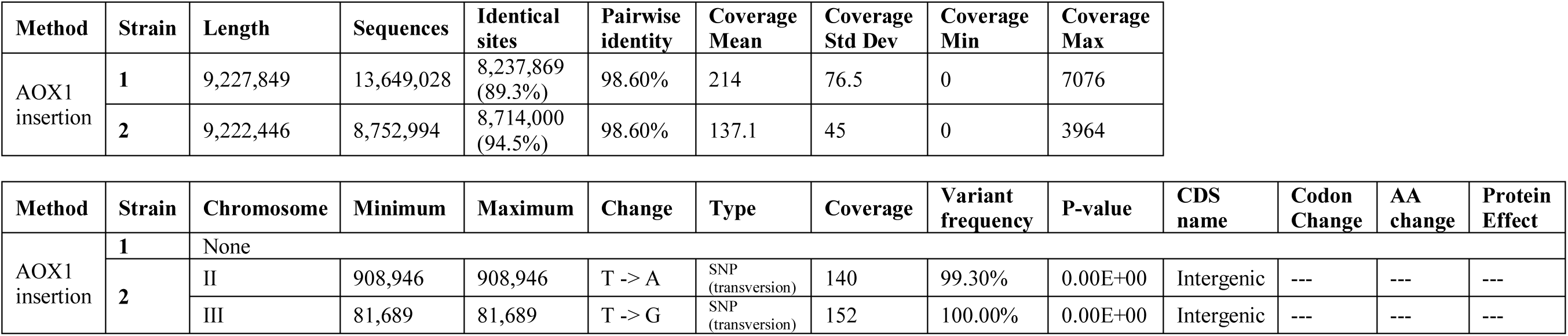
Sequencing statistics (upper table) and mutation analysis (lower table) of *K. phaffii* clones barcoded using AOX1 insertion method and compared against GS115 reference genome (GenBank assembly accession: GCA_000027005.1).

**Table S12.**
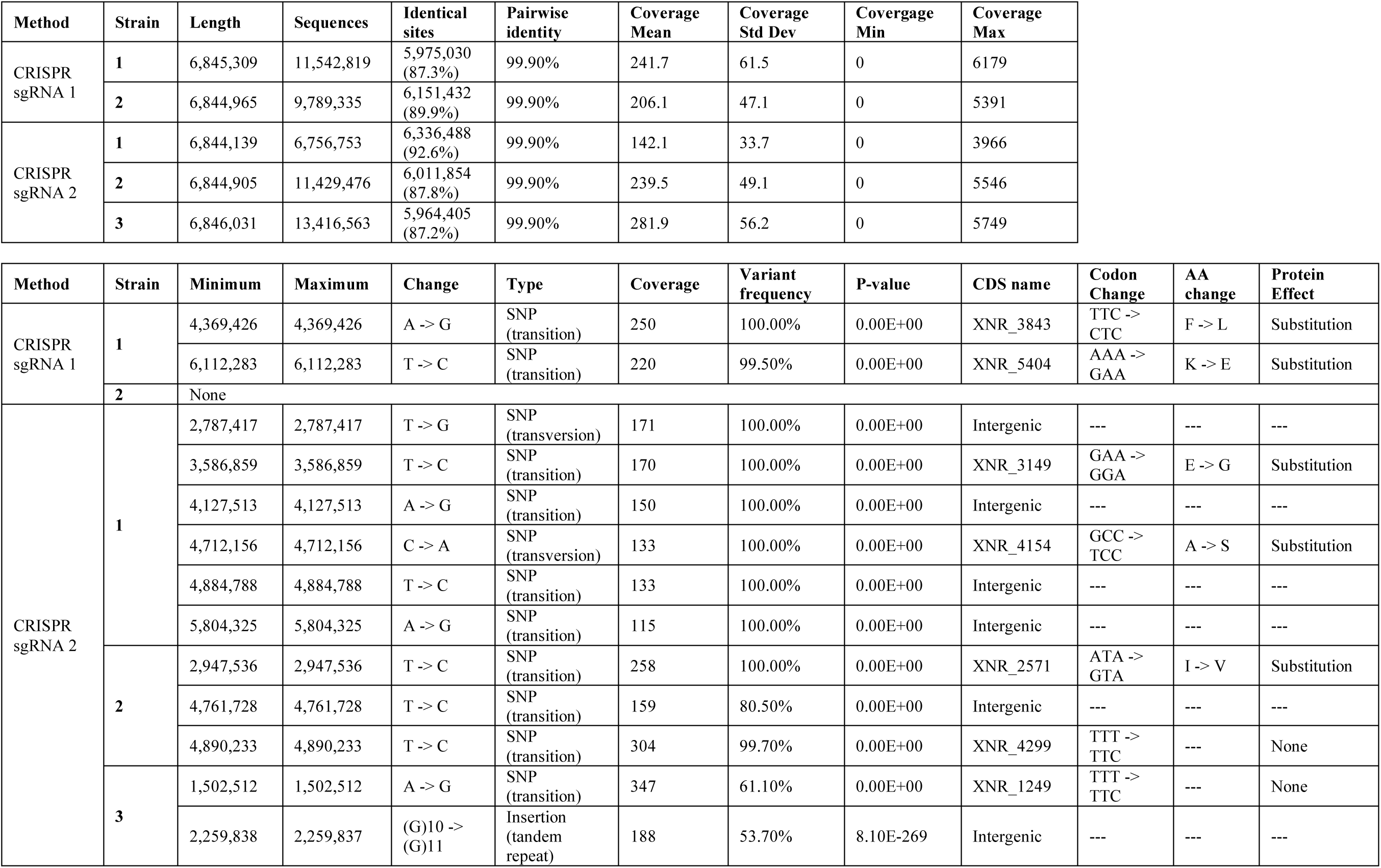
Sequencing statistics (upper table) and mutation analysis (lower table) of S. albidoflavus clones barcoded using gRNAs and compared against J1074 reference genome (CP004370.1).

**Table S13.**
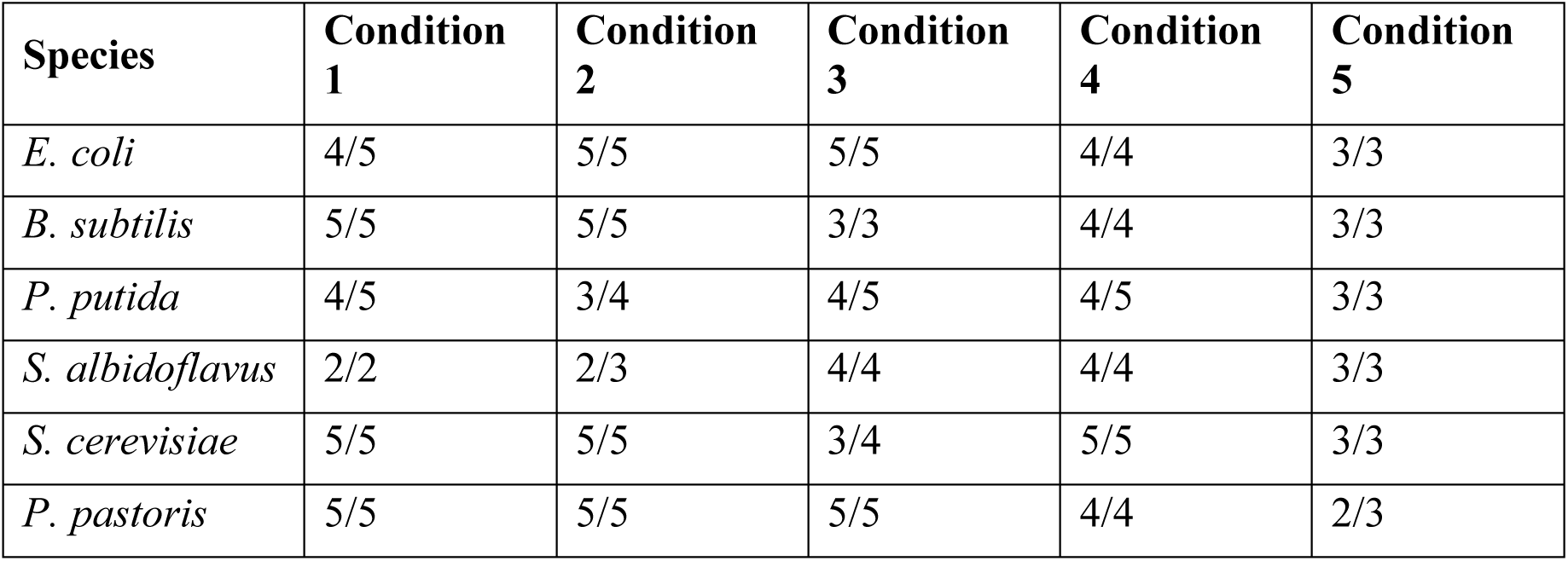
Sanger sequencing results of colonies after the 10 days or passes of each long-term survivability condition. Number of sequences identical to the barcode sequence/number of high-quality Sanger reactions retrieved. Important note: this was a preliminary experiment before NGS amplicon analysis, no replicas were sent for each sequencing reaction. This may explain low-quality reads and point mutations found. The discrepancies were single point mutations in all cases.

## References & Notes

1. A. M. Jinek, K. Chylinski, I. Fonfara, M. Hauer, J. A. Doudna, E. Charpentier, A programmable dual-RNA–guided DNA endonuclease in adaptive bacterial immunity. Science 17, 816–821 (2012).

2. B. R. Mehlenbacher, R. Kolbl, A. Lay, J.A. Dionne, Nanomaterials for in vivo imaging of mechanical forces and electrical fields. Nat Rev Mater 3, 17080 (2018).

3. A. Farhadi, F. Sigmund, G.G. Westmeyer, M.G. Shapiro, Genetically encodable materials for non-invasive biological imaging. Nat. Mater. 10.1038/s41563-020-00883-3 (2021).

4. B. Gilbert, T.C. Tang, W. Ott, B.A. Dorr, W.M. Shaw, G.L. Sun, T.K. Lu, T. Ellis, Living materials with programmable functionalities grown from engineered microbial co- cultures. Nat. Mater. 10.1038/s41563-020-00857-5 (2021).

5. C. V. de Lorenzo, K.L.J. Prather, G.Q. Chen, E. O’Day. C. von Kameke, D.A. Oyarzún, L. Hosta-Rigau, H. Alsafar, C. Cao, W. Ji, H. Okano, R.J. Roberts, M. Ronaghi, K. Yeung, F. Zhang, S.Y. Lee, The power of synthetic biology for bioproduction, remediation and pollution control. EMBO reports 19:e45658, (2018).

6. S.S. Yim, R.M. McBee, A.M. Song, Y. Huang, R.U. Sheth, H.H. Wang, Robust direct digital-to-biological data storage in living cells. Nat Chem Biol 17, 246–253 (2021).

7. A. C. Morrison, R. Lähteenmäki, Public biotech in 2017—the numbers. Nat. Publ. Gr. 36, 576–584 (2018).

8. C.A. Voigt, C.A. Synthetic biology 2020–2030: six commercially-available products that are changing our world. Nat Commun 11, 6379 (2020).

9. A. P. Zhou, X.-L. Yang, X.-G. Wang, B. Hu, L. Zhang, W. Zhang, H.-R. Si, Y. Zhu, B. Li, C.-L. Huang, H.-D. Chen, J. Chen, Y. Luo, H. Guo, R.-D. Jiang, M.-Q. Liu, Y. Chen, X.-R. Shen, X. Wang, X.-S. Zheng, K. Zhao, Q.-J. Chen, F. Deng, L.-L. Liu, B. Yan, F.-X. Zhan, Y.-Y. Wang, G.-F. Xiao, Z.-L. Shi, A pneumonia outbreak associated with a new coronavirus of probable bat origin. Nature 579, 270–273 (2020).

10. A.V. Anzalone, P.B. Randolph, J.R. Davis, A.A. Sousa, L.W. Koblan, J.M. Levy, P.J. Chen, C. Wilson, G.A. newby, A. Raguram, D.R. Liu. Search-and-replace genome editing without double-strand breaks or donor DNA. Nature 576, 149–157 (2019).

11. A. J. Blakes, O. Raz, U. Feige, J. Bacardit, P. Widera, T. Ben-Yehezkel, E. Shapiro, N. Krasnogor, ACS Synthetic Biology 3 (8), 529–542 (2014).

12. B. T. Ben Yehezkel, A. Rival, O. Raz, R. Cohen, Z. Marx, M. Camara, J.F. Dubern, B. Koch, S. Heeb, N. Krasnogor, C. Delattre, E. Shapiro, Synthesis and cell-free cloning of DNA libraries using programmable microfluidics, Nucleic Acids Research, 44(4) (2016).

13. C. S. Reardon, CRISPR creates wave of exotic model organisms. Nat. (London, U. K.). 568, 441–442 (2019).

14. D. V. de Lorenzo, N. Krasnogor, M. Schmidt, For the sake of the Bioeconomy: define what a Synthetic Biology chassis is!. New Biotechnology 60, (2021).

15. Identity crisis. Nature 457, 935–936 (2009).

16. American Type Culture Collection Standards Development Organization Workgroup ASN-0002. Cell line misidentification: the beginning of the end. Nat Rev Cancer 10, 441–448 (2010).

17. A. J. Masters. End the scandal of false cell lines. Nature 492, 186 (2012).

18. Challenges in irreproducible research. Nature, Special online collection, https://www.nature.com/collections/prbfkwmwvz/ (2018).

19. M.R. Mehra, F. Ruschitzka, A.N. Patel. Retraction: Hydroxychloroquine or chloroquine with or without a macrolide for treatment of COVID-19: a multinational registry analysis. The Lancet, 395(10240), P1820 (2020).

20. A. H. Akin, K.M. Rose, D.A. Scheufele, M. Simis-Wilkinson, D. Brossards, M.A. Xenos, E.A. Corley, Mapping the landscape of public attitudes on Synthetic Biology. BioScience 67 (3), (2017).

21. B. P. Sturgis, N. Allum, Science in society: re-evaluating the deficit model of public attitudes, Public understanding of science, 13 (1), 10.1177/0963662504042690 (2004).

22. C. S. Ho, D.A. Scheufele, E.A. Corley, Making sense of policy choices: understanding the roles of value predispositions, mass media, and cognitive processing in public attitudes toward nanotechnology. Journal of Nanoparticle Research 12 (2010).

23. D. S. Jasanoff, The ‘science wars’ and American politics,” in Between understanding and trust: the public, science and technology, M. Dierkes, C. von Grote, Eds., Routledge, (2000).

24. E. S. Yearley, What does science mean in the ‘public understanding of science’?” in Between Understanding and Trust: The Public, Science and Technology, M. Dierkes, C. von Grote, Eds., Routledge, (2000).

25. F. D. Bhattachary, J. Pascall Calitz, A. Hunter, Synthetic Biology Dialogue, TNS-BMRB research agency, https://royalsociety.org/-/media/about-us/international/g-science-statements/2019-g7-declaration-science-and-trust.pdf?la=en-GB&hash=32D575A44FA381AB16B9ADF762FA99FB (2010).

26. Summit of the G7 science academies, Science and Trust, Executive summary and recommendations. https://royalsociety.org/-/media/about-us/international/g-science-statements/2019-g7-declaration-science-and-trust.pdf?la=en-GB&hash=32D575A44FA381AB16B9ADF762FA99FB (2019).

27. A. D. Brossard, D., C.M. Nisbet, Deference to scientific authority among a low information public: understanding U.S. opinion on agricultural biotechnology. International Journal of Public Opinion Research, 19 (1), (2006).

28. B. M. Baker, 1,500 scientists lift the lid on reproducibility. Nature, 533(7604) (2016).

29. C. F. Fidler, J. Wilcox, Reproducibility of scientific results in The Stanford Encyclopedia of Philosophy, E.N. Zalta Ed., https://plato.stanford.edu/archives/win2018/entries/scientific-reproducibility/ (2018).

30. A. Nosek, G. Alter, G. C. Banks, D. Borsboom, S. D. Bowman, S. J. Breckler, S. Buck, C. D. Chambers, G. Chin, G. Christensen, M. Contestabile, A. Dafoe, E. Eich, J. Freese, R. Glennerster, D. Goroff, D. P. Green, B. Hesse, M. Humphreys, J. Ishiyama, D. Karlan, A. Kraut, A. Lupia, P. Mabry, T. Madon, N. Malhotra, E. Mayo-Wilson, M. McNutt, E. Miguel, E. Levy Paluck, U. Simonsohn, C. Soderberg, B. A. Spellman, J. Turitto, G. VandenBos, S. Vazire, E. J. Wagenmakers, R. Wilson, T. Yarkoni B.A. Nosek, G. Alter, G.C. Banks, Promoting an open research culture. Science 348, 1422–1425 (2015).

31. Research integrity: a landscape Study. Commissioned by UK Research and Innovation (UKRI), Vitae in partnership with the UK Research Integrity Office (UKRIO) and the UK Reproducibility Network (UKRN), Vitae (2020).

32. What Researchers Think About the Culture They Work In. Wellcome Trust (2020).

33. M.D. Wilkinson, M. Dumontier, I.J Aalbersberg, G. Appleton, M. Axton, A. Baak, N. Blomberg, J.W. Boiten, L. Bonino da Silva Santos, P.E. Bourne, J. Bouwman, A.J. Brookes, T. Clark, M. Crosas, I. Dillo, O. Dumon, S. Edmunds, C.T. Evelo, R. Finkers, Gonzalez-Beltran, A.J.G. Gray, P. Groth, C. Goble, J.S. Grethe, J. Heringa, P.A.C ’t Hoen, R. Hooft, T. Kuhn, R. Kok, J. Kok, S.J. Lusher, M.E. Martone, A. Mons, A.L. Packer, B. Persson, P. Rocca-Serra, M. Roos, R. van Schaik, S.A. Sansone, E. Schultes, T. Sengstag, T. Slater, G. Strawn, M.A. Swertz, M. Thompson, J. van der Lei, E. van Mulligen, J. Velterop, a. Waagmeester, P. Wittenburg, K. Wolstencroft, J. Zhao, B. Mons, The FAIR guiding principles for scientific data management and stewardship. Sci Data 3, 160018 (2016).

34. The Concordat on Open Research Data. United Kingdom Research and Innovation (2016).

35. A. J. Tellechea-Luzardo, C. Winterhalter, P. Widera, J. Kozyra, V. de Lorenzo, N. Krasnogor, Linking engineered cells to their digital twins: a version control system for strain engineering. ACS Synth. Biol. 9, 536–545 (2020).

36. B. S. More, V. Bampidis, D. Benford, C. Bragard, T. Halldorsson, A. Hernández-Jerez, S. Hougaard Bennekou, K. Koutsoumanis, K. Machera, H. Naegeli, S.S. Nielsen, J. Schlatter, D. Schrenk, V. Silano, D. Turck, M. Younes, B. Glandorf, L. Herman, C. Tebbe, J. Vlak, J. Aguilera, R. Schoonjans, P.S. Cocconcelli, Evaluation of existing guidelines for their adequacy for the microbial characterisation and environmental risk assessment of microorganisms obtained through synthetic biology. EFSA Journal 18(18):6263 (2020).

37. C. H. Baig, P. Fontanarrosa, V. Kulkarni, J.A. McLaughlin, P. Vaidyanathan, B. Bartley, J. Beal, M. Crowther, T.E. Gorochowski, R. Grünberg, G. Misirli, J. Scott-Brown, E. Oberortner, A. Wipat, C.J. Myers, Synthetic biology open language (SBOL) version 3.0.0. J Integr Bioinform. 25 (2020).

38. D. M. Baker, How quality control could save your science. Nature 529, 456–458 (2016).

39. M.C. Murphy, A.F. Mejia, J. Mejia, X. Yan, S. Cheryan, N. Dasgupta, M. Destin, S.A. Fryberg, J.A. Garcia, E.L. Haines, J.M. Harackiewicz, A. Ledgerwood, C.A. Moss-Racusin, L.E. Park, S.P. Perry, K.A. Ratliff, A. Rattan, D.T. Sanchez, K. Savani, D. Sekaquaptewa, J.L. Smith, V. Jones Taylor, D.B. Thoman, D.A. Wout, P.L. Mabry, S. Ressl, A.B. Diekman, F. Pestilli, Open science, communal culture, and women’s participation in the movement to improve science. Proc. Natl. Acad. Sci. U.S.A 117(39), (2020).

40. A. T. Baba, T. Ara, M. Hasegawa, Y. Takai, Y. Okumura, M. Baba, K. A. Datsenko, M. Tomita, B. L. Wanner, H. Mori, H. Mori, Construction of Escherichia coli K-12 in-frame, single gene knockout mutants: the Keio collection. Mol. Syst. Biol. (2006), doi:10.1038/msb4100050.

41. B. R. H. Michna, F. M. Commichau, D. Tö, C. P. Zschiedrich, J. Rg, S. Lke, SubtiWiki-a database for the model organism Bacillus subtilis that links pathway, interaction and expression information. Nucleic Acids Res. 42 (2014), doi:10.1093/nar/gkt1002.

42. C. P. D. Karp, R. Billington, R. Caspi, C. A. Fulcher, M. Latendresse, A. Kothari, I. M. Keseler, M. Krummenacker, P. E. Midford, Q. Ong, W. K. Ong, S. M. Paley, P. Subhraveti, The BioCyc collection of microbial genomes and metabolic pathways. Brief. Bioinform. 20, 1085–1093 (2017).

43. D. V. Solovyev, A. Salamov. Automatic Annotation of Microbial Genomes and Metagenomic Sequences. Metagenomics and its Applications in Agriculture (2013).

44. E. A. Lesnik, R. Sampath, H. B. Levene, T. J. Henderson, J. A. Mcneil, D. J. Ecker, Prediction of rho-independent transcriptional terminators in Escherichia coli. Nucleic Acids Res. 29, 3583–3594 (2001).

45. Molina-Henares MA, de la Torre J, García-Salamanca A, Molina-Henares AJ, Herrera MC, Ramos JL, Duque E. Identification of conditionally essential genes for growth of Pseudomonas putida KT2440 on minimal medium through the screening of a genome-wide mutant library. Environ Microbiol. 2010 Jun;12(6):1468–85. doi: 10.1111/j.1462-2920.2010.02166.x. Epub 2010 Feb 11. PMID: 20158506.

46. Liberati NT, Urbach JM, Miyata S, Lee DG, Drenkard E, Wu G, Villanueva J, Wei T, Ausubel FM. An ordered, nonredundant library of Pseudomonas aeruginosa strain PA14 transposon insertion mutants. Proc Natl Acad Sci U S A. 2006 Feb 21;103(8):2833–8.

47. Bradley E. Poulsen, Rui Yang, Anne E. Clatworthy, Tiantian White, Sarah J. Osmulski, Li Li, Cristina Penaranda, Eric S. Lander, Noam Shoresh, Deborah T. Hung, Defining the core essential genome of Pseudomonas aeruginosa, Proceedings of the National Academy of Sciences May 2019, 116 (20) 10072–10080

48. Samuel A. Lee, Larry A. Gallagher, Metawee Thongdee, Benjamin J. Staudinger, Soyeon Lippman, Pradeep K. Singh, Colin Manoil, P. aeruginosa essential genes, Proceedings of the National Academy of Sciences Apr 2015, 112 (16) 5189–5194

49. Velázquez, E., Lorenzo, V. de & Al-Ramahi, Y. Recombination-Independent Genome Editing through CRISPR/Cas9-Enhanced TargeTron Delivery. ACS Synth. Biol. (2019)

50. A. E. Velázquez, Y. Al-Ramahi, J. Tellechea, N. Krasnogor and V. de Lorenzo, Targetron-assisted delivery of exogenous DNA sequences into Pseudomonas putida through CRISPR-aided counterselection. In preparation (2021).

51. I. Borodina, P. Krabben, J. Nielsen, Genome-scale analysis of Streptomyces coelicolor A3(2) metabolism. Genome Res. 3, 820–829 (2005).

52. B. M. Dalgaard Mikkelsen, L. D. Buron, B. Salomonsen, C. E. Olsen, B. Gram Hansen, U. Hasbro Mortensen, B. A. Halkier, Microbial production of indolylglucosinolate through engineering of a multi-gene pathway in a versatile yeast expression platform. Metab. Eng. 14, 104–111 (2012).

53. C. M. M. Jessop-Fabre, T. Jakociunas, V. Stovicek, Z. Dai, M. K. Jensen, J. D. Keasling, I. Borodina, EasyClone-MarkerFree: A vector toolkit for marker-less integration of genes into Saccharomyces cerevisiae via CRISPR-Cas9. Biotechnol. J. 11, 1110–1117 (2016).

54. D. M. Ahmad, M. Hirz, H. Pichler, H. Schwab, Protein expression in Pichia pastoris: Recent achievements and perspectives for heterologous protein production. Appl. Microbiol. Biotechnol. 98, 5301–5317 (2014).

55. E. K. A. Datsenko, B. L. Wanner, J. Beckwith, One-step inactivation of chromosomal genes in Escherichia coli K-12 using PCR products. Proc. Natl. Acad. Sci. U. S. A. 97, 6640– 6645 (2000).

56. F. Y. Li, Z. Lin, C. Huang, Y. Zhang, Z. Wang, Y. Tang, T. Chen, X. Zhao, Metabolic engineering of Escherichia coli using CRISPR–Cas9 meditated genome editing. Metab. Eng. 31, 13–21 (2015).

57. G. K. Labun, T. G. Montague, M. Krause, Y. N. Torres Cleuren, H. Tjeldnes, E. Valen, CHOPCHOP v3: expanding the CRISPR web toolbox beyond genome editing. Nucleic Acids Res. 47, W171–W174 (2019).

58. H. Z. Lin, B. Deng, Z. Jiao, B. Wu, X. Xu, D. Yu, W. Li, A versatile mini-mazF-cassette for marker-free targeted genetic modification in Bacillus subtilis. J. Microbiol. Methods. 95, 207–214 (2013).

59. I. B. M. Koo, G. Kritikos, J. D. Farelli, H. Todor, K. Tong, H. Kimsey, I. Wapinski, M. Galardini, A. Cabal, J. M. Peters, A. B. Hachmann, D. Z. Rudner, K. N. Allen, A. Typas, C. A. Gross, Construction and Analysis of Two Genome-Scale Deletion Libraries for Bacillus subtilis. Cell Syst. 22, 291–305 (2017).

60. J. Altenbuchner, Editing of the Bacillus subtilis Genome by the CRISPR-Cas9 System. Appl. Environ. Microbiol. 82, 5421–7 (2016).

61. K. M. A. Priceid, R. Cruz, S. Baxter, F. Escalettes, S. J. Rosser, CRISPR-Cas9 In Situ engineering of subtilisin E in Bacillus subtilis. PLoS One. 14, e0210121 (2019).

62. Aparicio, T., de Lorenzo, V. & Martínez-García, E. CRISPR/Cas9-Based Counterselection Boosts Recombineering Efficiency in Pseudomonas putida. Biotechnol. J. (2018)

63. Choi, KH., Schweizer, H. mini-Tn7 insertion in bacteria with single attTn7 sites: example Pseudomonas aeruginosa. Nat Protoc 1, 153–161 (2006).

64. Kessler, B., de Lorenzo, V. & Timmis, K. N. A general system to integrate lacZ fusions into the chromosomes of gram-negative eubacteria: regulation of the Pm promoter of the TOL plasmid studied with all controlling elements in monocopy. MGG Mol. Gen. Genet. (1992). doi:10.1007/BF00587591

65. A. R. E. Cobb, Y. Wang, H. Zhao, High-Efficiency Multiplex Genome Editing of Streptomyces Species Using an Engineered CRISPR/Cas System. 18, 3 (2018).

66. B. T. Kieser, M. J. Bibb, M. J. Buttner, K. F. Chater, D. A. Hopwood, Practical Streptomyces Genetics (The John Innes Foundation, 2000).

67. C. F. Fang, K. Salmon, M. W. Y. Shen, K. A. Aeling, E. Ito, B. Irwin, U. P. C. Tran, G. W. Hatfield, N. A. Da Silva, S. Sandmeyer, A vector set for systematic metabolic engineering in Saccharomyces cerevisiae. Yeast (2011), doi:10.1002/yea.1824.

68. D. R. Geertsma, R. Dutzler, A Versatile and Efficient High-Throughput Cloning Tool for Structural Biology. Biochemistry. 50, 3272–3278 (2011).

69. E. Santos-Beneit, Genome sequencing analysis of Streptomyces coelicolor mutants that overcome the phosphate-depending vancomycin lethal effect. BMC Genomics. 19 (2018), doi:10.1186/s12864-018-4838-z.

70. F. K. Peebo, P. Neubauer, Application of Continuous Culture Methods to Recombinant Protein Production in Microorganisms. Microorganisms. 6, 56 (2018).

71. A. P. Masella, A. K. Bartram, J. M. Truszkowski, D. G. Brown, J. D. Neufeld, PANDAseq: PAired-eND Assembler for Illumina sequences. BMC Bioinformatics. 13 (2012), doi:10.1186/1471-2105-13-31.

72. G. Li, R. Durbin, Fast and accurate short read alignment with Burrows-Wheeler transform. Bioinformatics. 25, 1754–1760 (2009).

73. H. D. Li, J.-Z. Song, M.-H. Shan, S.-P. Li, W. Liu, H. Li, J. Zhu, Y. Wang, J. Lin, Z. Xie, A fluorescent tool set for yeast Atg proteins. Autophagy. 11, 954–960 (2015).

74. I. E. DiCarlo, A. Chavez, S. L. Dietz, K. M. Esvelt, G. M. Church, Safeguarding CRISPR-Cas9 gene drives in yeast. Nat. Biotechnol. 33, 1250–1255 (2015).

75. J. D. A. Treco, F. Winston, in Current Protocols in Molecular Biology (2008).

76. K. V. Stovicek, C. Holkenbrink, I. Borodina, CRISPR/Cas system for yeast genome engineering: advances and applications. FEMS Yeast Res. 17, fox030 (2017).

77. L. Y. Tong, P. Charusanti, L. Zhang, T. Weber, S. Y. Lee, CRISPR-Cas9 Based Engineering of Actinomycetal Genomes. ACS Synth. Biol. 4, 1020–1029 (2015).

78. M. Wijsman, M. A. Swiat, W. L. Marques, J. K. Hettinga, M. van den Broek, P. de la Torre Cortés, R. Mans, J. T. Pronk, J.-M. Daran, P. Daran-Lapujade, A toolkit for rapid CRISPR-SpCas9 assisted construction of hexose-transport-deficient Saccharomyces cerevisiae strains. FEMS Yeast Res. 19, foy107 (2019).

79. N. F. J. Ryan, D. Nakada, M. J. Schneider, Is DNA replication a necessary condition for spontaneous mutation? Z Vererbungsl. 91, 38–41 (1961).

80. O. F. J. Ryan, T. Okada, T. Nagata, “Spontaneous Mutation in Spheroplasts of Escherichia coli” (1963).

81. P. L. Loewe, V. Textor, S. Scherer, High Deleterious Genomic Mutation Rate in Stationary Phase of Escherichia coli. Sci. (Washington, DC, U. S.). 302 (2003) (available at http://science.sciencemag.org/).

82. Q. H. J. Bull, G. J. Mckenzie, P. J. Hastings, S. M. Rosenberg, Evidence That Stationary-Phase Hypermutation in the Escherichia coli Chromosome Is Promoted by Recombination. Genetics. 154, 1427–1437 (2000).

83. H.-M. Sung, R. E. Yasbin, Adaptive, or Stationary-Phase, Mutagenesis, a Component of Bacterial Differentiation in Bacillus subtilis. J. Bacteriol. 184, 5641–5653 (2002).

84. L. Kasak, R. Hõrak, M. Kivisaar, Promoter-creating mutations in Pseudomonas putida: A model system for the study of mutation in starving bacteria. Proc. Natl. Acad. Sci. U. S. A. 94, 3134–3139 (1997).

85. D. F. Steele, S. Jinks-Robertson, An examination of adaptive reversion in Saccharomyces cerevisiae. Genetics. 132, 9–21 (1992).

86. A. Achilli, N. Matmati, E. Casalone, G. Morpurgo, A. Lucaccioni, Y. I. Pavlov, N. Babudri, The exceptionally high rate of spontaneous mutations in the polymerase delta proofreading exonuclease-deficient Saccharomyces cerevisiae strain starved for adenine. BMC Genet. 5 (2004), doi:10.1186/1471-2156-5-34.

87. B. Noble, J. Min, J. Olejarz, J. Buchthal, A. Chavez, A. L. Smidler, E. A. DeBenedictis, G. M. Church, M. A. Nowak, K. M. Esvelt, Daisy-chain gene drives for the alteration of local populations. Proc. Natl. Acad. Sci. 116, 8275–8282 (2019).

88. N. Windbichler, M. Menichelli, P. A. Papathanos, S. B. Thyme, H. Li, U. Y. Ulge, B. T. Hovde, D. Baker, R. J. Monnat, A. Burt, A. Crisanti, A synthetic homing endonuclease-based gene drive system in the human malaria mosquito. Nature (2011), doi:10.1038/nature09937.

